# Cooperative multivalency converts disorder into rods, resolving a paradox in cellular architecture

**DOI:** 10.1101/2025.07.16.665182

**Authors:** Douglas R. Walker, Aidan Estelle, York-Christoph Ammon, Yujuan Song, Qianru H. Lv, Patrick N. Reardon, Muneyoshi Ichikawa, Anna Akhmanova, Elisar J. Barbar

## Abstract

The cortically anchored adaptor KANK1 organizes microtubules at focal adhesions through a long, intrinsically disordered linker (L2), yet how this linker spans the ∼35–50 nm membrane–microtubule gap is unclear. Here, we combine in-cell, biochemical, and biophysical assays, predictions of motif interaction and multivalent assembly using AlphaFold, and structural analysis by electron microscopy to show that the hub protein LC8, which binds more than 100 clients, converts the intrinsically disordered 600 amino acid L2 into an elongated, multivalent, rod-like assembly. In contrast, isolated motif peptides fail to bind LC8 at physiologically relevant concentrations, indicating that strong complex formation arises from cooperativity among multiple weak sites. These results establish LC8 as a molecular switch that rigidifies and extends KANK1 L2 via distributed weak motifs and short linkers. This interaction produces compositionally homogeneous yet conformationally adaptable rods, long enough to bridge the membrane–microtubule gap, resolving the paradox. This work expands the LC8 binding repertoire, reveals design principles for multivalent assembly, and suggests a generalizable strategy for tuning length, rigidity, and flexibility in large protein architectures.

## Introduction

Intrinsic disorder is increasingly recognized as a central structural factor and regulator of large protein assemblies with examples in cellular transport and sensing(*1–3*), muscular elasticity(*4*), DNA transcription and storage(*5–7*), and protein degradation(*8*). In the dynein-dynactin transport complex, disordered regions are integral parts of many adaptor proteins(*9*, *10*) and of the complex itself(*1*, *11*, *12*), where they contribute to assembly and regulation of autoinhibition. In the nuclear pore complex (NPC), over one third of proteins belong to the FG family, characterized by large disordered swathes (200-700 aa) of phenylalanine-glycine repeats that mediate selective transport of karyopherin-cargo complexes(*2*, *13*, *14*). Adaptor proteins involved in clathrin-mediated endocytosis assemblies also harbor large intrinsically disordered regions (IDRs) that promote membrane curvature(*15*) and contain numerous short linear motifs (SLiMs). Importantly, mutations within IDRs have been linked to diverse disease states(*16*), reinforcing findings that disorder can encode precise functional constraints through sequence-dependent interactions(*17*, *18*). Here we explore how intrinsic disorder orchestrates the bridging of two cytoskeletal regulatory complexes with the cell cortex.

At the cell cortex, focal adhesions (FAs) and cortical microtubule stabilizing complexes (CMSCs) serve as key hubs, linking the actin and microtubule cytoskeletons. FAs contain more than 50 distinct proteins(*19*) that form highly organized plaques at the plasma membrane(*20*), while CMSCs are composed of a small set of proteins which together are responsible for cortical microtubule targeting and regulation of FA turnover(*21*). Bridging these two large complexes is the tumor suppressor protein KANK1(Fig. 1a)(*22*, *23*). KN motif and ankyrin repeat protein 1 (KANK1) is a ∼150 kDa member of the KANK protein family (KANK1-4)(*23*, *24*), characterized by a KANK N-terminal (KN) domain and C-terminal ankyrin repeat (Ank) domain(*23–25*). KANK1 contains predicted coiled-coil (CC) domains within residues 258-499 and two long disordered linkers, L1 and L2, spanning residues 60-258 and 500-1080 (Fig. 1b,c). Notably, L2 stands out as an IDR due to its remarkable length of 580 amino acids. Both the UniProt-AlphaFold model of KANK1(*26*) and JPRED(*27*) analysis confirm that L2 is largely disordered with only a short, low-confidence coil near residue 825.

**Fig. 1.**
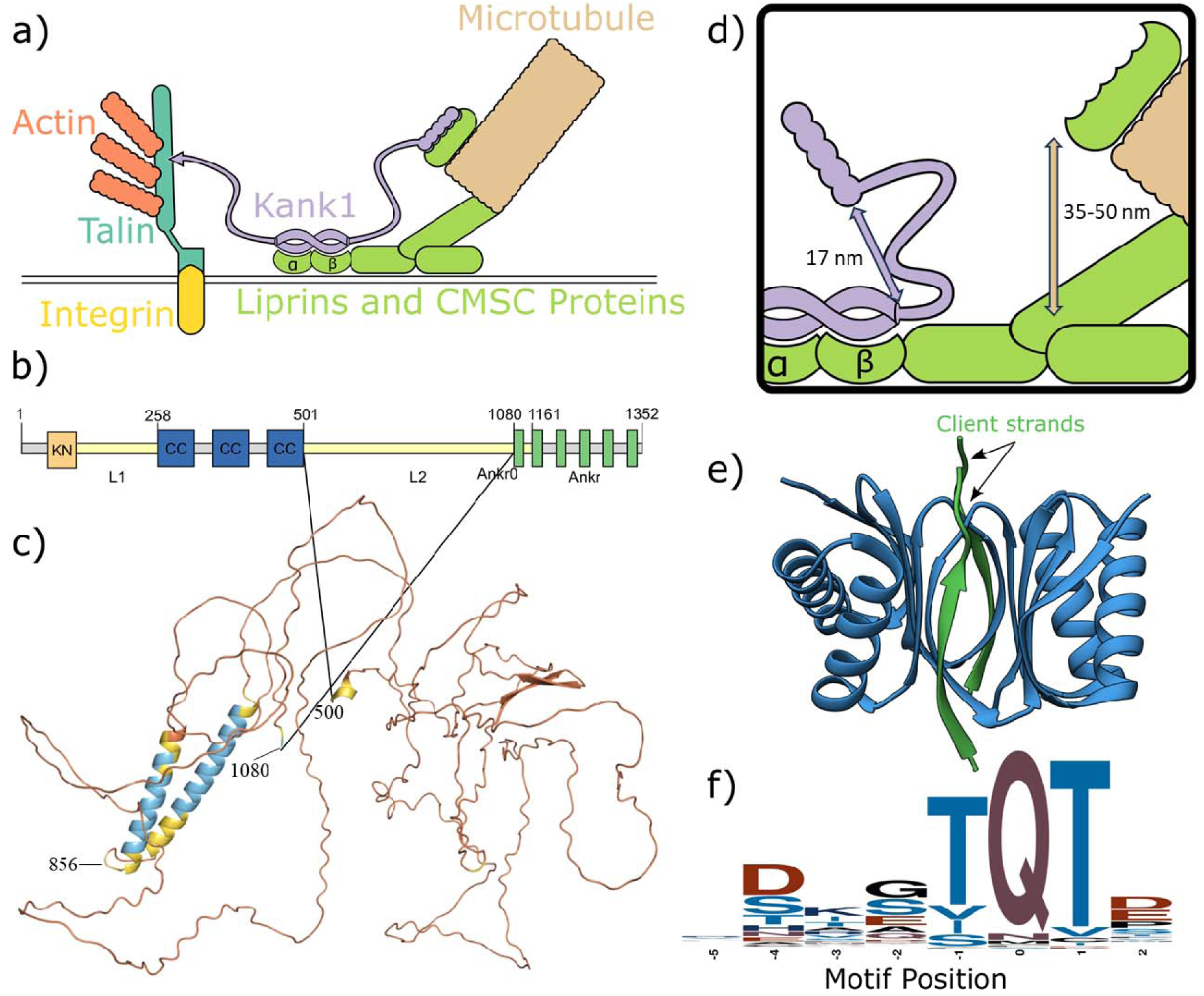
KANK1 and LC8 structure and function. (a) Schematic of KANK1 (purple) binding to FAs (left shown in teal, orange, and yellow) and CMSCs (right shown in green and tan). KANK1 interacts with Talin1 (teal) to bind FAs and is additionally recruited to CMSCs by Liprins (light green) through interactions with the ankyrin and coiled-coil domains. (b) Diagram outlining KANK1 structure, including KN domain (orange), ankryin repeat domain (Ankr, green), coiled-coils (CC, blue) and linker sequences (L1, L2, yellow). (c) AlphaFold structure as deposited on UniProt(*26*) of KANK1_500-1080_, colored by confidence. Most of the chain is predicted disordered, but a lower confidence coil can also be seen. (d) Zoom-in of model similar to (a), into L2 and the microtubule region, emphasizing the mismatch between the L2 end-to-end distance and the separation between the cell membrane and microtubules. (e) Ribbon diagram of LC8, drawn in blue, with two client strands (PDB 2P2T). Clients (green) take a beta-strand structure when bound to LC8. (f) Sequence logo of the 8 amino acid LC8 motif, constructed from all known LC8-binding sequences in the LC8Hub database(*37*).

L2 links the CC domain, which binds Liprin-β1, a component of the CMSC (*28*), to the Ank domain which recruits the kinesin-4 KIF21A to CMSCs(*24*, *25*, *28–31*). KIF21A inhibits cortical microtubule growth(*28*) and localizes to microtubules which terminate 35-50 nm from the cell membrane(*32*). Consequently, the 580-residue L2 must span this substantial gap to enable interaction between Ank domain and KIF21A to connect microtubules to the cell membrane. To assess whether L2 alone can bridge this gap, we first computed its end-to-end distance using the worm-like chain model(*33*) which accounts for polymer stiffness and predicts end-to-end distance distributions in disordered protein chains(*34*). Applying this model to L2, as outlined in the methods, yields a rms end-to-end distance of ∼17 nm (Fig. 1d). Similarly, ALBATROSS, a sequence-dependent predictor of IDR ensemble properties(*35*), estimates an average end-to-end distance of 15-20 nm. These values fall far short of the 35-50 nm gap. To evaluate the probability of extreme extensions, we calculated the fraction of the ensembles that are stretched far enough to span the gap. According to the worm-like chain model, 0.41% of the ensemble of structures extend further than 35 nm while 0.0004% exceed 50 nm(*36*). ALBATROSS predicts a higher ensemble fraction (4% and 0.008%, respectively), and yet still suggests that L2 is not long enough to efficiently span the gap between the cell membrane and microtubule ends to allow the Ank domain to interact with KIF21A, especially when including an alpha helical structure which will shorten the effective length. If L2 cannot bridge the required gap on its own, then there should be an additional mechanism to extend L2 and stabilize its reach.

A promising candidate for interacting with KANK1’s L2 linker is LC8. LC8 has been observed to form highly concentrated puncta near the cell cortex, in proximity to FAs(*37*) suggesting a potential role in FA-associated complexes, possibly through interaction with KANK1. LC8, a small homodimer hub protein (20 kDa dimer mass), binds 100+ clients at a short linear motif (SLiM) found in IDRs(*37*, *38*). Hub proteins like LC8 interact with many binding partners and serve as central nodes in protein-protein networks(*39*, *40*) and thus control diverse regulatory mechanisms. The LC8 recognition motif is characterized by a threonine-glutamine-threonine (TQT) amino acid triad which anchors clients within the binding site (Fig. 1e,f)(*37*, *41*). LC8 has two rotationally symmetric binding grooves which accommodate two client strands(*42*) and promote client dimerization when bound to two copies of the same protein (Fig. 1e)(*38*, *43–47*). LC8 partners are involved in a host of cellular processes, including at the nuclear pore(*48*, *49*), transcriptional regulation(*50–52*), neuronal synaptic organization(*46*, *53*), and cytoskeleton dynamics during cell division(*41*), among others(*54–56*). Multivalent LC8 partners assemble as beads-on-a-string, stringing two intrinsically disordered client strands through LC8 “beads”(*50*, *57*), thereby forming polybivalent complexes with variable rigidity. These multivalent LC8-client assemblies exhibit substantial heterogeneity due to their dynamic nature, variation in LC8 occupancy, and variable cooperativity among multiple motifs(*50*, *58–60*).

Although it is challenging to characterize biophysically, understanding this structural and conformational diversity is essential for elucidating LC8’s functions.

We developed two complementary computational methods for predicting LC8 binding partners. The first, LC8Pred, utilizes amino acid propensities within SLiMs of known LC8 binding sites and predicts at least 374 potential motifs in the human proteome(*37*). The second, AlphaFold-Pred, uses AlphaFold structural predictions of LC8 bound to short peptides parsed from the protein of interest to identify potential LC8 binding sites(*61*).

While LC8Pred is designed specifically for LC8 and is useful for predicting canonical motifs, AlphaFold-Pred can identify binders with SLiM sequences that diverge from known binders. Together these methods provide complementary strategy for interrogating protein sequences for their potential for binding LC8.

Here, we demonstrate that KANK1-L2 binds LC8 in HeLa cells, where KANK1 localizes with LC8, recruiting it to FA edges. In vitro, we establish that although L2 contains only one canonical LC8 motif, multiple weak sites are required to facilitate multivalent binding, resulting in a cooperativity-driven ladder-like assembly. We identify these binding sites through a combination of isolated peptide binding, LC8Pred, and AlphaFold predictions. Remarkably, the LC8/KANK1 complex is compositionally homogenous, which is unusual among multivalent LC8 assemblies. Negative stain electron microscopy (NSEM) reveals a hinged rod-like structure with conformational heterogeneity around the hinge. Both NSEM and multivalent AlphaFold predictions indicate that LC8 binding elongates L2 to longer than the ∼50 nm gap between the cell membrane and microtubule ends. These results uncover a new structural role for intrinsic disorder and LC8-meciated interactions in bridging large cytoskeletal complexes.

## Results

### KANK1 colocalizes with LC8 around focal adhesions

Pulldown mass spectrometry provided the first evidence of an interaction between KANK1 and LC8, with LC8 co-purifying along with known KANK1 partners Talin1, Liprin-β1, and KIF21A (Fig. S1)(35). To validate this interaction is specific to KANK1, HeLa cells stably expressing LC8 tagged with GFP were stained for Paxillin (a FA marker), and KANK1 or KANK2 (Fig. 2a). LC8 signal strongly overlaps with KANK1 in patches at FA edges whereas LC8 and KANK2 do not co-localize, suggesting that the KANK1/LC8 interaction may be unique in the KANK family. Depletion of KANK1 via siRNA-mediated knockdowns eliminates localization of LC8 to FAs, confirming KANK1 as responsible for LC8’s presence near FAs. In contrast, knockdown of KANK2 does not impact LC8 localization, further confirming that KANK2 does not interact with LC8.

**Fig. 2.**
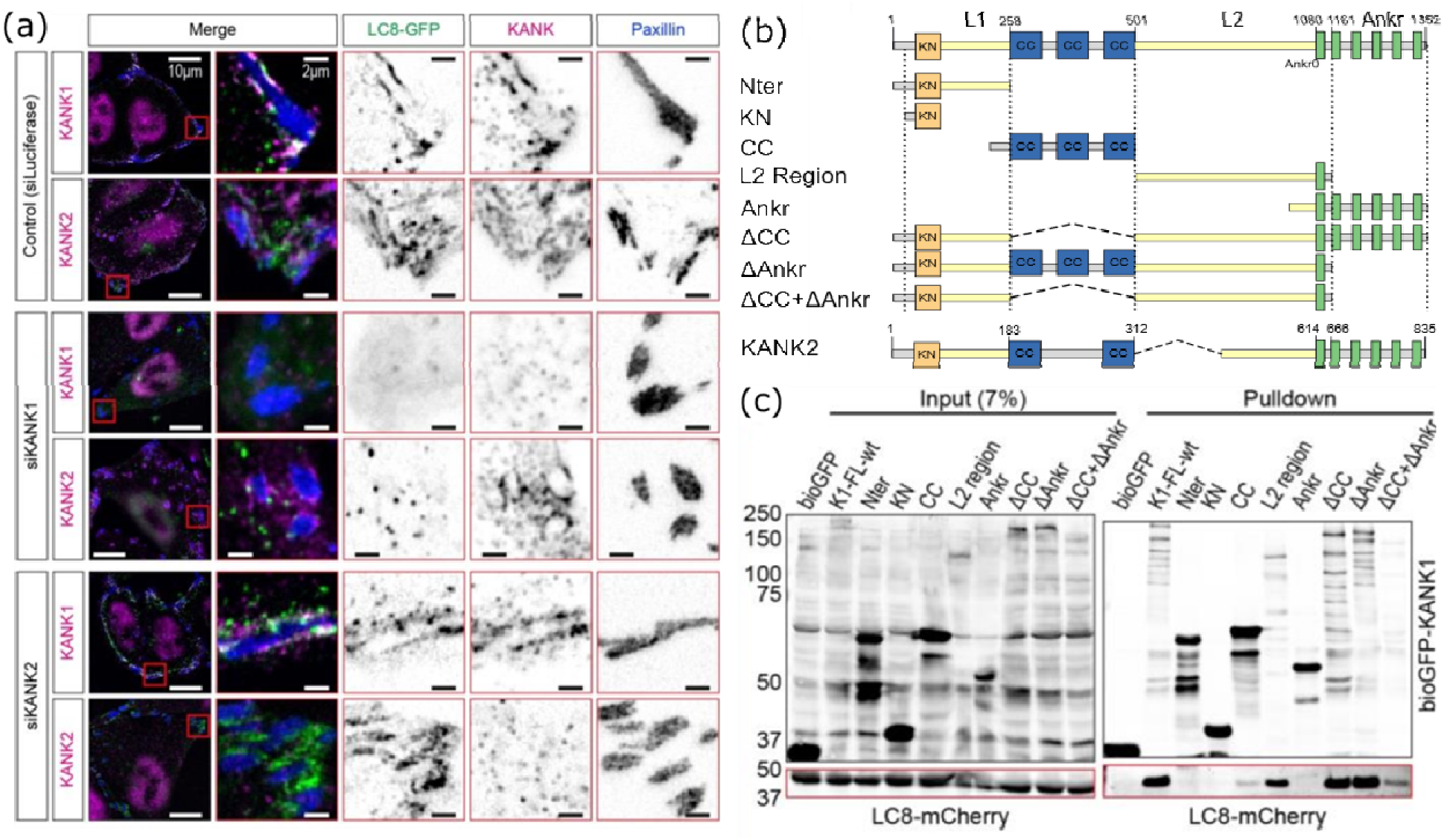
LC8 Colocalizes in KANK1 in HeLa cells. (a) Cells expressing LC8-GFP (green), stained for Paxillin (blue) and KANK1 or KANK2 (red). LC8 and KANK1 colocalize at the edges of focal adhesions. Cells were transfected with siRNA for luciferase (as a control), siKANK1 and siKANK2. Silencing of KANK1 abolishes LC8 colocalization. (b) Diagram of KANK1 structure, with bio-GFP tagged constructs designed for pulldowns shown and KANK2 for comparison. (c) Streptavidin pulldowns of LC8-mCherry with bioGFP-KANK1. Left is input of crude cell lysate pulled down, while right is blotted for GFP (top) and mCherry (bottom).

Structurally, KANK1 and KANK2 differ slightly in the L1 and CC domains, but their most striking difference is in the L2 length: 353 residues in KANK2, compared to 580 residues in KANK1.

### KANK1’s intrinsically disordered L2 binds LC8

To identify the LC8-binding region in KANK1, we performed streptavidin-based pulldown assays, using bioGFP-KANK1 constructs as bait. In these experiments Ankr0 was included in L2 because it is not essential for binding to KIF21A. These experiments revealed that LC8 binds L2 (residues 501 – 1161, Fig. 2b,c). L2 strongly pulls down LC8 on its own, and all constructs that pull down LC8 contain L2. Importantly, no other isolated domain of KANK1 pulls down LC8.

To refine the binding region, we tested multiple fragments of L2 in pulldown assays (Fig. 3a,b). Fragments including 501-750, 501-650, and 650-750 strongly pull down LC8, whereas 750-918 and 750-1161 show weak pulldown, and 800-1161 does not pull down LC8. These results implicate residues 501-750 as the primary LC8 binding domain with strong binding in 501-650 and 650-750, and weaker binding in 750-800.

**Fig. 3.**
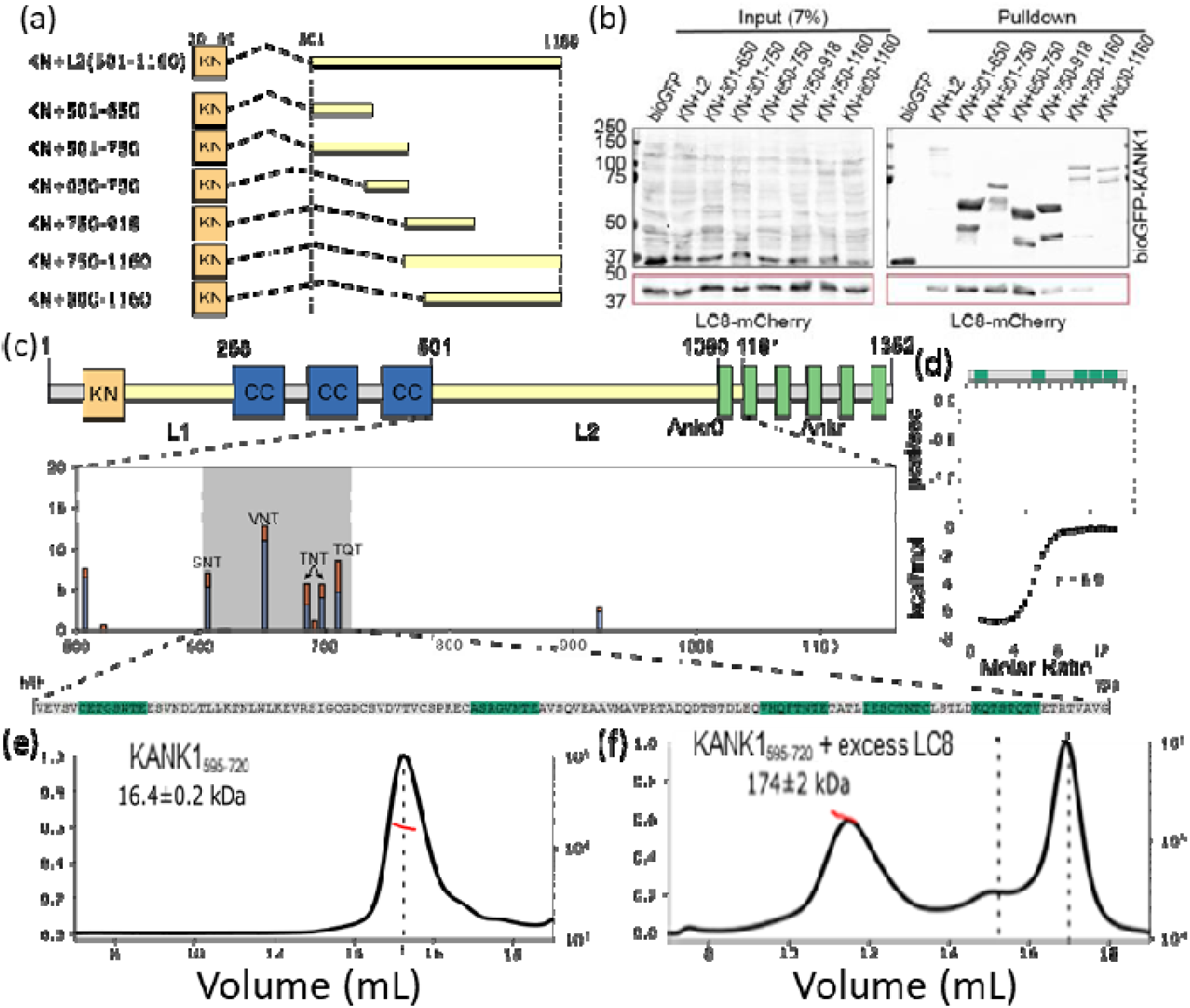
KANK1_595-720_ binds LC8 multivalently. (a) Constructs of KANK1 including the KN domain and sections of L2 to narrow down the binding region for LC8. (b) Pulldown with KN-L2 constructs indicating that LC8 binding occurs between residue 500 and 750. (c) Domain structure of KANK1 showing folded domains (orange, blue, green) and regions of disorder (yellow) (top). Beneath, the LC8pred scores for 500-1160 (L2 + Ankr0) are shown (middle). Due to the density of sequences that score well in LC8pred within residues 595-720, we designed and express constructs of this region (bottom). (d) Isothermal titration calorimetry of binding between LC8 and KANK1_595-720_. (e,f) SEC-MALS for (e) free KANK1_595-720_ and (f) KANK1_595-720_ with excess LC8 to investigate the mass of the fully bound complex.

We next utilized LC8Pred(*37*) to predict LC8 binding motifs within L2. Although no sites surpass out stringent thresholds for reducing false positives, within residues 600-720 five motifs with low prediction scores were identified (Fig. 3c). The strongest predicted site has a sequence of ASRGVNTE with N as anchor instead of Q at residue 651. The other four are much weaker: a SNT motif at position 605, two TNT motifs at 685 and 697, and a TQT motif at 710. Despite low prediction scores, the density of predicted sites warrants further investigation.

### KANK1 binds LC8 multivalently

To investigate KANK1/LC8 interactions, we recombinantly expressed a construct corresponding to residues 595-720 of L2, which includes all predicted LC8-binding motifs and preformed isothermal titration calorimetry (ITC) experiments, titrating LC8 into KANK1_595-720_ (Fig. 3d). The ITC fit resulted in a binding affinity of 893 ± 49 nM, and a - 6.96 ± 0.03 kcal/mol change in enthalpy, consistent with a tight-binding complex. Notably, the fitted stoichiometry was 5.78 ± 0.02 indicating a high LC8 binding ratio. While the *n* value is not always reliable due to poor precision and accuracy of concentration measurements and assumptions of the independent-sites model(*62*) its high value is an indicator that KANK1 binds LC8 at multiple motifs. KANK1_595-720_ binding to several LC8 dimers, despite none of the predicted motifs exceeding the LC8Pred threshold, indicates that multivalency is critical for the strong binding observed by ITC.

To further assess stoichiometry, we performed size exclusion chromatography multiangle light scattering (SEC-MALS) on KANK1_595-720_. In isolation, the protein elutes as a single peak (Fig. 3e) with a MALS-derived mass of 16.4 ± 0.2 kDa, matching the expectation for the monomeric mass. When mixed with excess LC8 and purified by SEC, a dominant new peak corresponding to the complex appears (Fig. 3f), with a MALS-derived mass of 174 kDa, closely matching the mass expected for a 14:2 LC8:KANK1_595-720_ complex (181 kDa). The mass along this peak is not uniform due in part to partial dissociation during SEC, which produces peaks for dimeric LC8 and monomeric KANK1_595-720_. Nevertheless, the observed complex mass corroborates the high stoichiometry from ITC and supports a 14:2 complex, where seven LC8 dimers bridge two strands of KANK1_595-720_.

### KANK1-LC8 binding is highly cooperative

To investigate details of LC8-client binding and assess the affinity of the five predicted LC8-binding motifs individually, we synthesized peptides corresponding to each motif and measured their binding to LC8 by ITC. None of the peptides exhibited measurable binding affinity (Fig. 4a). The isotherm for peptide 1 showed no evidence of binding while peptides 2 to 5 showed only very weak binding. All isotherms were collected at an LC8 concentration of 60 µM, indicating that the Kd for LC8 and peptides binding is well above 60 µM in all cases. These results align with LC8Pred predictions that none of these sites are expected to bind with high affinity. The stark contrast between the negligible binding of isolated peptides and the tight binding observed for KANK1_595-720_ suggests that LC8-KANK1 complex formation is strongly driven by cooperativity among multiple LC8 motifs within KANK1.

**Fig. 4.**
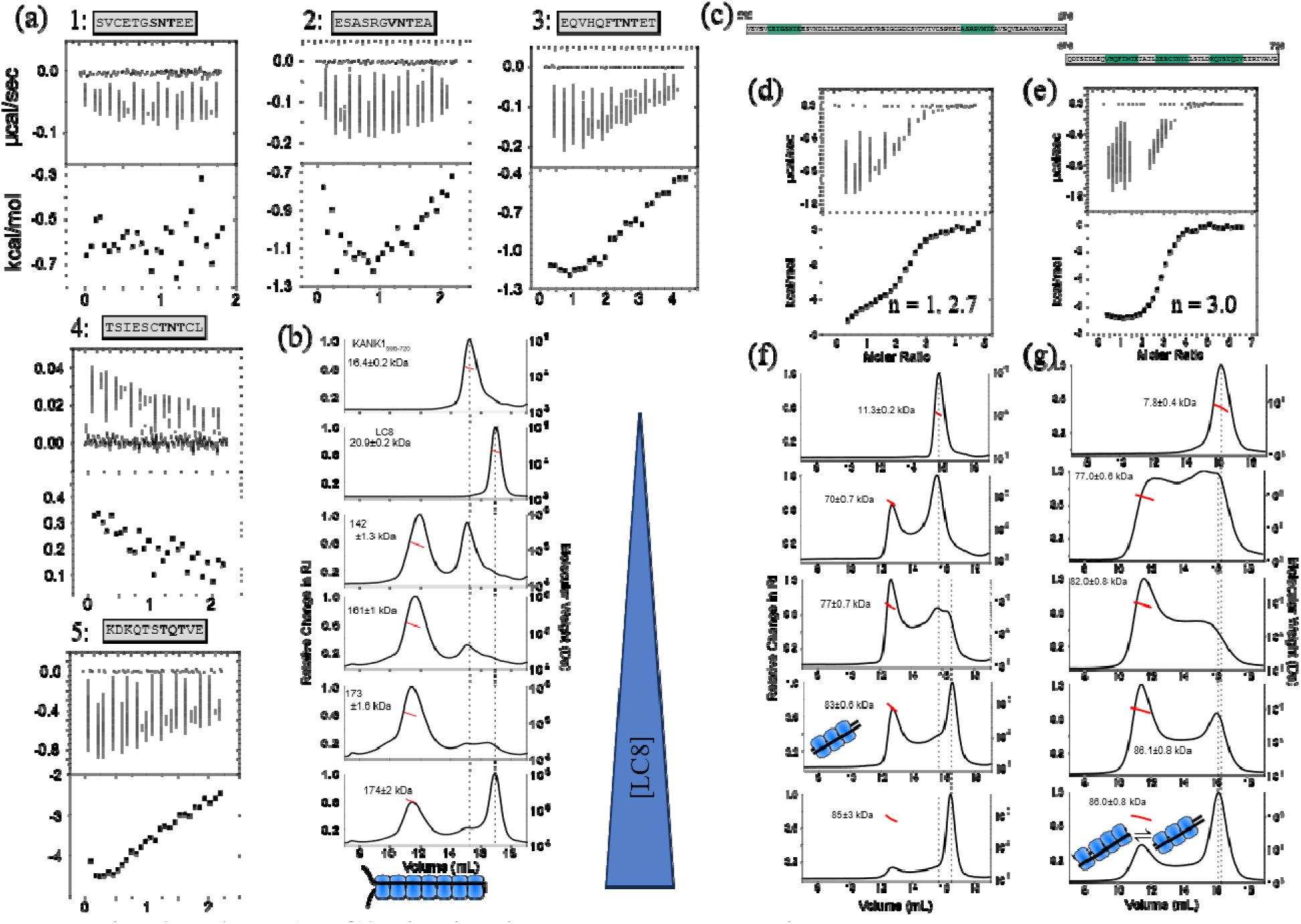
KANK1-LC8 binding is strongly cooperative. (a) ITC of binding between LC8 and KANK1_595-720_ peptides. (b) MALS of KANK1_595-720_, LC8, and their titration. LC8 concentration increases as you go down the panel as illustrated by the blue triangle. (c) Short KANK1 constructs for locating binding sites. (d,e) ITC titrations of LC8 into KANK1_595-670_ (d) and KANK1_670-720_ (e). Fits are shown for a one-set-of-sites model. (f,g) MALS titration of 50 uM KANK1_595-670_ (f) and KANK1_670-720_ with LC8 (g). Top panel shows MALS trace of KANK1 construct alone. Saturated complex masses correspond to 2:6 and 2:7 stoichiometries, respectively. LC8 concentration increases down the panel as signified by the blue triangle. For all SEC-MALS, RI is normalized to the highest intensity. Red lines indicate measured masses for complex peaks, with the average mass listed in each panel. Dotted lines mark the elution times of each protein alone. Models in blue illustrate complexes consistent with the largest masses observed in the titrations.

Sedimentation velocity analytical ultracentrifugation (SV-AUC; Fig. S2) further supports the strong cooperativity conclusion. At low LC8:KANK1_595-720_ ratios, three AUC species are evident corresponding to free KANK1_595-720_, an intermediate size complex at ∼3 svedbergs (S), and a large complex at ∼5.5 S. As the titration progresses, the minor intermediate species and the KANK_595-720_ peaks both disappear. Specifically, a shift away from excess KANK_595-720_ and towards a new peak for excess LC8 occurs at higher titration points, suggesting a saturated complex. At higher titration points, the bound state peak shifts from 6 S to 7 S. While this could reflect a distinct structural state at these higher titrations, it is more likely that the shift in latter titration points is due to a reduction in exchange rate between partially and fully bound states and an equilibrium shift toward the fully bound state, consistent with the simple model used for analysis(*63*, *64*).

Finally, SEC-MALS titration corroborates strong cooperativity (Fig. 4b). Even at low LC8:KANK1_595-720_ ratios, chromatograms exhibit high mass complexes while a separate large peak corresponding to excess KANK1_595-720_ is still evident. Before a peak for excess LC8 is observed, the complex reaches its maximum mass corresponding to a 14:2 complex. Dynamic exchange is once again apparent in this titration through the raised baseline between complex peaks and free protein peaks and by the range of masses measured by MALS for each peak. This is consistent with the exchange observed by AUC. However, under the diluting conditions of SEC and a 1 µM starting concentration for the complex, this dynamic exchange is minimal, reinforcing the conclusion that binding is highly cooperative and stabilizes a strongly bound, fully occupied complex.

### Shorter KANK1 constructs indicate the locations of the additional LC8 binding sites

To locate the LC8 binding sites missed by LC8Pred, we generated two pared constructs, dividing KANK1_595-720_ into KANK1_595-670_ and KANK1_670-720_ (Fig. 4c). ITC analysis of KANK1_595-670_ (Fig. 4d) exhibits less cooperativity than KANK1_595-720_, with two transitions at *n*=1 and n=2.7, suggesting one site that binds independently and two others that bind cooperatively. In contrast, the KANK1_670-720_ isotherm (Fig. 4e) depicts a cooperative transition, like with KANK1_595-720_, with n=3.

SEC-MALS titrations provide further insight. KANK1_595-670_ initially forms a complex with mass corresponding to a mixture of 4:2 and 6:2 species (LC8:KANK1, Fig. 4f). As the titration progresses, the complex does not grow beyond the 6:2 ratio, indicating 3 LC8 binding sites within this fragment. KANK1_670-720_ initially exhibits a 6:2 complex (Fig. 4g) but later shows a mixture of 6:2 and 8:2 species. The 8:2 species never dominates, even at large excess of LC8, suggesting the presence of four binding sites, with one too weak to remain bound during SEC. Both truncated constructs exhibit pronounced peak tailing on SEC, indicative of dynamic exchange and weaker complexes compared to KANK1_595-720_. Consistently, ITC measurements of these shorter constructs report weaker binding than seen for the full-length fragment.

### AlphaFold predictions identify 15+ LC8 binding sites

To complement LC8Pred predictions, we applied AlphaFold using our recently developed method(*61*)(Fig. 5a-d). We parsed KANK1-L2 into 16 amino acid long peptides with 8 amino acids overlap between successive sections and used AlphaFold to predict interactions of LC8 dimers with each of the parsed peptides. In developing this approach, we established two evaluation thresholds: an exclusive threshold, optimized for a low false positive rate and best suited for screening new LC8-binding proteins, and an inclusive threshold, optimized for a low false negative rate and best suited for detecting weak or atypical binding sites in proteins already known to bind LC8.

**Fig. 5.**
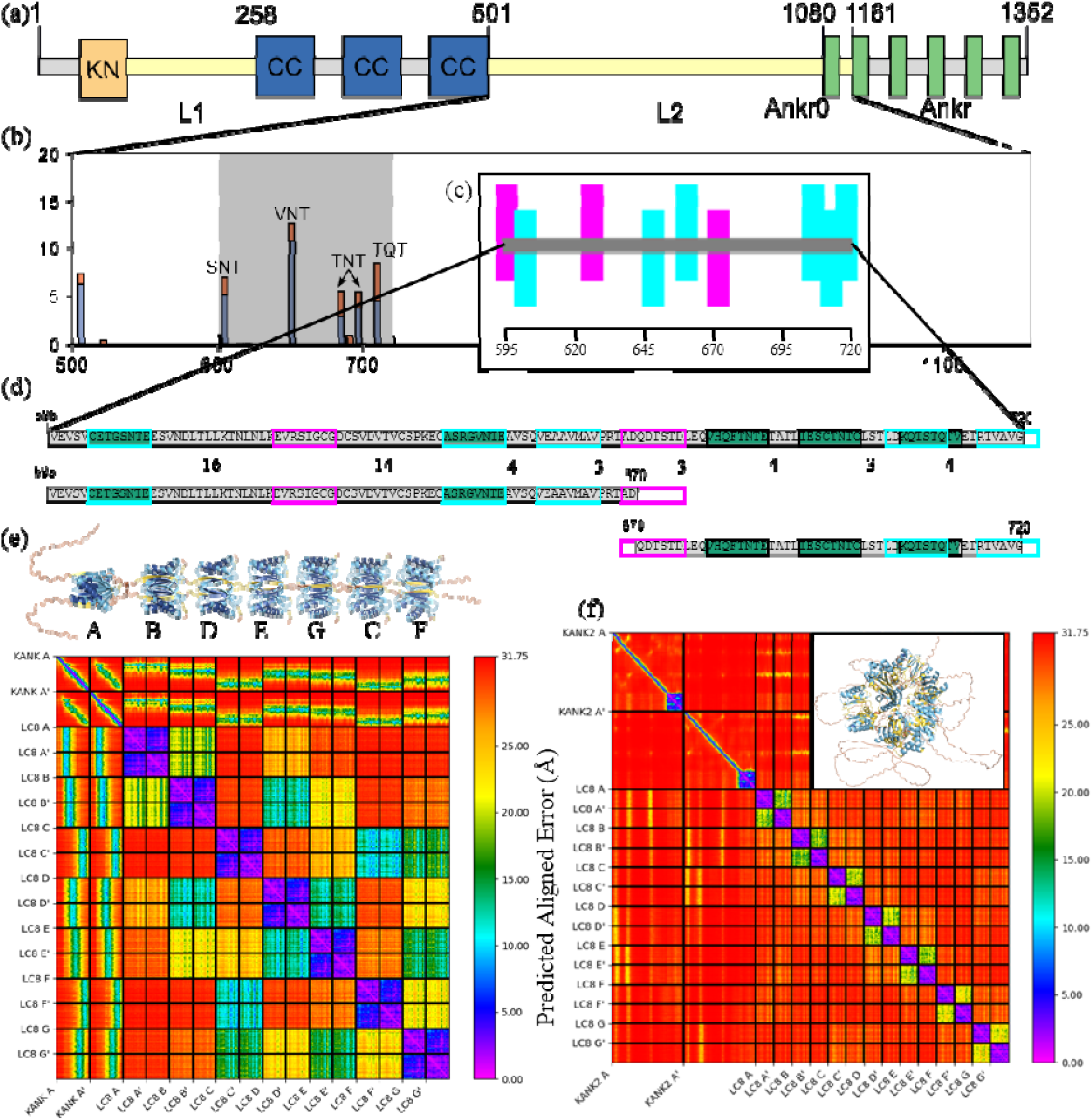
AlphaFold finds remaining LC8 binding sites. (a) Domain structure of KANK1 showing folded domains (orange, blue, green) and regions of disorder (yellow). (b) LC8pred scores for 500-800 are shown. (c) AlphaFold predictions for 595-720, the region spanned by experimental constructs, find the missing binding sites. Cyan results are from the exclusive threshold and magenta from the inclusive. (d) Construct sequences with predicted sites boxed by prediction method: LC8Pred, black; AF exclusive, cyan; AF inclusive, magenta. Overlapped AlphaFold predictions are not marked, for clarity. (e) AlphaFold prediction of KANK1_595-720_ with 7 LC8 dimers and the associated PAE heatmaps. The structure is colored according to AlphaFold pLDDT confidences and LC8s are lettered according to their position in the AlphaFold output and in the heatmap. Heatmap colors indicate highest confidence in violet and lowest confidence in red. In both, KANK1 residues bound to LC8s are much more confident than the surrounding sequence. (f) PAE heatmap for AlphaFold prediction of KANK2-L2 with 7 LC8 dimers and (inset) the associate structure, showing low confidence globally.

Using the exclusive threshold, we identified five binding sites between residues 595 and 722, including a motif which is partially truncated in KANK1_595-720_ and KANK1_670-720_ (Fig. 5c,d). Interestingly, AlphaFold-Pred did not predict the canonical TQT motif, but rather predicted a TST-anchored motif located two residues N-terminal to the TQT. It is noteworthy that this TQT motif is not evolutionarily conserved while the TST motif is conserved across animalia KANK1 sequences as either TST or TNT(*65*), giving more credibility to the AlphaFold prediction. Applying the inclusive threshold revealed two additional motifs: an IGC motif anchored at residue 629 and a TST motif anchored at 674.

SEC-MALS experiments indicate seven binding motifs within KANK1_595-720_, however, combining LC8Pred and AlphaFold predictions yields nine predicted LC8 binding sites. ITC analysis of the peptide SVCETGSNTEE (Fig. 4a) to investigate the motif anchored at 605 exhibited no binding. Given its distal position relative to the other motifs, this site likely does not bind in KANK1_595-720_, though it may bind in the context of the full-length protein. Similarly, the predicted VGE site anchored at residue 720 is partially truncated in KANK1_595-720_ and the missing residues appear essential for binding. Thus, of the nine predicted sites, two likely do not bind, matching experimental observations.

The proximity of the remaining seven predicted sites, separated by short linkers, explains the high cooperativity observed despite their individually weak affinities. In the truncated constructs, KANK1_595-670_ binds three LC8 dimers, consistent with AlphaFold predictions of the IGC, VNT, and VMA anchored binding sites. KANK1_670-720_ exhibits a more complex stoichiometry, forming a mixture of 1:3 and 1:4 complexes. However, the predicted motif anchored at residue 674 is truncated in this construct, missing two N-terminal residues (Fig. 5d). This truncation likely weakens LC8 binding to this motif, consistent with our experimental observations.

AlphaFold Multimer titrations (see Methods) with KANK1_595-720_ (Figs. 5e, S3) predict a highly confident rod-like structure consisting of seven LC8 “beads” packed tightly on two parallel KANK1 “strings”. In contrast, titrations with the 353-residue L2 region of KANK2 with one to seven LC8 dimers showed no evidence of interaction (Figs. 5f, S4); instead LC8 dimers packed in a lattice-like configuration wrapped around the KANK2 strands. This outcome is consistent with our experimental observation that KANK2 does not bind LC8 (Fig. 2a). Importantly, AlphaFold titrations of shorter KANK2-L2 subconstructs (∼120 amino acids in length, to match the length of KANK1_595-720_) similarly do not predict binding (Figs. S5-7).

AlphaFold-Pred(*61*) was applied to predict LC8 binding along the entire L2 region (Fig. 6a). Beyond residue 800, predicted motifs are sparse and weak, consistent with pull-down assays which show no LC8 binding in that region (Fig. 3b). In contrast, residues 500-600 display many weakly predicted motifs separated by short linkers, as seen in KANK1_595-720_. To investigate multivalency in this region, we ran AlphaFold titrations on KANK1_500-615_ and KANK1_541-670_ (Fig. 6b; Figs. S8,9). Titrations with KANK1_500-615_ indicate an optimal stoichiometry of 1:8 KANK:LC8, including an LC8 bound to the LC8Pred predicted motif anchored at residue 605. We argue above that this site does not bind in KANK1_595-720_ but may bind in the context of the full L2. Similarly, KANK1_541-670_ binds eight LC8 dimers optimally, with two binding regions divided by a 16-residue gap containing the nuclear export domain.

**Fig. 6.**
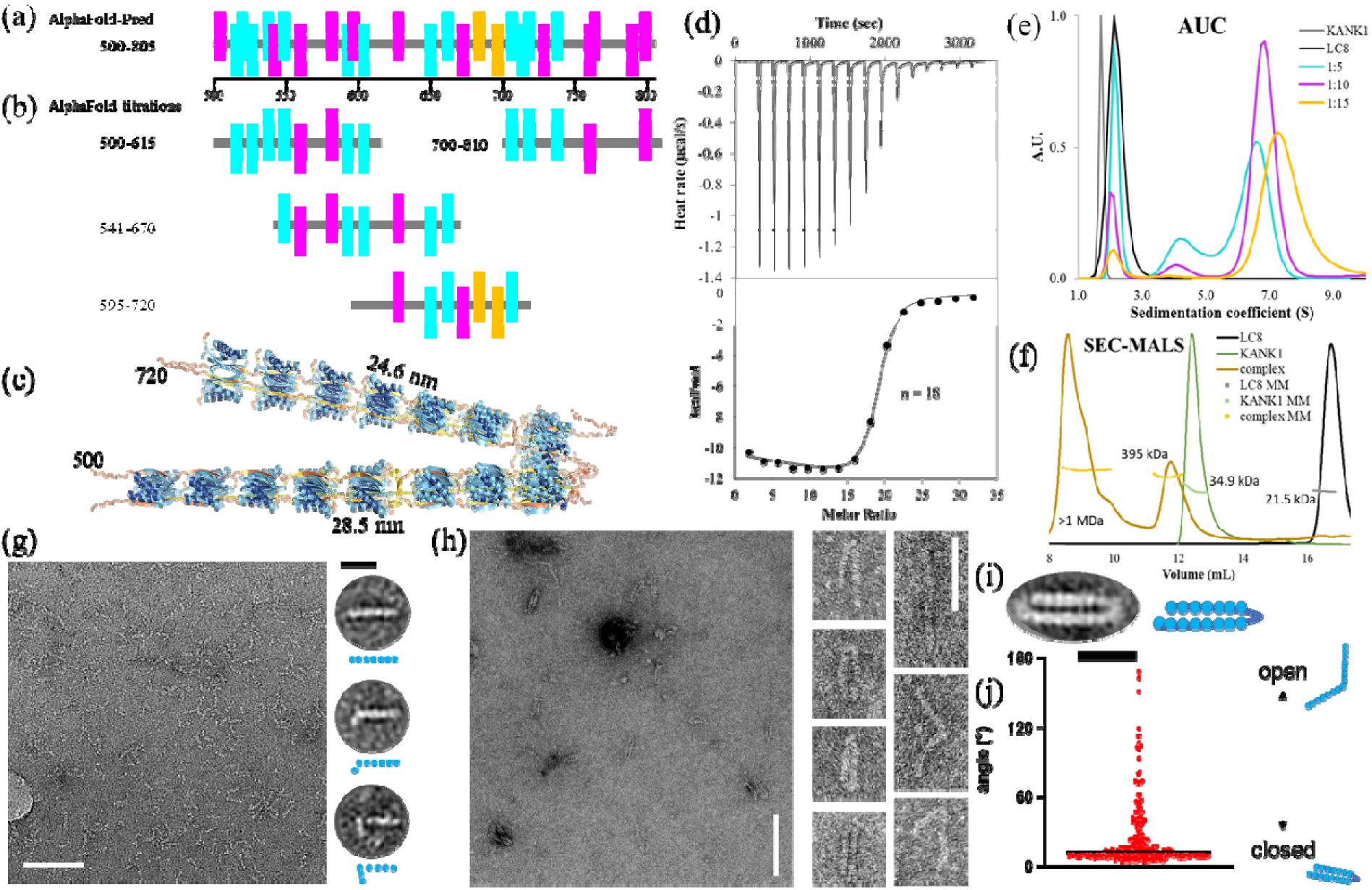
KANK1_500-805_ binds 15 LC8 dimers. (a) AlphaFold predictions of LC8 binding across residues 500-805 of L2. Cyan and magenta correspond to predictions by the exclusive and inclusive thresholds respectively. (b) Locations of bound LC8 dimers in AlphaFold titrations of KANK1_595-720_, KANK1_500-615_, KANK1_541-670_, and KANK1_700-810_ aligned with (a) for comparison. (c) PyMOL aligned structures of predictions of KANK1_595-720_, KANK1_500-615_, and KANK1_541-670_ with 7, 8, and 8 LC8 dimers, respectively, showing the potential of 15 simultaneous LC8 binding events forming a rod with a hinge at the nuclear export domain. (d) ITC of 1.12 mM LC8 titrated into 10 µM KANK1_500-805_. (e) SV-AUC titration of KANK1_500-805_ and 90 µM LC8. (f) SEC-MALS of KANK1_500-805_, LC8, and their complex; y-axis is normalized to the tallest peak in the trace. Measured mass is in red with average mass listed. (g) Representative NSEM image of the KANK1_595-720_ bound to LC8. Scale bar, 100 nm. (g, right) Representative class averages, and corresponding schematic diagrams, showing seven domains in three conformations: straight (top), 6+1 L-shape (middle), and 5+2 L-shape (bottom). Scale bar, 20 nm. (h) NSEM images of negatively stained LC8-bound KANK1_500-805_ construct. (left) General view and (right) representative particles showing the transition from closed to open conformations. Scale bars: 100 nm and 50 nm, respectively. (i) 2D class average of KANK1_500-805_ bound to LC8 in a closed conformation showing 15 globular densities (left) with a schematic model (right). Scale bar, 20 nm. (j) Scatter dot plot of internal angles of LC8-bound KANK1_500-805_ proteins (left) showing conformational flexibility as illustrated in schematics (right). Raw data (n = 239) are plotted in red, with the median value (13.2°) indicated by a black line.

PyMOL alignment of LC8-saturated KANK1_500-615_, KANK1_541-650_, and KANK1_595-720_ suggests binding to 15 LC8 dimers leading to rigidification and elongation of the first 250 residues of L2 into a hinged rod (Fig. 6c), with an extended length of ∼53 nm. This requires experimental validation, but being greater than the distance between the cell cortex and microtubule ends. Importantly, no subconstruct of KANK1-L2 beyond KANK1_595-720_ displayed multivalent cooperativity in AlphaFold titrations (Figs. S10-13). For clarity, in AlphaFold titrations KANK1_700-810_ shows evidence of binding, but not multivalent cooperativity.

### Experimental verification of multivalent binding in KANK1_500-805_

To validate AlphaFold titrations we generated a construct spanning residues 500-805 (KANK1_500-805_). This region was selected because both pull-down assays (Fig. 3a,b) and the AlphaFold site predictions (Fig. 6a) indicate LC8 binding sites between residues 500 and 615 and between 720 and 800. Because KANK1_595-720_ binds seven LC8 dimers, we expected KANK1_500-805_ to also bind at least seven LC8 dimers. However, ITC titrations of LC8 into KANK1_500-805_ (Fig. 6d), revealed a striking increase in stoichiometry that far exceeds that of KANK1_595-720_, at n=17.98 ± 0.07. Additionally, the interaction is stronger for KANK1_500-805_ than for KANK1_595-720_, at K_d_ = 579 ± 65 nM, despite a slightly weaker enthalpy at -5.62 ± 0.06 kcal/mol.

To complement ITC, we performed SV-AUC and SEC-MALS experiments. SV-AUC (Fig. 6e) showed a major peak for the complex at a value of ∼7 S. Titration experiments showed that at lower LC8:KANK1 ratios, this large complex is in exchange with a smaller complex as indicated by the appearance of a second peak and a downward shift of the main peak. Interestingly, samples with lower LC8 ratios also exhibited higher levels of free LC8, suggesting that cooperativity across the complex is not uniform and that a critical concentration of LC8 is necessary to drive assembly from the initial state to the fully bound state. The AlphaFold titration model (Fig. 6c) supports the following interpretation: two LC8 binding regions, separated by the nuclear export domain, bind with some degree of independence from one another, allowing a tiered assembly pathway.

SEC-MALS (Fig. 6f) analysis of KANK1_500-805_ alone gave a mass of 34.9 kDa, matching its expected monomeric size. When 9 µM KANK1_500-805_ was mixed with 1000 µM LC8, the chromatogram displayed a dominant peak with a mass of 395 kDa, along with a convoluted set of three peaks close to the excluded volume with a mass over 1 MDa. The smaller mass corresponds to a complex of two KANK1 molecules and 15 LC8 dimers, in agreement with the AlphaFold predictions. The larger species likely represent higher-order assemblies which is not surprising given that sample preparation of these mixtures sometimes resulted in gel-like aggregates which separated from the solution. In the shorter constructs, similar aggregation was resolvable through optimization of protein and salt concentrations. For KANK1_500-805_ aggregation was unavoidable at all conditions.

Negative-stain electron microscopy of KANK1_595-720_ with LC8 showed micrographs dominated by straight rod-like structures consisting of seven segments along with 6+1 and 5+2 L-shape structures (Fig. 6g), consistent with the AlphaFold predictions for this complex. Analogous experiments with KANK1_500-805_ (Fig.6h-j), showed structures of 15 LC8 dimers, primarily forming a closed hairpin configuration with seven beads on one side and eight on the other, matching the overlay of AlphaFold predictions (Fig. 6c). A small population was assembled as partially open hairpins, indicating conformational flexibility, and access to the fully open hinge state. In these open conformations, particles exceed 50 nm in end-to-end length, surpassing the distance necessary to span the gap between the cell cortex and microtubule ends.

## Discussion

To investigate how the exceptionally long disordered L2 of KANK1 could bridge the cell cortex and microtubule ends and whether it can span the distance between these two structures, we performed extensive experimental and computational analyses of this region. Our cell-based studies demonstrate, for the first time, a direct interaction between the cytoskeletal protein KANK1 and the hub protein LC8. KANK1 recruits LC8 to the edges of focal adhesions, which connect intracellular actin bundles to the extracellular matrix. Given that KANK1 is a cortical adaptor involved in microtubule organization, this interaction represents a critical component of the crosstalk between focal adhesions and microtubules.

Here we elucidate the structural and biological roles of LC8 in this system and develop predictive and experimental approaches to characterize a rod-like assembly involving at least 15 LC8 dimers. Remarkably, this interaction was discovered serendipitously, without any canonical LC8 binding motifs, posing an opportunity to refine predictive algorithms and experimental strategies to study structure and thermodynamics of such complexes. We show that this multivalent assembly is highly cooperative, compositionally homogeneous, conformationally heterogeneous and capable of switching from disordered to extended rod-like structures. We discuss how this switch could be an essential property for bridging large cellular assemblies.

### Multivalent binding expands the sequence diversity of LC8 binding motifs

Our findings reveal that LC8 binds a far broader variety of motifs than previously appreciated. None of the motifs identified in KANK1 closely resemble known LC8 binding sequences, as evidenced by LC8Pred’s difficulty in producing confident predictions. The presence of at least 15 noncanonical binding sites in KANK1 suggests that LC8Pred’s current estimate of 374 LC8 binding sites in the human proteome(*37*) is a conservative underestimate. Further work should include systematic characterization of these motifs and others like them, through deletion studies, and incorporation of the expanded motif diversity and multivalent context into future algorithms like LC8Pred.

Such improvement will likely uncover an even larger selection of LC8 binding sites across the human proteome and in disease-causing pathogens.

### Conformational and compositional heterogeneity are encoded in the protein sequence

The 7:1 binding stoichiometry of LC8 with KANK1_595-720_ is reminiscent of ASCIZ, one of the most extensively studied multivalent LC8 partners(*50–52*, *58–60*). However, despite identical stoichiometry, their multivalent behaviors are quite different, offering insight into how different multivalent structures can inform function. Because LC8 dimerizes partners, one might expect all multivalent LC8 binding interactions to exhibit cooperativity due to increased effective concentration of the partner after forming the first LC8 “bridge”. Yet this is not observed for ASCIZ: its strong sites bind non-cooperatively and heterogeneously to form multiple partially occupied complexes that are more stable than the fully occupied complex. In contrast, KANK1’s weak sites undergo strong cooperativity that strengthens binding by two orders of magnitude over any individual site, resulting in homogeneous, fully occupied complexes.

Nup159 is another LC8 partner which exhibits short (4-5 amino acid) linkers, and serves a structural role. Previously it was considered to be the most homogeneous LC8 binder(*49*). However, Nup159 motifs are strong and easily predicted, yet they bind with a complicated interplay of positive and negative cooperativity leading to a stable, but entropically unfavorable complex. This results in a substoichiometric major species similar to ASCIZ, exemplifying the need for both bivalent binding enhancement and enthalpic compensation for entropically unfavorable interactions(*48*). Negative-stain electron microscopy reveals significant flexibility in its LC8-bound “rod”(*58*) which is rigidified in the context of the larger protein assembly(*49*). These comparisons underscore KANK1’s uniqueness among LC8 partners for forming homogeneous, high-affinity complexes despite relying on individually weak motifs.

We previously proposed(*60*) that both motif sequence and inter-motif linker length govern the cooperativity in multivalent LC8 interactions. Analysis of KANK1-L2 reinforces this principle, highlighting the importance of homogeneity in both binding site affinities and linker lengths for promoting homogeneity of complex assembly. We hypothesize that the weakness of each individual binding site precludes the possibility of alternative complexes assembling due to low stability in any sub-stoichiometric or misaligned complex making the fully saturated complex the only viable state. Furthermore, the short linkers in KANK1 (3-4 amino acids) appear optimized for rigidification and cooperativity, in line with our previous conclusions(*59*, *60*). Contrastingly, the mismatched linker lengths and diverse affinities in ASCIZ stabilize alternative complexes, at the expense of the fully occupied complex, explaining the heterogeneity and lack of cooperativity in LC8/ASCIZ binding.

Multivalent LC8 binding is not the only system for which negative control has been shown to be functionally important. For example, recent studies of collagen triple helix assembly revealed that in the hours-to-months long process of folding a triple helix, the rate limiting step is actually unfolding off-target triple helices, emphasizing the importance of negative design for destabilizing alternative helices to increase the helix formation rate of the target(*66*). The parallel between LC8 multivalency and collagen triple helical assembly illustrates a broader biological strategy for considering solutions that are not the strongest, most homogeneous, or most optimized. Overly stable folding could result in promiscuous function(*67*) and optimized affinities may increase off-target interactions(*68*). These findings highlight the delicate balance when considering complexes for which assembly is driven by multivalency.

### LC8 binding as a molecular switch

LC8 binding to the disordered L2 region of KANK1 rigidifies the linker and organizes it into two consecutive rod-like segments with a flexible hinge separating them. This rod formation elongates the first half of L2. We propose that elongation provides the necessary distance of separation between the coiled-coil domain, at one end of L2, which binds to liprins at the cell membrane, and the ankyrin domains, at the other end, which bind to KIF21A at microtubule ends (Fig. 7), to span the 35-50 nm gap(*32*). As an intriguing thought, depending on the binding order, the elongated KANK1/LC8 assembly binding to KIF21A may actually be the reason microtubule growth stops at 35-50 nm from the cell membrane, however, it may just be that this assembly needs to be this long to span a gap that is defined by other mechanisms. Planned experiments using shorter L2 regions will clarify this relationship.

**Fig. 7.**
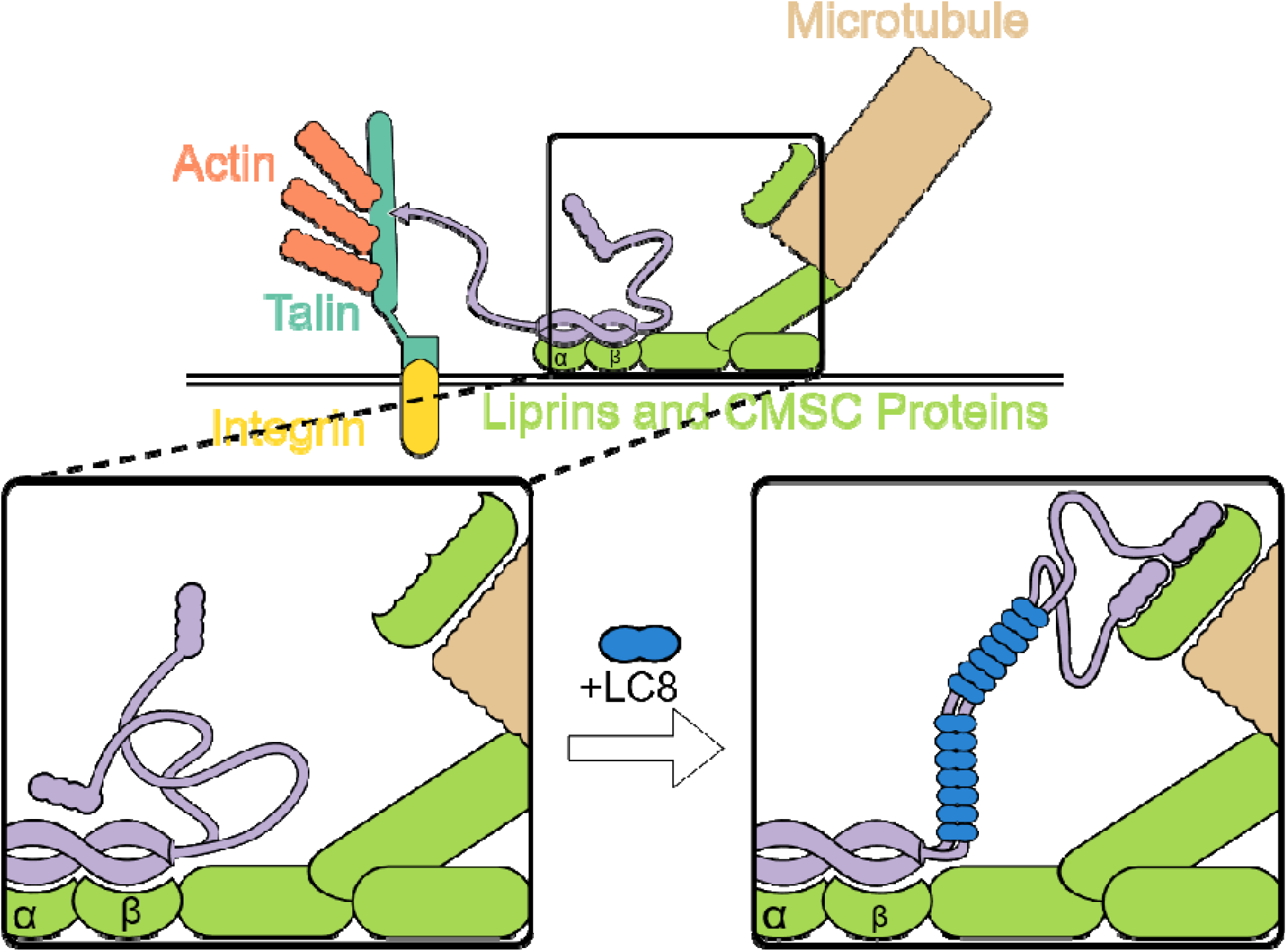
A model showing LC8 binding to KANK1 rigidifies L2 enabling reach between Liprins and KIF2A. Model of the protein complexes of which KANK1 is a central player bridging cytoskeletal components, recalling Fig. 1a,d for context. Inset shows more detail on the region impacted by LC8 binding. Upon the addition of LC8, L2 is rigidified and elongated to reach across the gap between the cell membrane and microtubules. End-to-end distance for a random coil is to 17 nm on average, whereas interaction with LC8 rigidifies and elongates L2 to >50 nm, allowing it to span the distance to KIF21A at the end of a microtubule.

It is noteworthy that KANK1 and KANK2 have similar affinity for KIF21A(*30*, *31*), and are both expressed in HeLa cells; yet KANK1, shown here to undergo LC8-induced elongation plays a more dominant role in controlling KIF21A localization and microtubule organization than KANK2, which does not bind LC8(*28*). This distinction underscores the functional significance of LC8-mediated structural remodeling in KANK1. It seems significant that KANK1-L2 is 281 amino acids longer than KANK2-L2 (Fig. 2b) and the LC8 binding domain in KANK1-L2 is 220-300 amino acids long, hinting that perhaps evolution specifically deleted this region from KANK2.

Short linkers between adjacent LC8 binding sites eliminate local flexibility and ensure optimal extension, whereas a single long linker between two rod segments likely allows for variation in membrane-microtubule distances and angles of approach. This hinge may also protect against fragility that could arise from a rigid uninterrupted 15-segment long rod. A natural question follows: “Rods are often formed by alpha helical coiled-coils, why wouldn’t biology use that same strategy here?” There are several factors that favor using LC8-mediated assembly over an intrinsic coiled-coil: (1) Sequence economy: in binding LC8, KANK1 takes only 220 amino acids to make a structure that spans over 50 nm, whereas a coiled-coil requires over 330 amino acids per alpha helix to achieve the same distance. (2) Efficiency: While production of 15 LC8 dimers is cellularly expensive, the LC8 being used in this interaction is already readily present in the cell(*37*) and it can be recycled in other interactions throughout the cell, so the LC8 needed for this rod requires minimal additional cost. (3) Dynamic adaptability: LC8-mediated rods can assemble and disassemble or allow easier access to a built-in zone of flexibility, or a hinge, in the rod to enable a wide range of orientations for reaching out to engage microtubules. Regardless of the reason, biology employed LC8 in this situation as a tool for rigidifying and lengthening a partner. This mechanism highlights LC8 as a versatile architectural module in cellular design, raising the intriguing question as to whether similar LC8-driven mechanisms are at work in other large protein assemblies.

### Summary

KANK1 emerges as a unique LC8 interactor, where binding to at least 15 LC8 dimers places it among the highest occupancy LC8 partners. No other LC8 partner exhibits the extreme cooperativity among individually weak motifs, forming protein rods that are readily visualized by negative stain electron microscopy. This architecture reveals how short, uniform linkers and distributed weak sites can together drive cooperative multivalent assembly. Our findings underscore the functional importance of weak interactions, and the role of short consistent linkers in promoting cooperativity. It also illustrates how multivalency and structural adaptability can serve as design principles in cellular architecture.

## Materials and Methods

### Worm-like Chain Assessment

The root mean squared end-to-end distance of a worm-like chain is represented by the equation Eq. 1

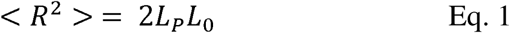

where R is the end-to-end distance, L_P_ is the persistence length, and L_0_ is the maximum length of the chain. L_0_ can be represented by Eq. 2,

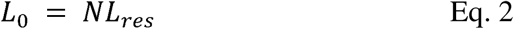

where N is the number of segments in the chain (number of peptide bonds), and L_res_ is the length of each segment. For IDPs, Lapidus *et al*. measured P as 0.64 nm(*34*). The length of a peptide bond has been measured as 0.38 nm. The range in distance reported was calculated by varying P as 0.64 or 1 and varying L_res_ as 0.38 or 0.34. The proportion of the ensemble at a given length, r is given by the equation Eq. 3(*36*)

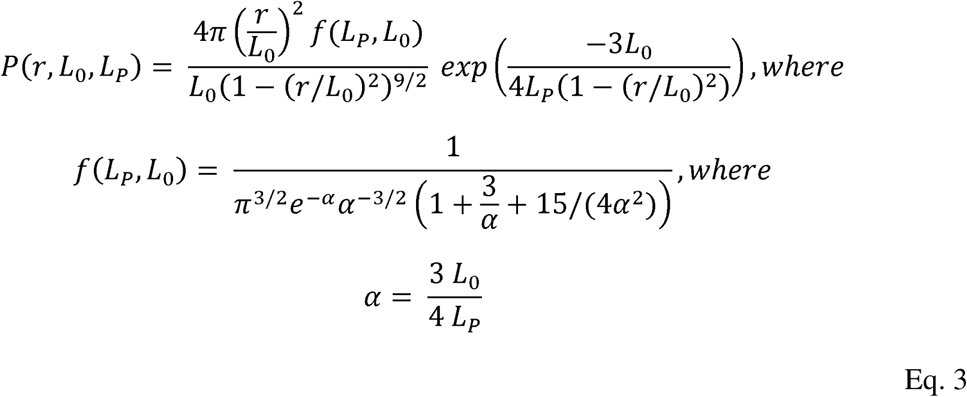

The explicit integral of P(r) was calculated from 35 to L_0_ and from 50 to L_0_ with Wolfram Alpha to calculate the proportion of the ensemble population above 35 and 50 nm, respectively.

### Cell culture and DNA/siRNA transfections

HeLa (Kyoto) cells stably expressing LC8-GFP were a kind gift from I. Poser and A. Hyman (Max Planck Institute of Molecular Cell Biology and Genetics, Dresden, Germany)(*69*). HeLa cells were cultured in DMEM medium. HEK293T cells were cultured in DMEM:F10 medium (1:1, v/v). All media were supplemented with 10 % (v/v) fetal calf serum and with Penicillin/Streptomycin (100 units/mL penicillin and 100 μg/mL streptomycin). All cells were grown at 37 °C in humidified incubators at 5% CO2 atmosphere. The cells were routinely checked for mycoplasma contamination using the Mycoalert assay (Lonza, LT07-518).

Transfection of siRNA into HeLa cells and of DNA into HEK293T cells was performed as previously described(*28*). In brief, HeLa cells were transfected with siRNAs using HiPerFect (Qiagen). The siRNA-transfected cells were analyzed 48 – 72 hours after transfection. The cells were analyzed 24 hours after DNA transfection. HEK293T cells were transfected with DNA using polyethyleneimine (PEI). The cells were harvested for pulldown assays around 24 hours after transfection.

### DNA constructs and siRNAs

The BioGFP-tagged KANK1 and KANK2 fusion constructs were made using PCR-based amplification of KANK1 or KANK2 fragments. These fragments were cloned into pBioGFP-C1 vector using Gibson Assembly mix (New England Biolabs) as described previously(*25*). The LC8-mCherry fusion construct was made by PCR-based amplification of LC8 and cloning it into the pmCherry-N1 vector.

The same siRNAs against KANK1 and KANK2 were used as described previously(*25*) to deplete HeLa LC8-GFP cells of KANK1 and KANK2: control siRNA (siLuciferase) GTGCGTTGCTAGTACCAAC; siKANK1 CAGAGAAGGACATGCGGAT; siKANK2 ATGTCAACGTGCAAGATGA.

### Streptavidin-based pulldown assays

Streptavidin-based pulldown assays of biotinylated BioGFP-tagged-KANK constructs expressed in HEK293T cells were performed as previously described(*28*). In brief, HEK293T cells were seeded in 10 cm plastic culture dishes to ∼80 % confluence, the cells were allowed to attach overnight. The cells were then transfected with the BioGFP-tagged KANK constructs together with biotin-ligase BirA (5μg DNA per construct) using PEI. In parallel, 1 – 2 10 cm plastic culture dishes were transfected with the prey protein only (LC8-mCherry, or HA-KANK1-FL, 10μg DNA) using PEI. The DNA/PEI suspensions were added to the cells dropwise. Thereafter, the cells were incubated for 24 hours at 37 °C in 5 % CO2 atmosphere in humidified incubators. After incubation, the cells were harvested on ice using ice-cold PBS and scraped from the dishes, pelleted and subsequently lysed in a non-ionic detergent-based lysis buffer (50 mM HEPES pH 7.4, 150 mM NaCl, 1 mM EDTA, protease inhibitor cocktail (Roche), 1 % Triton-X100) for 10 minutes on ice. Afterwards, the suspension was centrifuged at 13,200 rpm at 4 °C for 15 minutes. In the meantime, the magnetic streptavidin beads (ThermoFisher, 11206D) were blocked in lysis buffer supplemented with 5% chicken egg white (Sigma) for 30 minutes at RT. Thereafter, the lysate of the prey protein was equally distributed among the biotinylated GFP-tagged KANK constructs and then incubated with the streptavidin beads for 2 hours at 4 °C, except for 7 % of each lysate mix which was kept as “input” control.

DTT-containing SDS-PAGE protein sample buffer was added to the input control. After incubation of the lysates with the streptavidin beads, the beads were washed five times using the lysis buffer and a magnetic “DynaMag” rack (ThermoFisher, 12321D) before adding DTT-containing SDS-PAGE protein sample buffer.

SDS-PAGE and Western blot analysis were performed according to standard procedures. Commercially available antibodies against GFP (Abcam, ab6556), mCherry (clone 1C51, Abcam, ab125096) and HA (clone 16B12, BioLegend/Covance, MMS-101P) were used. For detection, near infrared fluorescence technology was used (Li-Cor Biosciences, Odyssey system), with infrared dye conjugated goat antibodies against mouse and rabbit (Li-Cor Biosciences).

### siRNA-mediated knockdown of KANK1 and KANK2 in HeLa LC8-GFP cells

HeLa LC8-GFP cells were seeded in 6-well culture plates and transfected using 20nM per siRNA per well using HiPerFect (Qiagen) and adding the siRNA/transfection reagent mix to the cell suspension. The cells were allowed to incubate in the siRNA/transfection reagent-containing medium overnight. The cells were then washed twice with PBS and incubated for another 24 hours. The following day (∼48 hours after siRNA transfection), the cells were trypsinized and seeded on 12mm glass coverslips (10,000 – 15,000 cells per coverslip) and allowed to attach overnight. The following day (∼72 hours after siRNA transfection), the cells were washed with PBS and fixed for immunofluorescence staining.

### Antibodies and Immunofluorescence staining

Cell fixation and staining were performed as previously described(*28*). In brief, the cells were fixed with ice-cold methanol for 7 minutes at -20 °C. After washing once with PBS, the cells were permeabilized for 4 minutes at room temperature (RT) using permeabilization buffer (0.2 % TritonX-100 (Sigma) in PBS). The cells were then blocked in blocking buffer (0.05 % Tween-20 (Sigma) and 2 % BSA (Carl Roth) in PBS) for 1 hour at RT, followed by incubation with primary antibodies for 1 hour at RT. After incubation, the cells were washed five times for two minutes with wash buffer (0.05 % Tween-20 in PBS); followed by 1 hour incubation with the fluorescently labeled secondary antibodies diluted in blocking buffer. The cells were washed five times for two minutes with wash buffer and afterwards, dehydrated using first 70 % and then 100 % ethanol. Finally, the coverslips were mounted on glass slides using Vectashield mounting medium (Vector laboratories, H-1000).

Commercially available antibodies against KANK1 (Atlas antibodies, HPA005539), KANK2 (Atlas antibodies, HPA015643), and paxillin (clone 165, BD Biosciences, 610620) were used. Alexa-405, Alexa-488 and Alexa-594 conjugated goat antibodies against rabbit and mouse IgG were purchased from Invitrogen.

### Microscopy and image analysis

Images of fixed cells were collected with a Nikon Eclipse Ni upright wide field fluorescence microscope and a Nikon DS-Qi2 CMOS camera (Nikon), using Plan Apo Lambda 100x N.A. 1.45 oil objective (Nikon) and Nikon NIS (Br) software (Nikon). Nikon Intensilight C-HGFI was used as a light source. For imaging of blue, green and red fluorescence the filter sets ET-BFP2 (49021), ET-GFP (49002), and ET-mCherry (49008) (all Chroma) were used, respectively. For presentation, images were adjusted for brightness and contrast using ImageJ 1.50b (NIH).

### Protein Expression and Purification

For biophysical assays, LC8 and KANK1_595-720_ constructs were expressed recombinantly in *E. coli*. Plasmids were cloned into the pET24d expression vector, with an N-terminal 6xHis and TEV-cleavable site and expressed in either Rosetta DE3 (LC8) or C41 DE3 (KANK1) *E. coli* cells (both are derived from BL21 DE3 cells). Cells were grown in ZYM5052 auto-induction media at 37 °C for 24 hours. LC8 and KANK1_595-720_ were both purified by affinity chromatography on TALON cobalt resin, as previously described(*50*). KANK1 constructs express into inclusion bodies and were therefore purified using buffers containing 8 M urea as previously described(*50*). Following affinity purification, these were dialyzed out of urea, and both KANK1 and LC8 were further purified by size exclusion chromatography (SEC) using a Superdex S75 hi-load column (GE Healthcare).

For KANK1_500-805_, a synthesized pET 24d containing a 10x histidine tag, cleavage site for the tobacco etch virus (TEV) enzyme, and codon optimized, human KANK1_500-805_ sequence was generated by GenScript Biotech. The plasmid was transformed into *E. coli* Rosetta 2 (DE3) plysS cell and culture was grown in 2xYT medium at 37 °C until OD_600nm_ reached 0.8-0.9. Recombinant protein expression was induced with 0.5 mM IPTG and growth continued at 37 °C for 5 hours. The cells were harvested and frozen at -80 °C.

KANK1_500-805_ was purified under denaturing conditions using Ni-NTA agarose (Qiagen, Cat. No. 30210). Cell pellet was thawed on ice for 15 min and resuspended in native lysis buffer (50 mM sodium phosphate, 300 mM sodium chloride, pH 8.0) with 1 mg/ml lysozyme, 5 µg/ml DNase I, 10 µg/ml RNase A, and protease inhibitor at 5 ml per gram wet weight. The cell suspension was incubated on ice for 30 min, then the cells were disrupted by sonication (12 x 10 sec with 30 sec cooling in between). The cell lysate was centrifuged at 10,000 g for 30 min at 4 °C. After discarding the supernatant, the pellet was washed twice with 20 mM Tris-HCl, pH 8.0. To completely resuspend the inclusion bodies (IB), the IB suspension was briefly sonicated (3 x 10 sec), then centrifuged at 12,000 g for 20 min at 4 °C each time. The IB pellet was suspended in freshly made denaturation buffer (8 M Urea, 100 mM NaH_2_PO_4_, 10 mM Tris, pH 8.0) and incubated on a shaker for 30 min. The solubilized IB was centrifuged at 10,000 g for 10 min at RT to remove any insoluble debris. The supernatant was mixed with equilibrated Ni-NTA agarose by mixing on a rotator for 30 min at RT. Above mixture was loaded into an empty BioRad column and the settled resin was washed with 5 bed volumes of 6 M urea, 100 mM NaH_2_PO_4_, 10 mM Tris, pH 8.0, followed by 5 bed volumes of 4 M urea, 100 mM NaH_2_PO_4_, 10 mM Tris, pH 8.0. The protein was eluted in 3 M urea, 250 mM imidazole, 100 mM NaH_2_PO_4_, 10 mM Tris, pH 8.0. The elution was diluted to 2 M urea, then dialyzed in 25 mM Tris HCl pH7.5, 150 mM sodium chloride, 1mM sodium azide, 1 M urea for six hours. TEV protease was added before dialysis in 25 mM Tris-HCl pH7.5, 150 mM NaCl, 1mM sodium azide buffer overnight. TEV protease and histidine tag were removed by binding to Ni-NTA agarose. Purified protein was over 95% pure as analyzed by SDS-PAGE.

### Buffer conditions and protein handling

For biophysical experiments, unless otherwise stated, all buffers were 25 mM Tris, pH 7.5, 150 mM NaCl, and 1 mM NaN_3_. With shorter constructs, 5 mM β-mercaptoethanol was also added. Purified proteins were stored in buffer and either used within a week or flash frozen to -80 °C for storage.

### Isothermal titration calorimetry

We performed isothermal titration calorimetry at 25 °C with a VP-ITC microcalorimeter (Microcal). For KANK1_595-720_, LC8 was titrated into a cell sample of KANK1_595-720_, with a syringe and cell concentration of 400 and 4 μM, respectively. Each injection had an 8 μL volume, and a total of 36 injections were performed. For KANK1 peptides, which were synthesized in-house using solid-phase synthesis and purified by HPLC, each peptide was dissolved into buffer at 500 μM and titrated into a cell sample of LC8 at 60 μM for a total of 28 injections, at injection volumes of 10 μL. Peaks were integrated and fit to the n-independent sites model in Origin 7.0.

For KANK1_500-805_ a TA Instruments Affinity-ITC was used, and data were collected at 25 °C. LC8 was titrated into KANK1_500-805_ in triplicates with slightly different parameters: Run 1 with 1.12 mM LC8, 10 μM KANK1_500-805_, and 15x3 μL injections, Run 2 with 1.12 mM LC8, 10 μM KANK1_500-805_, and 25x2 μL injections, and Run 3 with 1.12 mM LC8, 9 μM KANK1_500-805_, and 25x2 μL injections. Data was fit using the independent sites model in TA instrument’s software NanoAnalyze. Fit values of each replicate were consistent with one another and reported for Run 1. As a note, NanoAnalyze has a maximal fit stoichiometry of 10, so the fit was calculated with twice the cell concentration and then the stoichiometry was doubled. This ensures that all other values are fit correctly and allows for a simple adjustment to the stoichiometry. Because NanoAnalyze cannot access a high enough stoichiometry for this system, the fit line in the figure was produced with the ITC Planning tool from the Tsodikov laboratory(*70*).

### Analytical ultracentrifugation

We performed sedimentation velocity analytical ultracentrifugation (SV-AUC) on a Beckman Coulter Optima XL-A analytical ultracentrifuge, equipped with optics for absorbance measurements. We mixed purified LC8 and KANK1_595-720_ at a series of increasing concentrations, with a fixed KANK1_595-720_ concentration of 7.2 μM, and at increasing LC8 concentration varied from 9 μM (1.25:1) to 144 μM (20:1). The 1.25, 2.5, and 3.75:1 complexes were monitored by absorbance at 280 nm, 5 and 10:1 used absorbance at 292 nm, and the 15 and 20:1 complexes used 298 nm. For KANK1_500-805_ alone, 9 μM protein was collected at 280 nm. In complex with LC8, concentrations of 90:18, 90:9, and 90:6 μM LC8:KANK1_500-805_ were collected at 290 nm.

Samples were dialyzed the night before SV-AUC was performed, to ensure minimal β-mercaptoethanol degradation. Samples were loaded into 12 mm path-length 2-channel cells, and centrifuged at 42,000 rpm and 20 °C. We acquired 300 scans at the relevant wavelength, with no delay between scans. Absorbance profiles were fit to a c(S) distribution in SEDFIT(*71*), with a calculated buffer density of 1.0009 g/ml, calculated using Sednterp(*72*).

### Size exclusion chromatography – multiangle light scattering

SEC-MALS experiments were performed on a 10/300 Superdex 200 analytical SEC column (GE Healthcare) attached to an AKTA-FPLC (GE Healthcare) and routed through a DAWN multiangle light scattering and Optilab refractive index system (Wyatt Technology). We equilibrated the system to our buffer, then injected 100 μL of sample onto the column. For KANK1_595-720_ alone we injected a 30 μM sample and for the titrations we injected 2 μM KANK1_595-720_ with increasing concentrations of LC8.

KANK1_500-805_ was run at 196 μM alone and at a 9:1000 μM ratio in complex with LC8. Molar masses were estimated in the ASTRA software package, using a Zimm scattering model.

### AlphaFold LC8 motif predictions

LC8 binding motifs predictions were achieved as previously described(*61*). Briefly, the sequences of the proteins to be investigated were parsed into 16 amino acid long segments with 8 residues of overlap between successive parsed peptides. The resultingly parsed peptides were then used in AlphaFold predictions (run with a locally installed instance of ColabFold(*73*)) in which an LC8 dimer was predicted with one and with two copies of the parsed peptide to simulate the half-bound and the fully bound state. In-built AlphaFold metrics were then extracted and compared to thresholds previously derived for both exclusive (low false positive, high false negative) and inclusive (low false negative, high false positive) cutoffs.

### AlphaFold protein titrations

Two copies of sequences of full proteins or of sub-constructs were predicted with increasing numbers of LC8 dimers up to and beyond the number of binding sites that were identified by the motif prediction analysis. Heatmaps of the predicted aligned error (PAE) of the highest ranked prediction were analyzed and compared throughout the titration to confirm AlphaFold confidence and saturation point. Due to the quality performance of AlphaFold with protein segments of ∼120 residues in length we also ran all sequences that were longer than 300 residues in parsed sections of 100-130 residues in length to confirm the lack of binding observed was not due to the length of protein used. PAE heatmaps of the best predictions for each titration point of all titrations run are included in the Supplementary Material (Figs. S3-13).

### Negative stain electron microscopy

LC8 and KANK1_595-720_ were combined at a 7:1 ratio at an estimated complex concentration of 12 μM. Samples were kept on ice and diluted in buffer to 1:1000. LC8 and KANK1_500-805_ were combined at 90 and 6 μM concentrations, respectively. Samples were kept on ice and diluted in buffer to 1:10 and 1:50. Four microliters of the diluted sample were placed on a glow discharged copper grid for 30 seconds. The grid was blotted on the edge with filter paper, then rinsed quickly in two drops of buffer. They were next negatively stained with uranyl formate for 20 seconds and blotted dry from the edge with filter paper. Grids were imaged at 80 keV on an FEI Titan ChemiSTEM TEM (ThermoFisher, Hillsboro, Oregon).

### Angle measurements and single-particle analysis

Single-particle analysis was performed using RELION-4.0(*74*). For KANK1_595-720_, with seven LC8 molecules, the images were taken with a pixel size of 1.978 Å. Particles in three different conformations were processed separately. First, 24 straight-conformation particles were picked manually and extracted with a box size of 220 pixels. These particles were classified into four classes by 2D classification using a 400 diameter mask and a T value of 3. The best class average containing twelve particles is shown in Fig. 6g. Next, 43 L-shaped (6+1) particles were manually picked, extracted similarly, and classified into 10 classes using a 372 diameter mask and a T value of 3. The best class, having 17 particles, is shown in Fig. 6g. Finally, 10 L-shaped (5+2) particles with similar appearances were picked manually, boxed in the same way, and then averaged as one class using a mask diameter of 372 and a T value of 2.9 (Fig. 6g).

For KANK1_500-805_, with 15 LC8 molecules, 627 particles in the closed conformation were picked manually from the micrographs (pixel size: 1.978 Å). The closed particles were selected since they showed homogenous structures. Extracted particles were boxed with a box size of 314 pixels. Then, four rounds of 2D classification were performed. First, particles were classified into 15 classes using a mask diameter of 586 Å and a T value of 2. Subsequently, the best classes were subjected to sub-classification into 30 classes (a 586 Å diameter mask and a T value of 3), repeated twice. Finally, a further sub-classification was performed into 7 classes (mask diameter: 586 Å, T value: 3). After these rounds of classification, the class average containing 53 particles was obtained (Fig. 6i).

The internal angle at the bending point of each particle was measured using ImageJ (NIH). In total, 239 particles were analyzed. The graph was prepared using GraphPad Prism (version 9.2.0).

## Acknowledgments

We are grateful to Rebecca Jackson of the OSU electron microscopy facility for staining and collecting NSEM images, and Lena Kinion for help with synthesis of and ITC measurements on the peptides.

## Author contributions

Conceptualization: EJB, AA

Methodology: DRW, AE, Y-CA, YS

Investigation: DRW, AE, Y-CA, YS, QHL, PNR

Visualization: DRW, AE, Y-CA, QHL

Supervision: EJB, AA, MI

Writing—original draft: DRW, AE, Y-CA

Writing—review & editing: DRW, AE, MI, AA, EJB

## Data and materials availability

All data are available in the main text or the supplementary materials.

## Supplementary Materials

Figures S1 to S13 can be found in the associated Supplementary Materials.

