## Supplemental Figures 1-13 for "Cooperative multivalency converts disorder into rods, resolving a paradox in cellular architecture"

Supplementary Materials for

**Multivalent cooperativity converts intrinsic disorder into a rigid nanorod, resolving a paradoxical spatial mismatch in cellular architecture**

Douglas R. Walker *et al.*

**This PDF file includes:**

Figs. S1 to S13

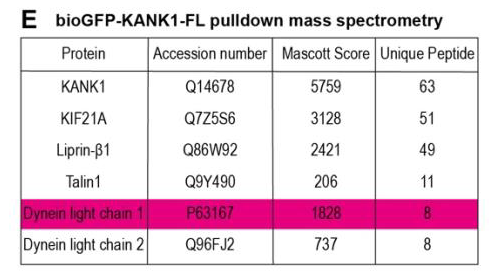
Fig. S1. Identification of the binding partners of bioGFP-tagged full-length KANK1 by using streptavidin pull down assays from HEK293T cells combined with mass spectrometry. The dynamic hub protein dynein light chain LC8 is indicated in magenta. KANK1 is included in the listing as its mass was measured in the mass spec as well.

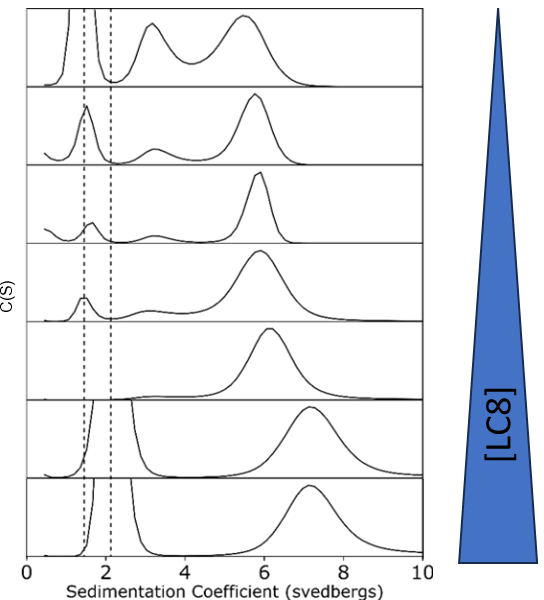

Fig. S2. A series of sedimentation velocity AUC experiments performed at 7.2 µM KANK1_595-720_, and increasing ratios of LC8. Dotted lines show centers of AUC peaks for KANK1 and LC8 in absence of their binding partner. To account for variation in the measured wavelength at high LC8 concentrations, Y axes are normalized to the tallest bound state (<2.5 svedbergs) peak. Triangle indicates that LC8 concentration increases as one moves down the panels.

Fig. S3. PAE heatmaps of titration points in the AlphaFoldMultimer titration of LC8 with KANK1_595-720_. Each heatmap corresponds to a different titration point, where KANK1_595-720_ was predicted with 1-8 LC8 dimers. Heatmap colors indicate highest confidence in violet and lowest confidence in red. Heatmaps are sized such that each LC8 dimer is similarly sized across all heatmaps. Importantly, LC8 monomers are violet across all predictions and dimers remain highly confident (violet to dark blue) even at high titration points as well. Consistent with KANK1_595-720_ binding to 7 LC8 dimers experimentally, the prediction’s confidence drops off significantly at a 1:8 ratio. The drop in confidence partway through the titration (points 3 and 4) likely are indicative of cooperativity stabilizing the complex for 5, 6, and 7 bound LC8.

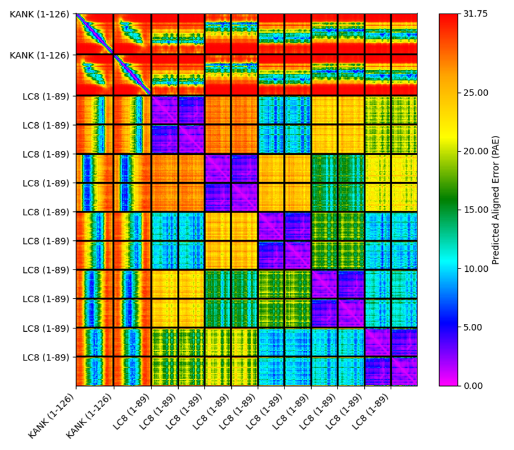

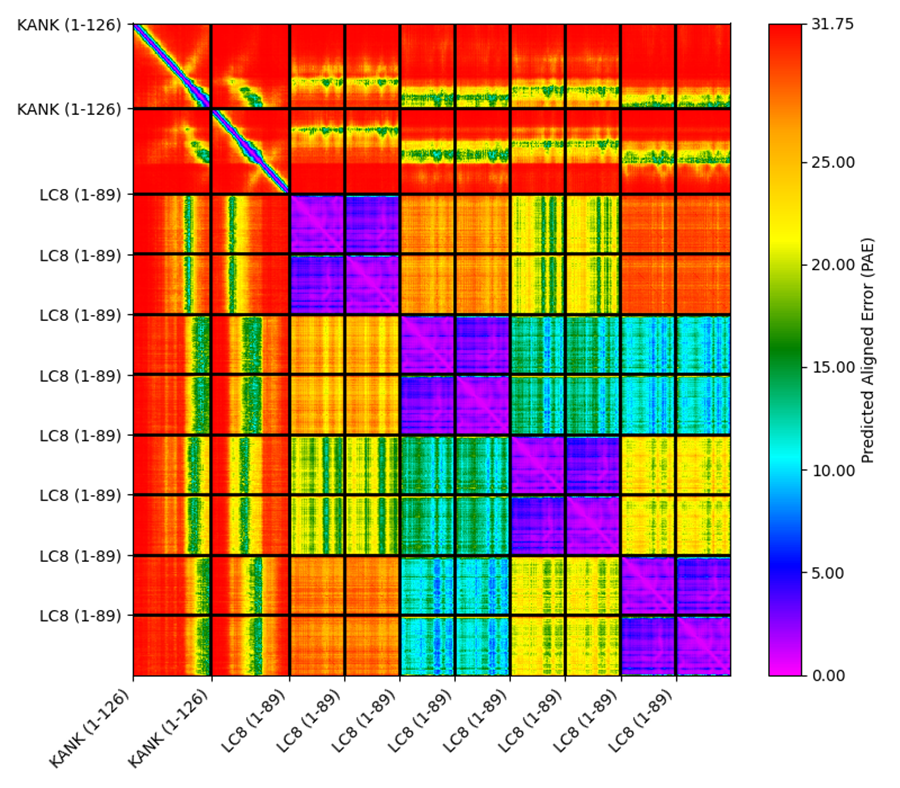

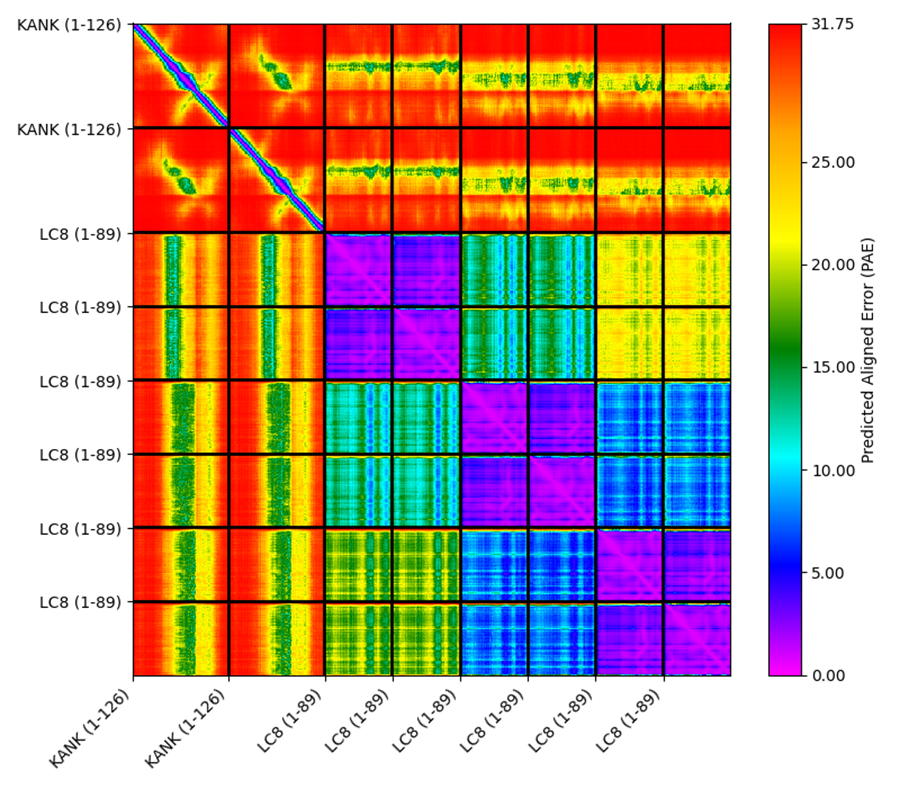

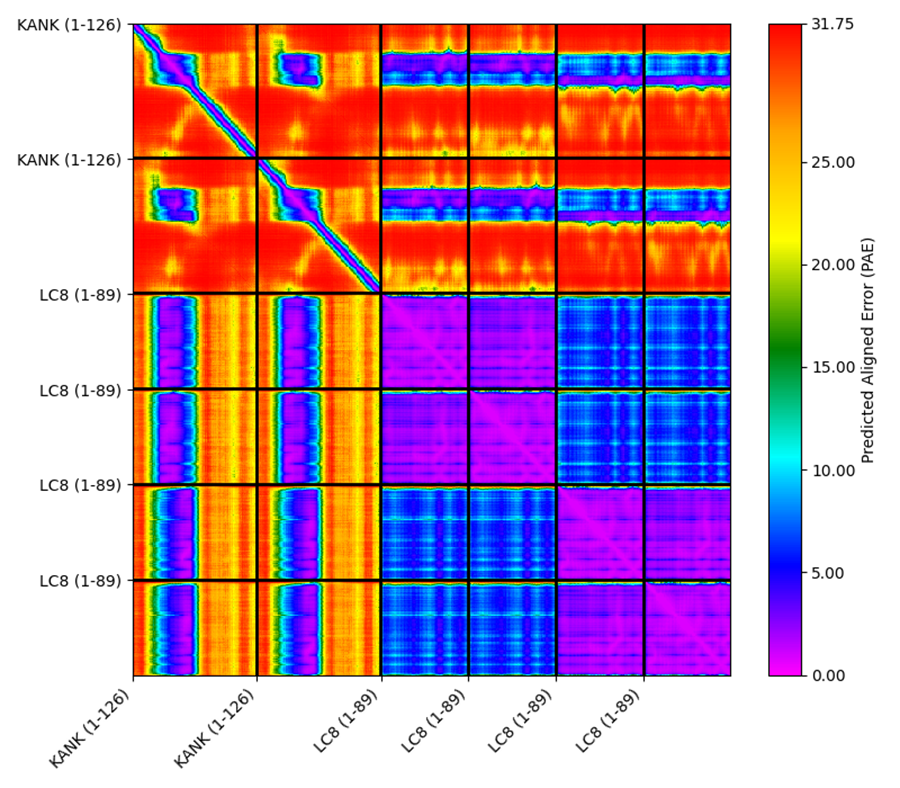

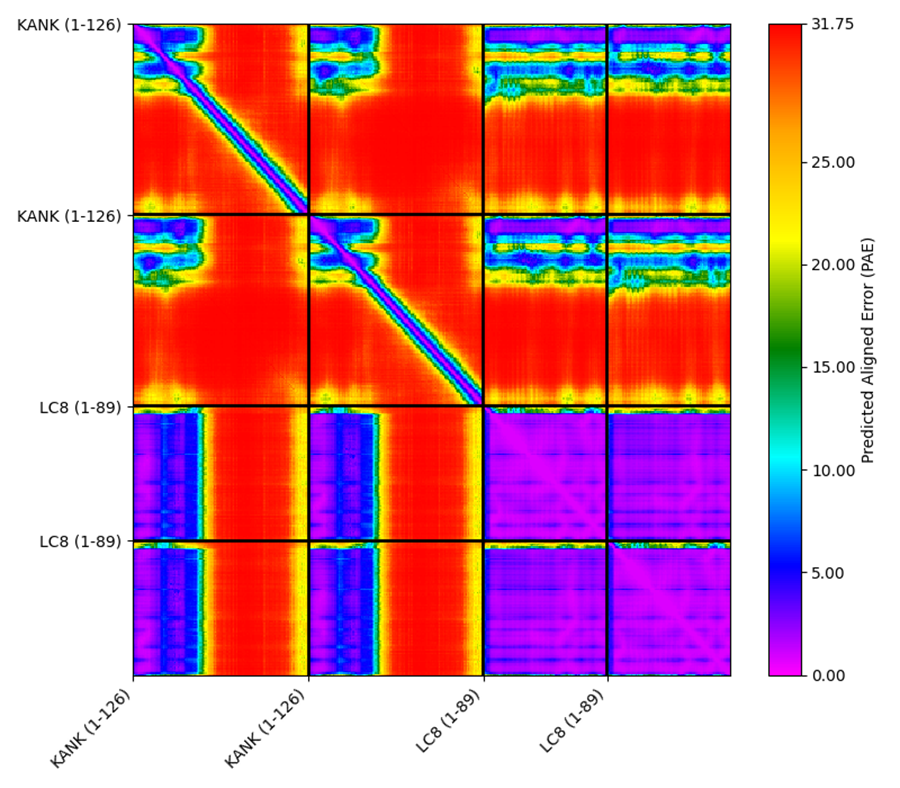

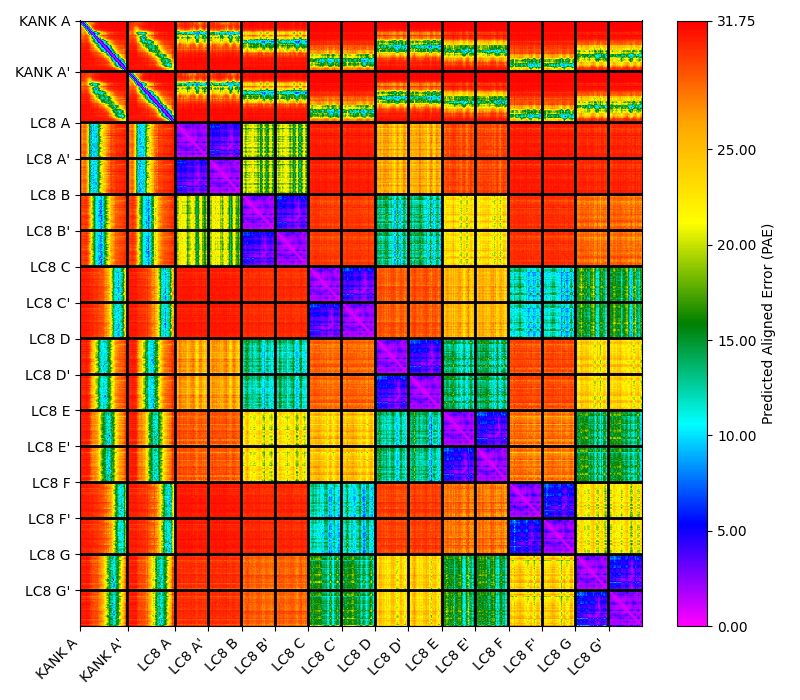

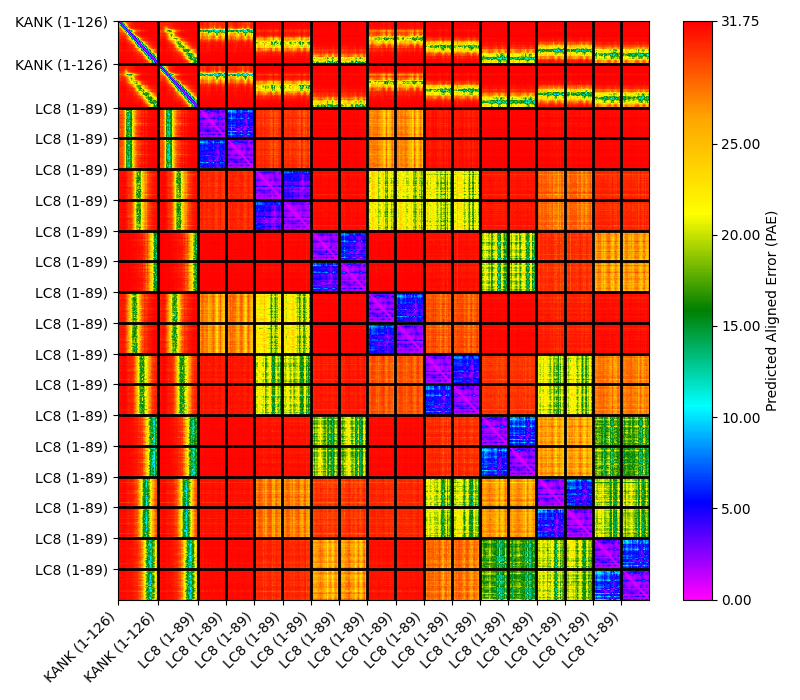

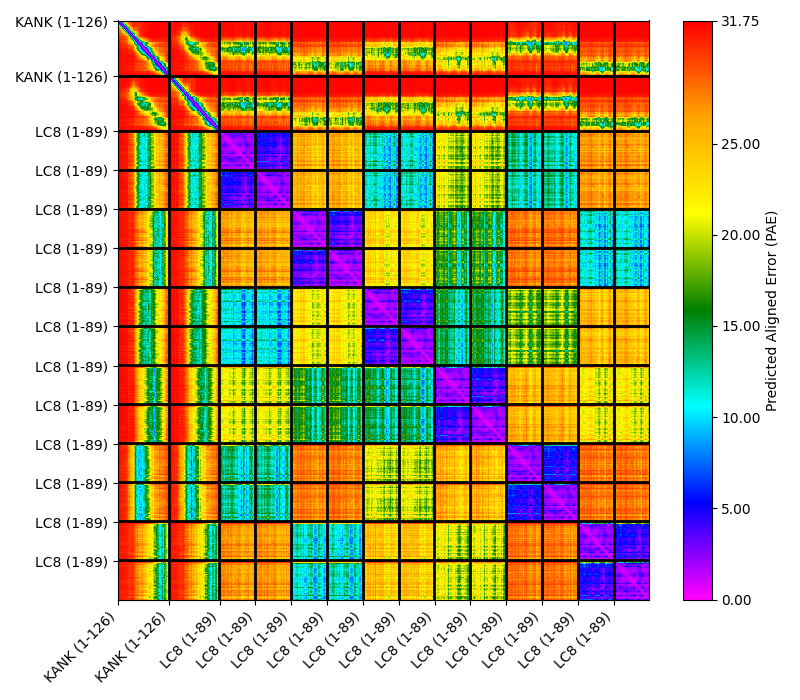

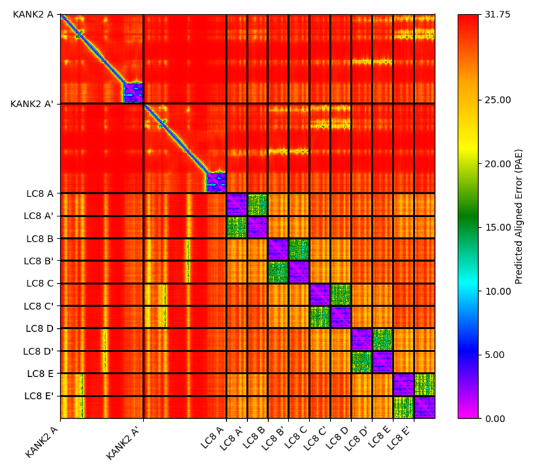

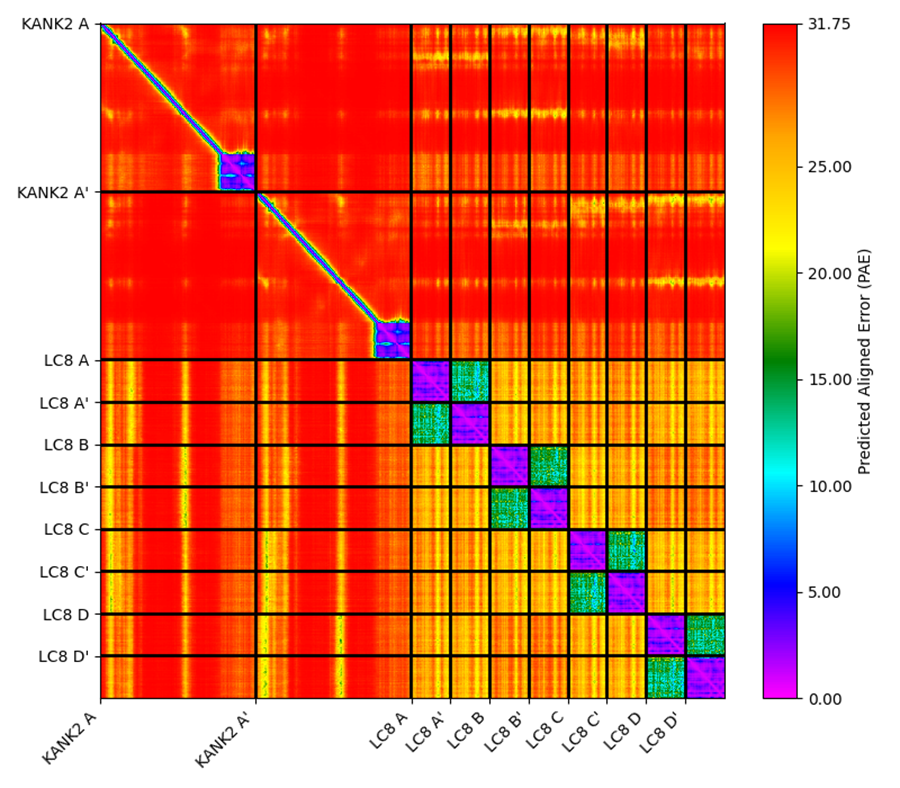

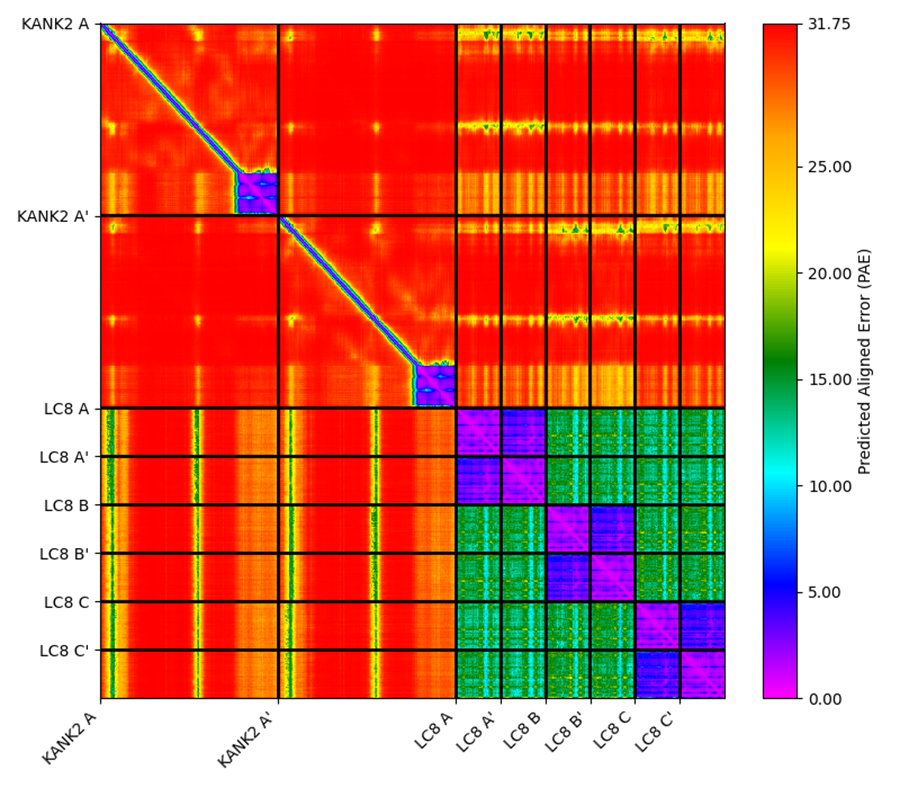

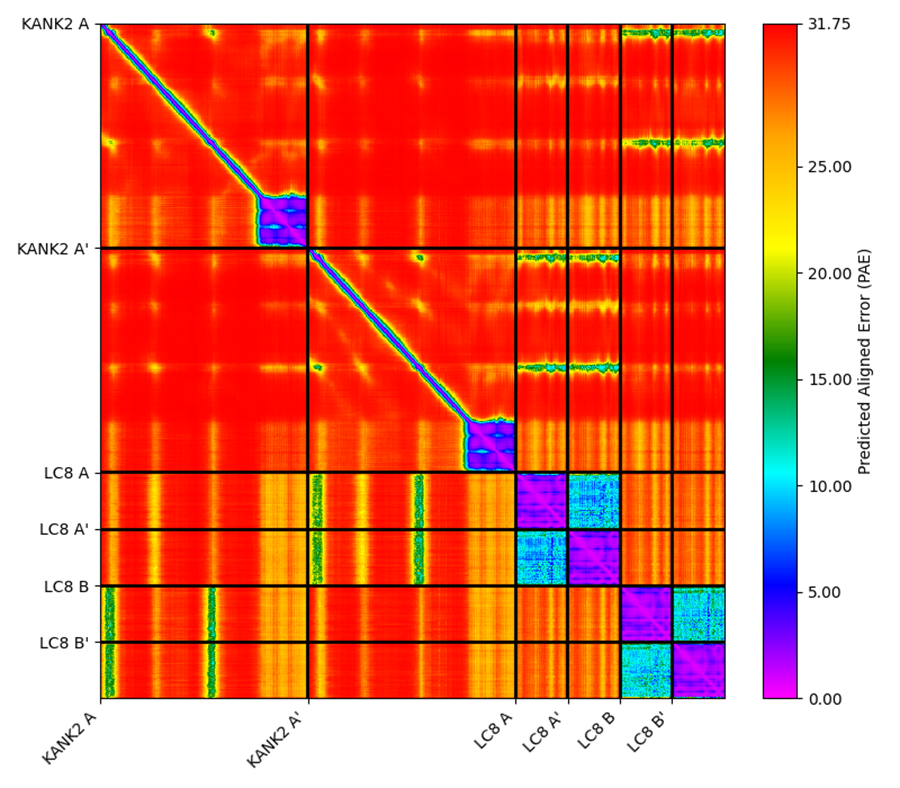

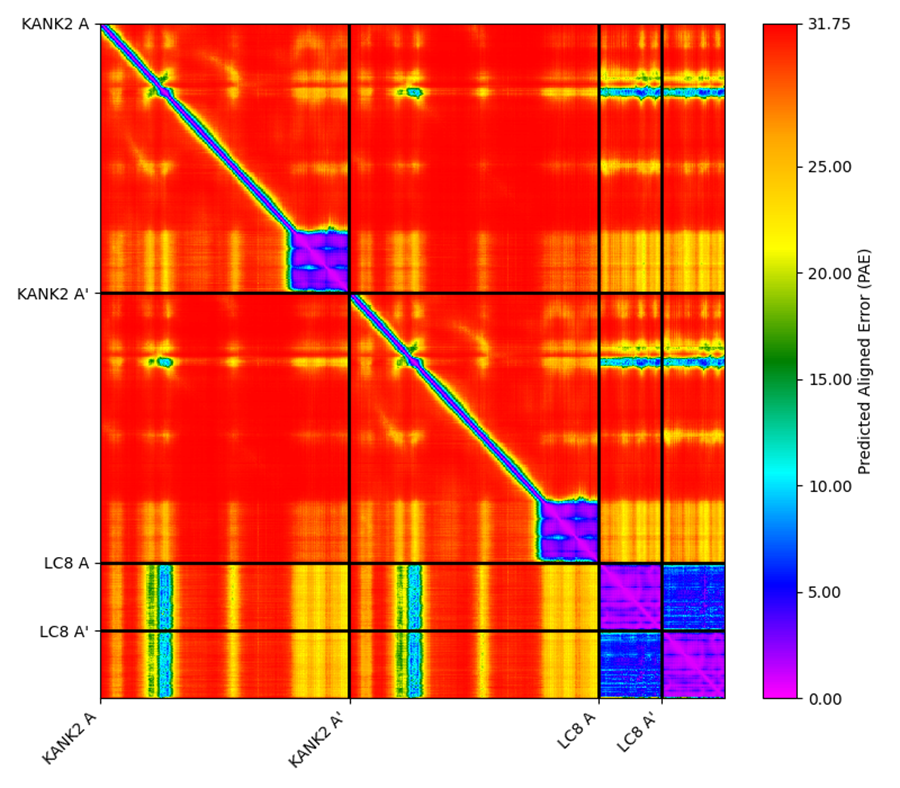

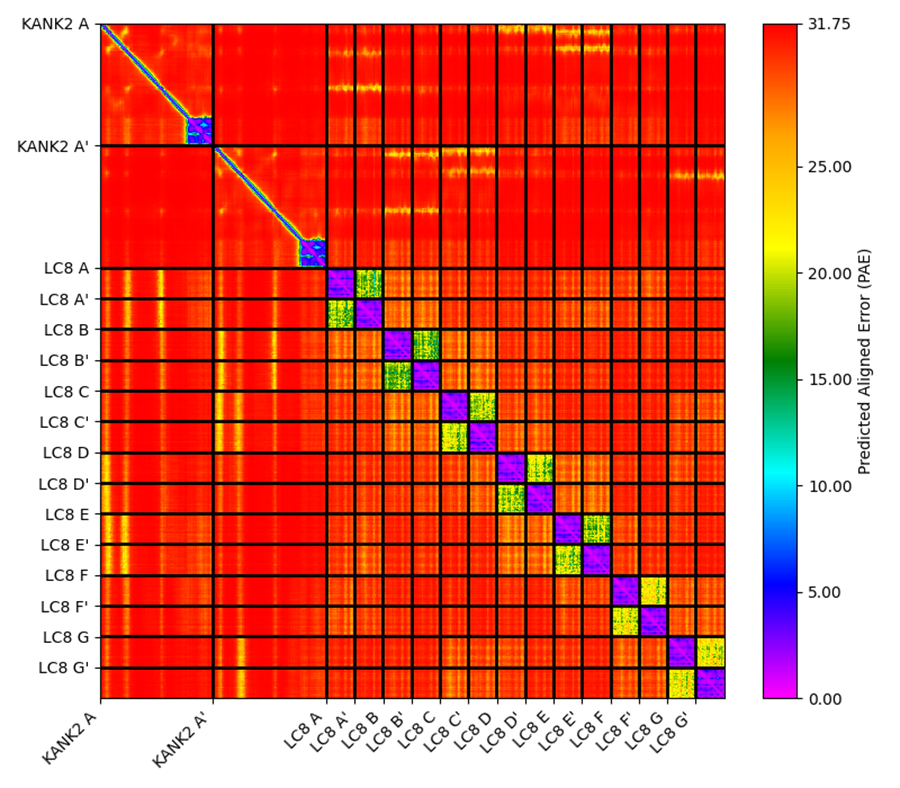

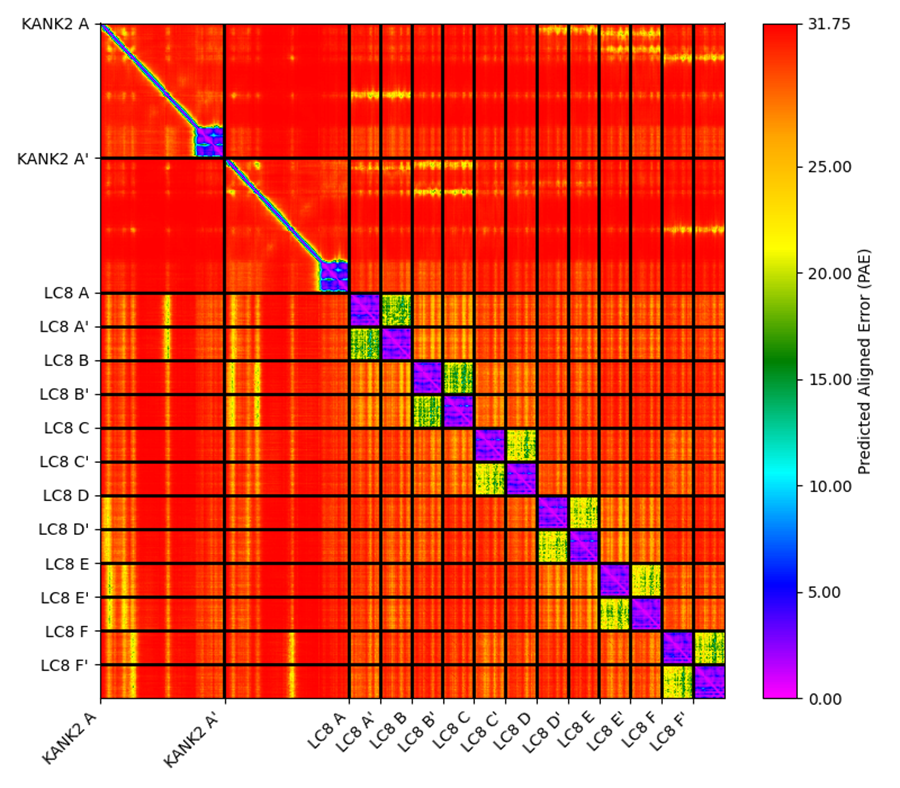

Fig. S4. PAE heatmaps of titration points in the AlphaFoldMultimer titration of LC8 with KANK2_L2_. Consistent with experiments, the predictions are not confident in KANK2/LC8 interactions at any point in the titration.

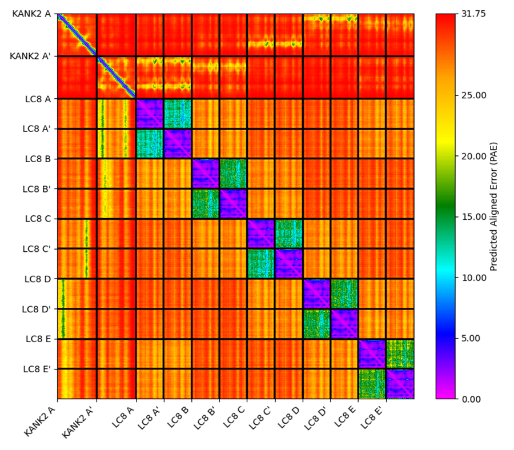

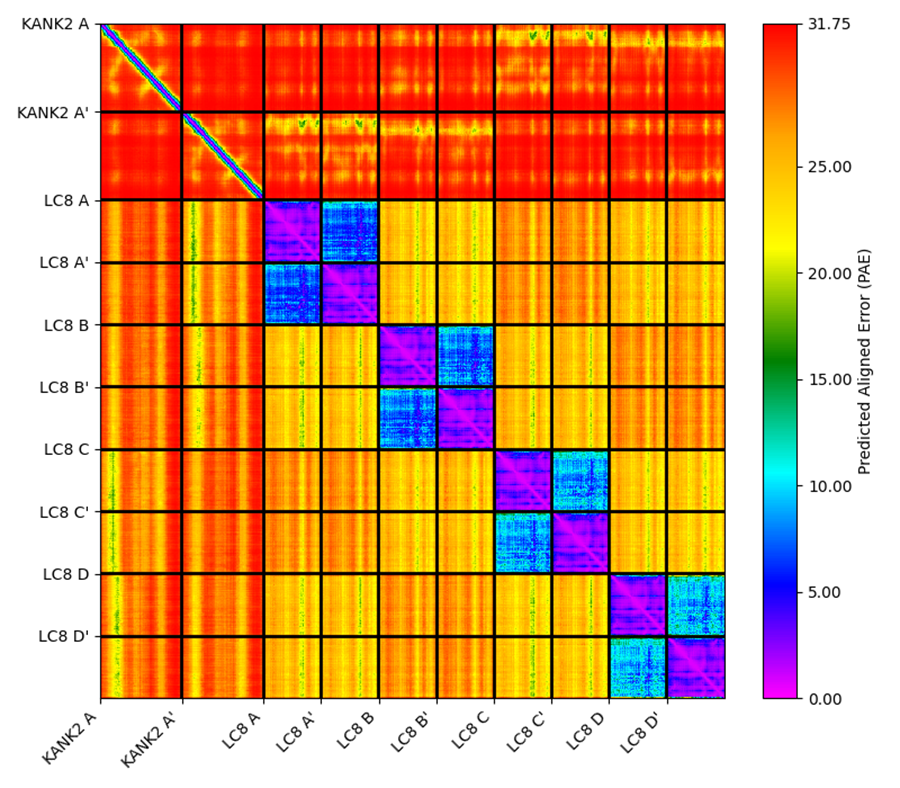

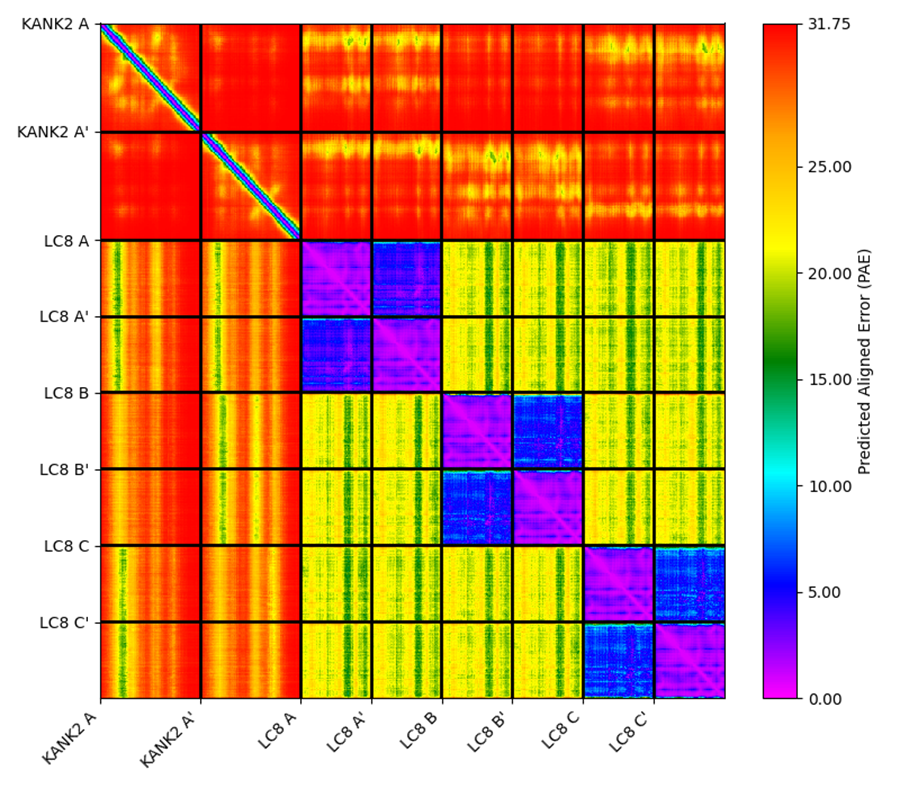

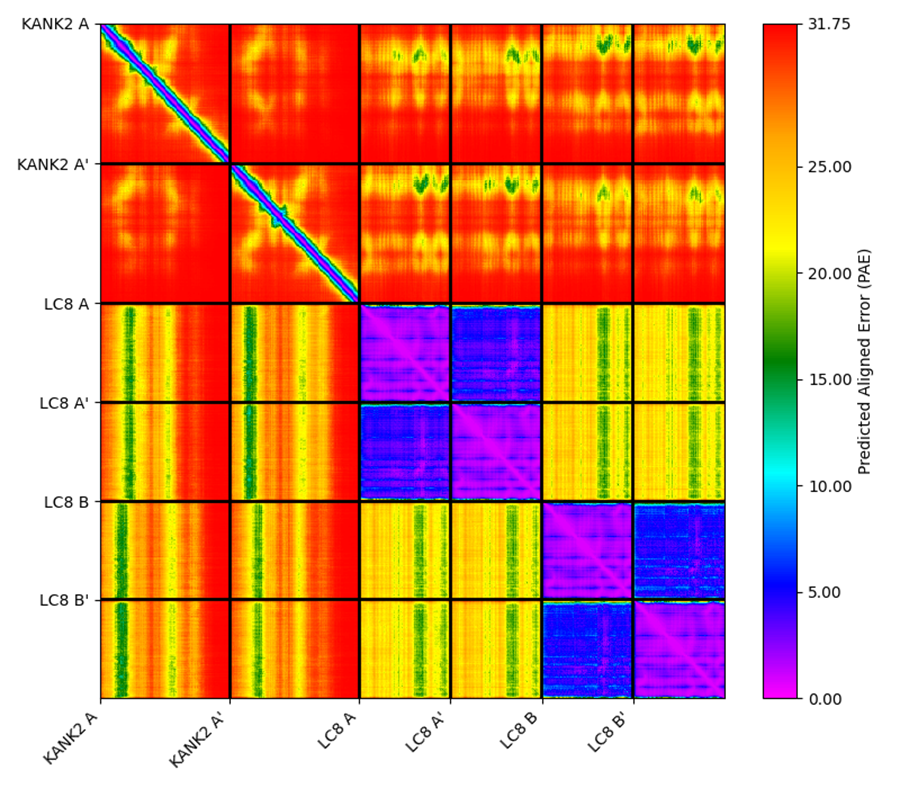

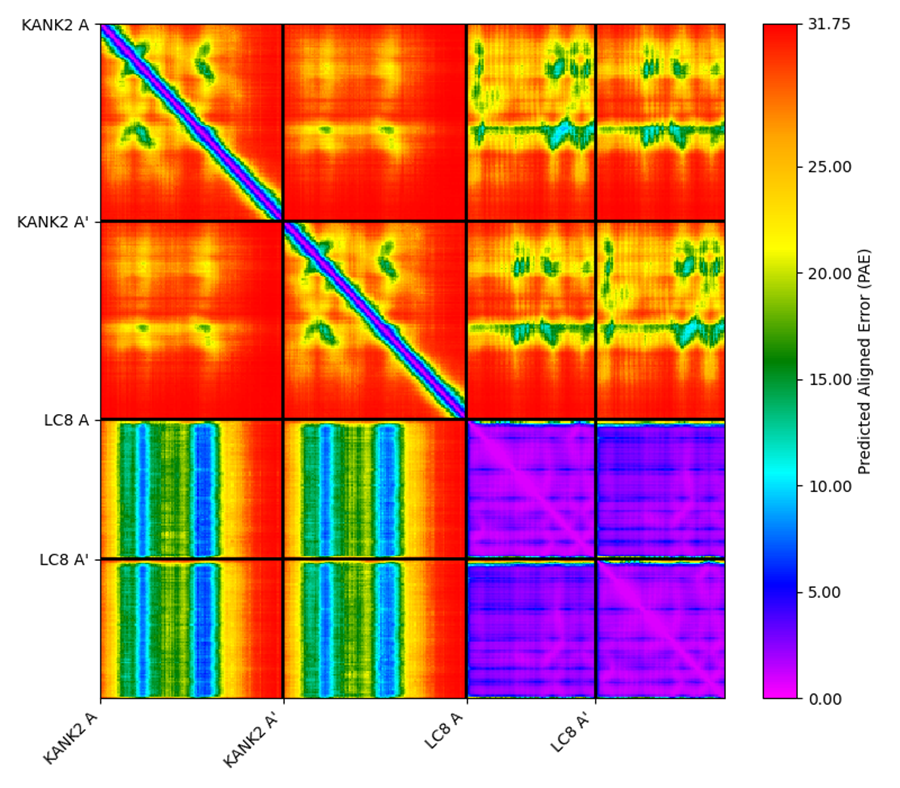

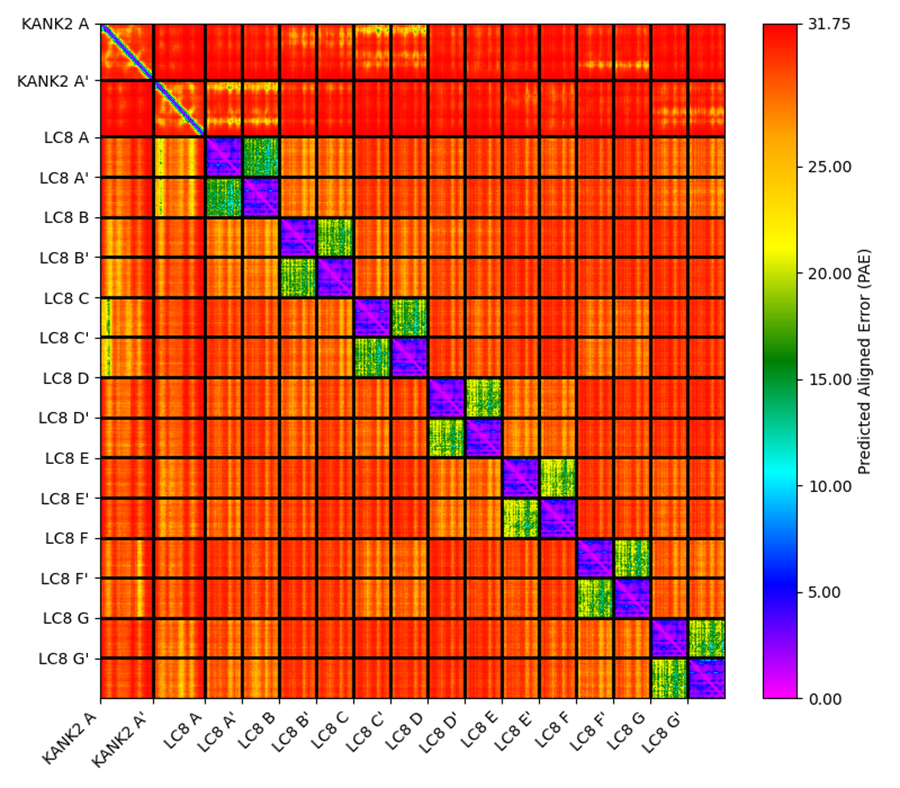

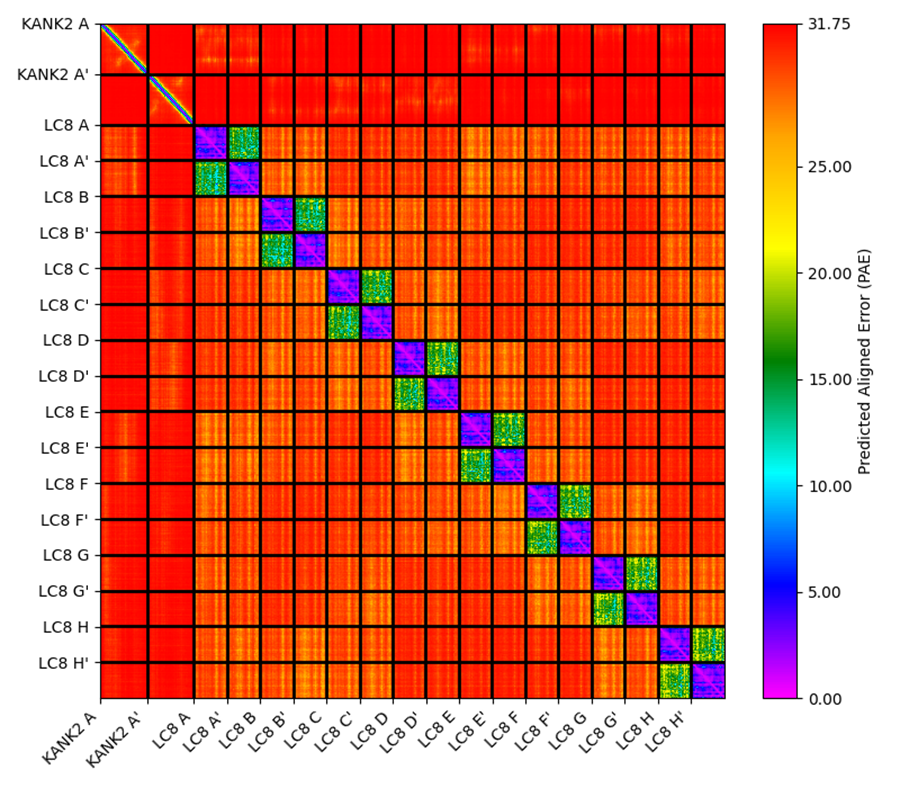

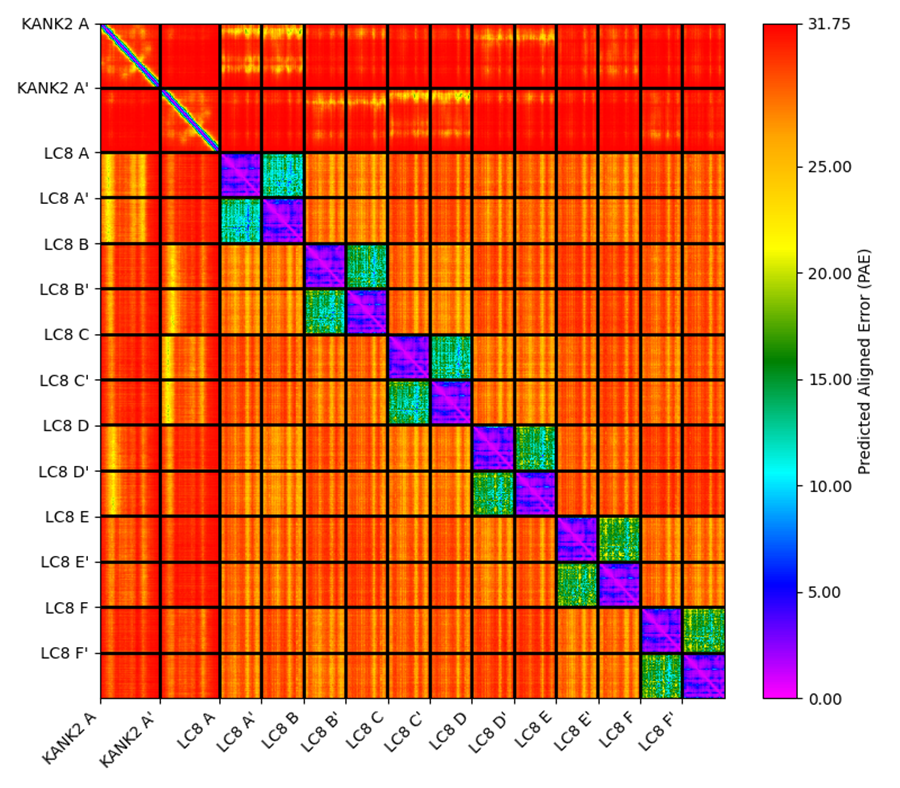

Fig. S5. PAE heatmaps of titration points in the AlphaFoldMultimer titration of LC8 with KANK2_313-437_. Consistent with experiments, none of the titration points are confident in KANK2/LC8 interactions.

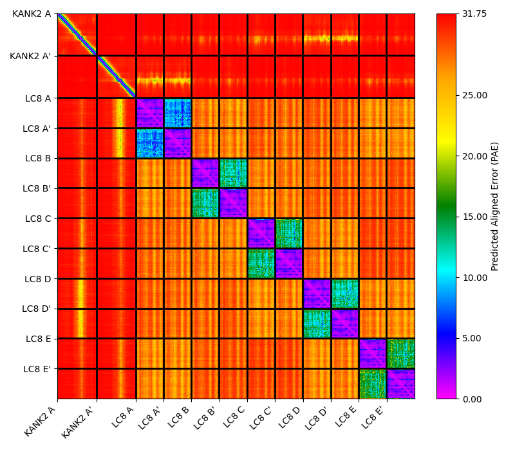

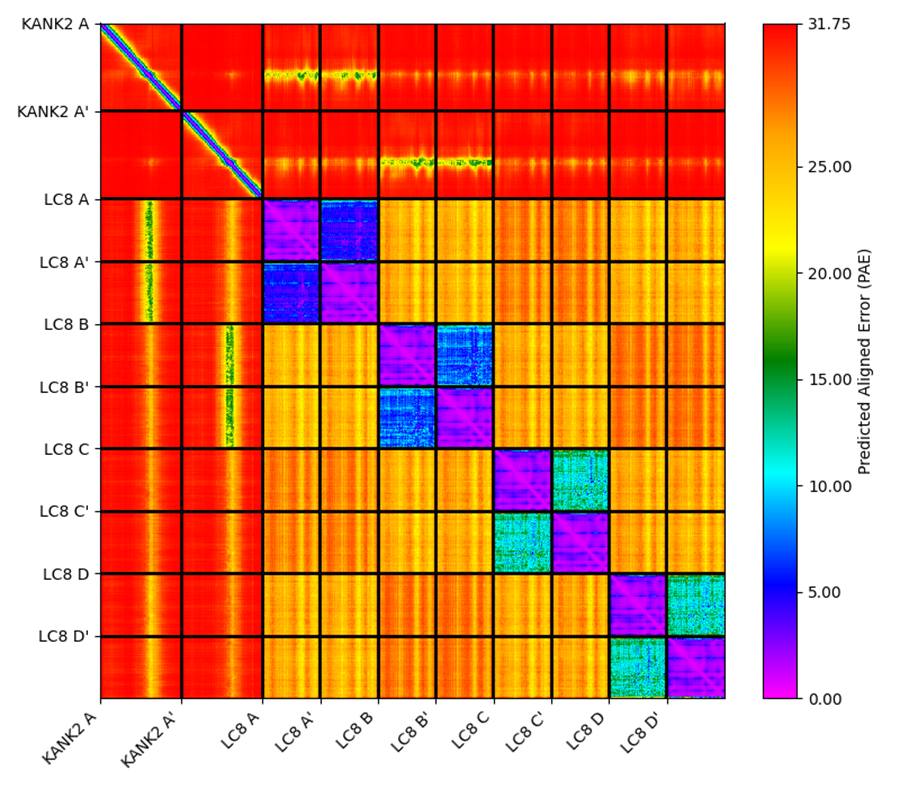

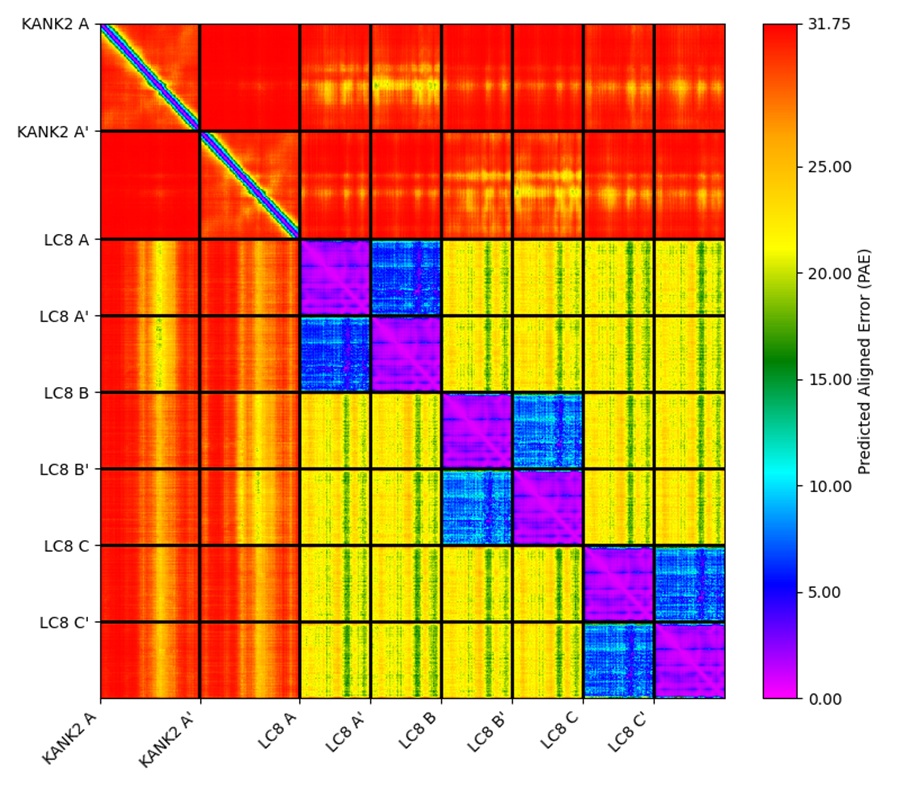

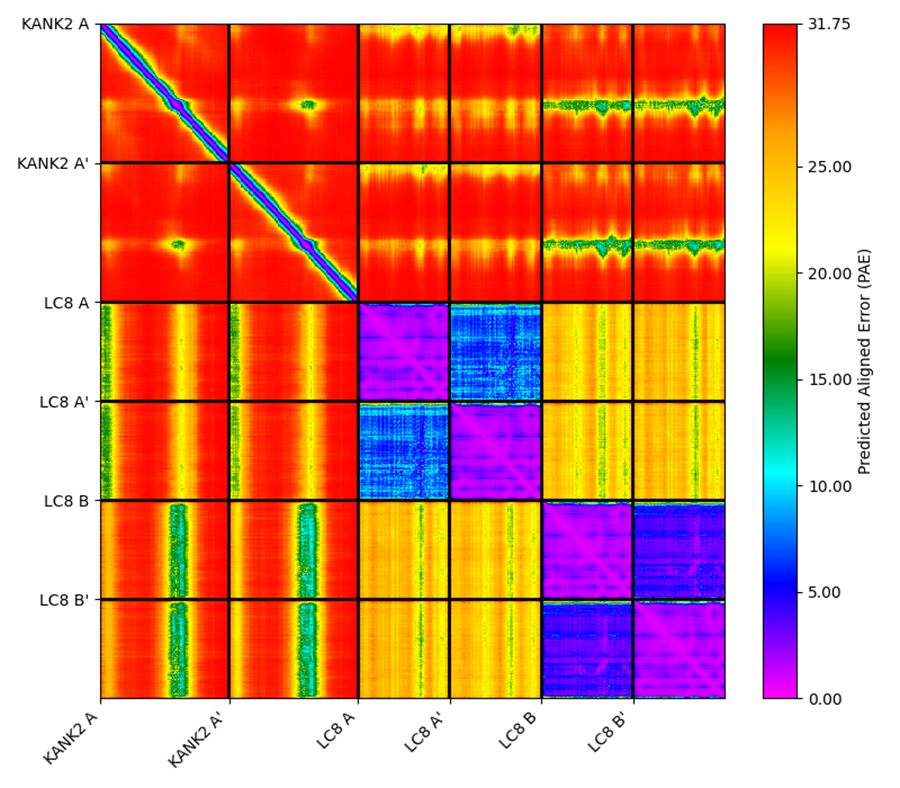

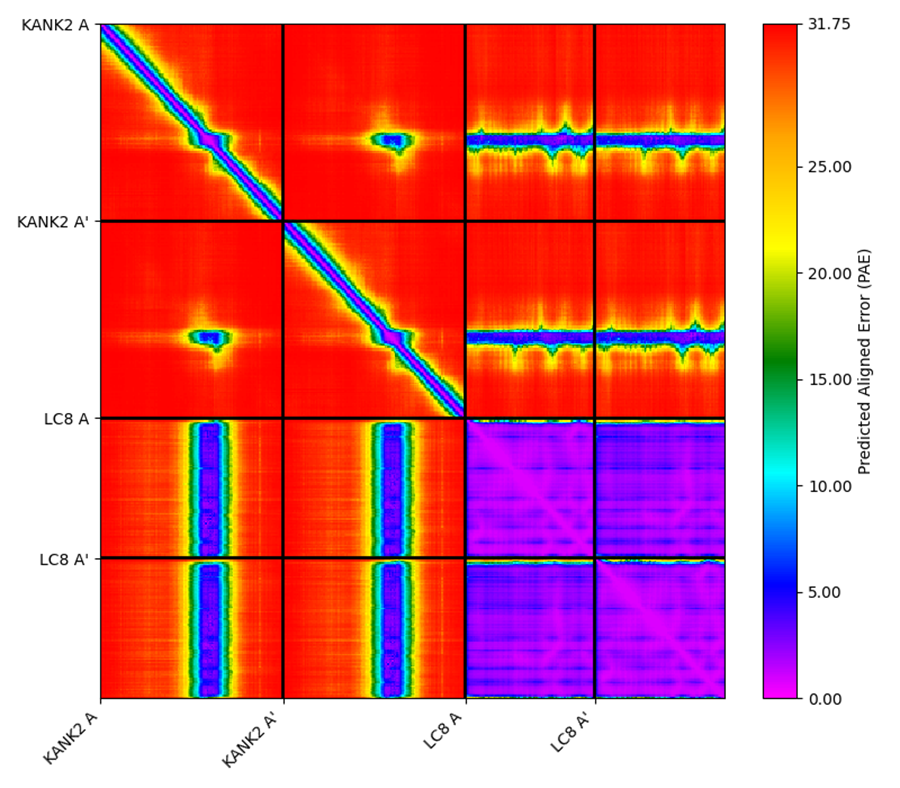

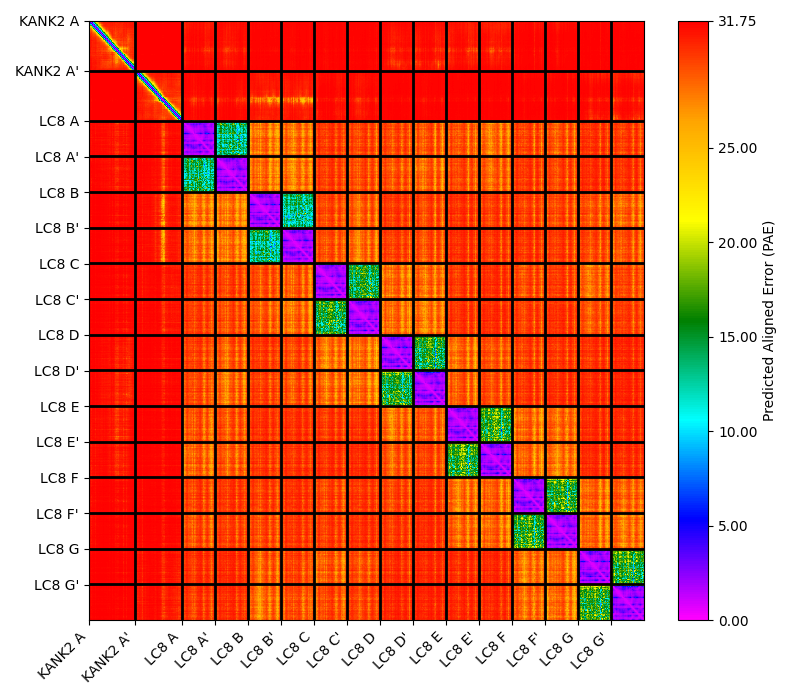

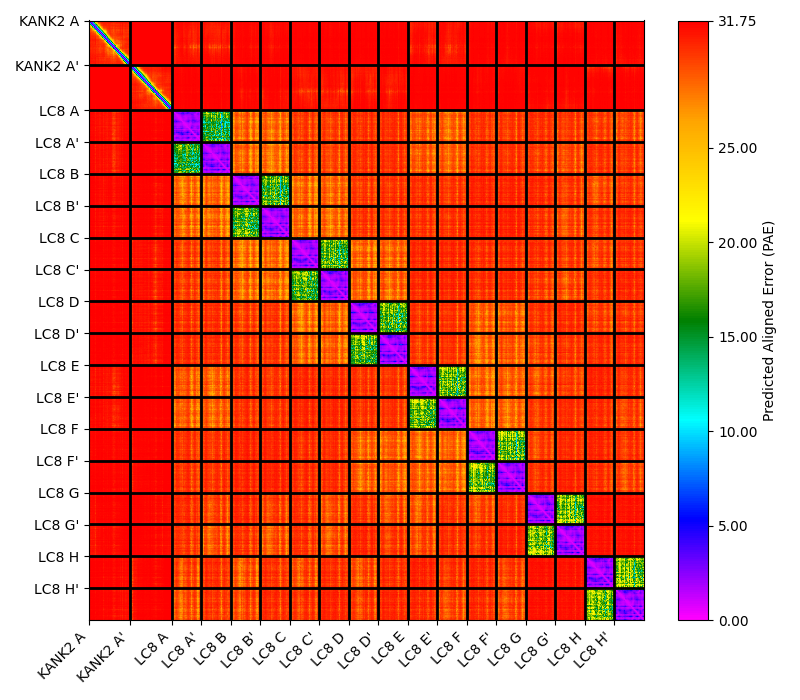

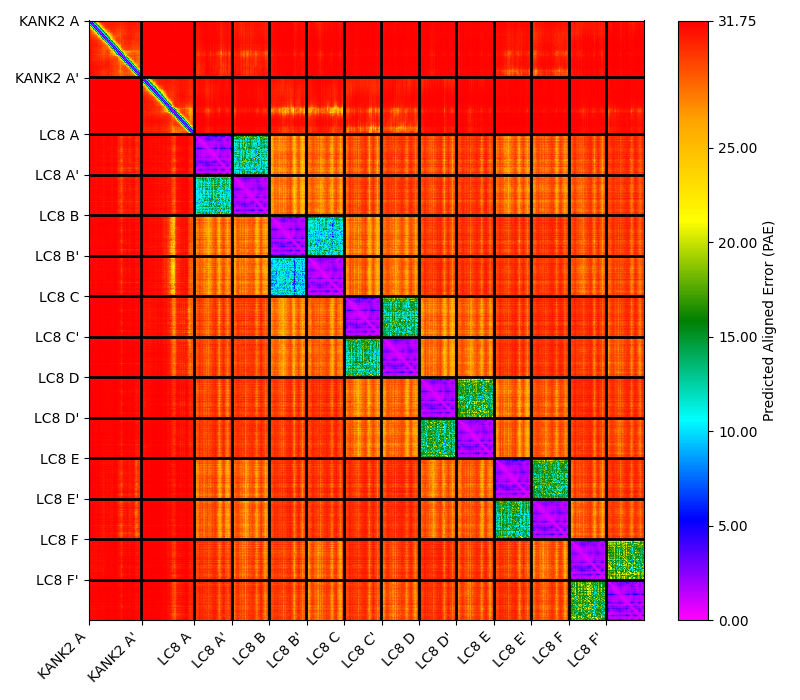

Fig. S6. PAE heatmaps of titration points in the AlphaFoldMultimer titration of LC8 with KANK2_427-552_. The first titration point suggests a potential binding site, which is a false positive, but the low confidence in KANK2/LC8 interactions at every other point is consistent with experiments.

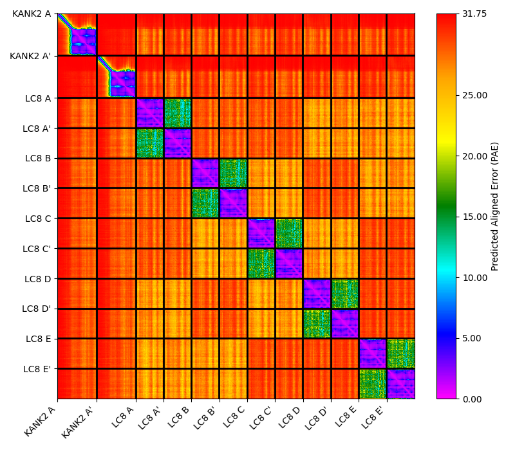

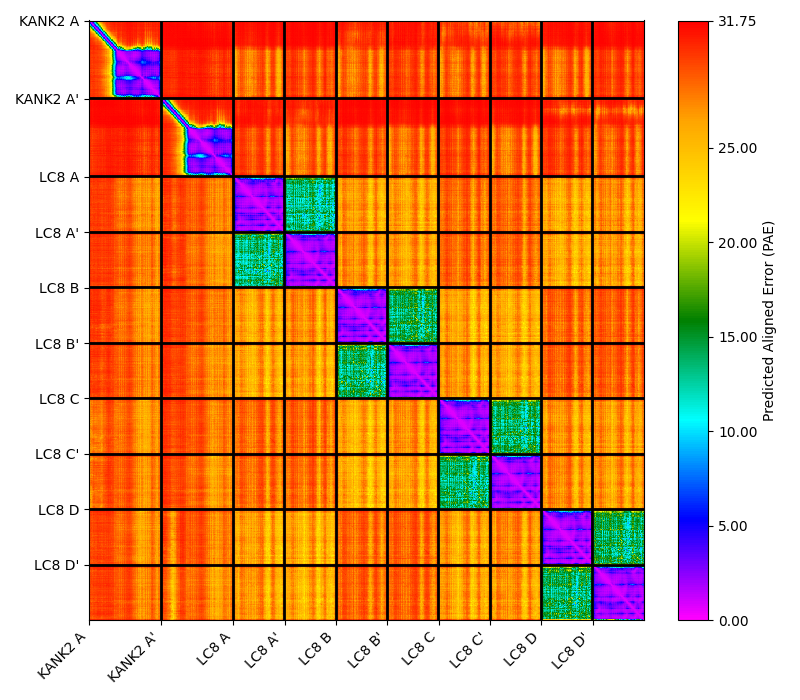

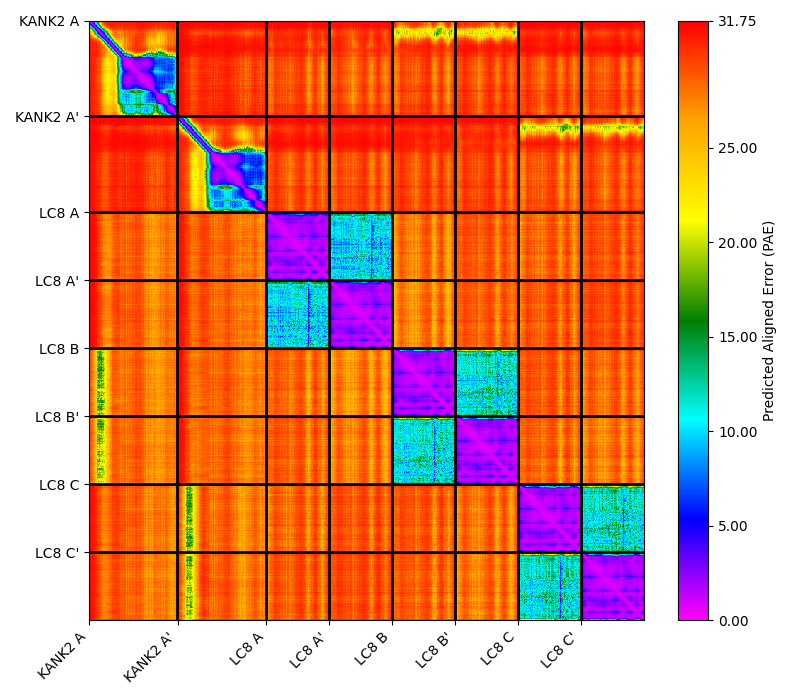

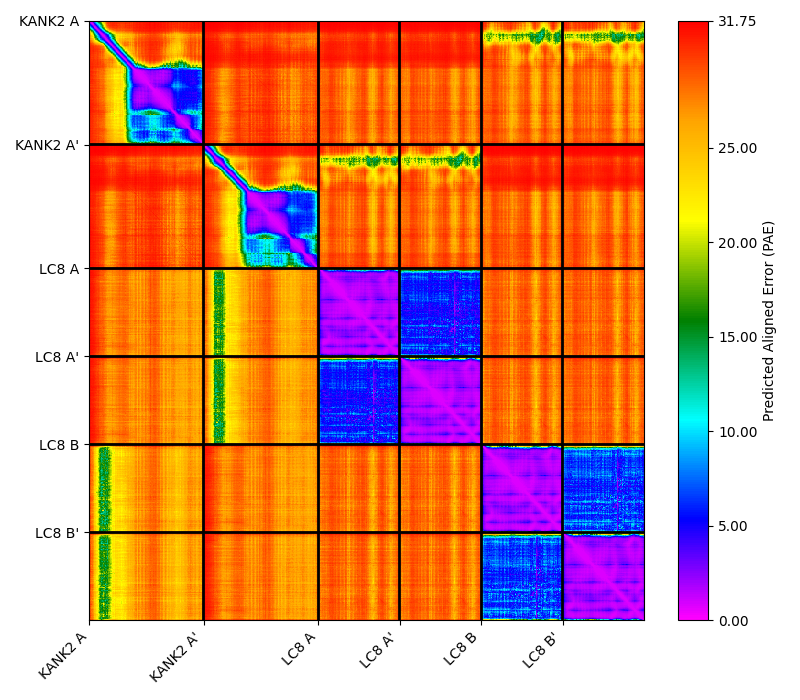

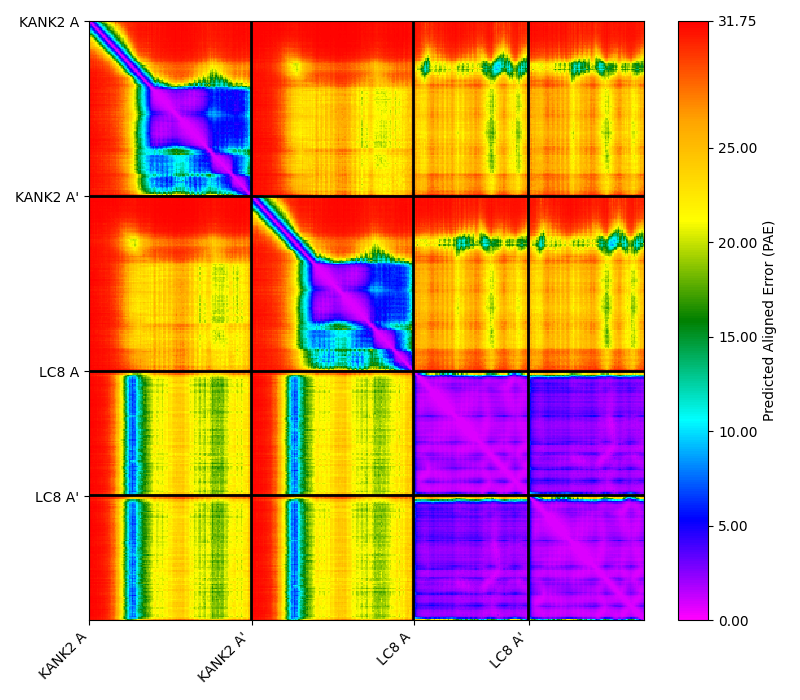

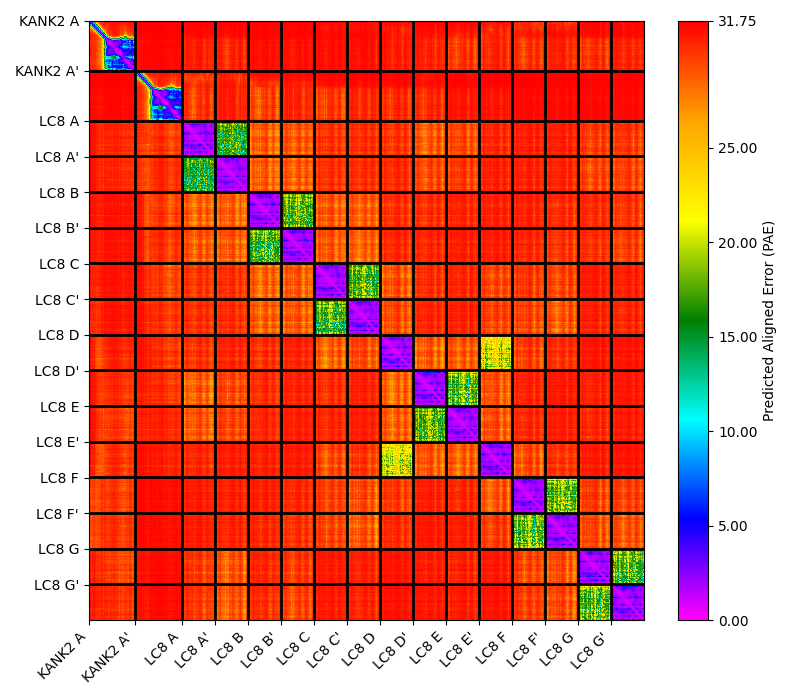

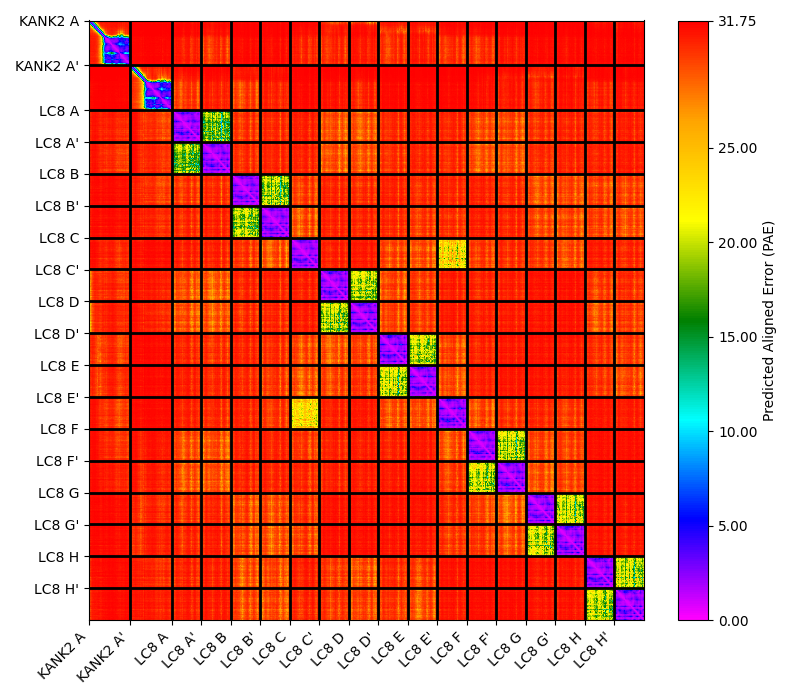

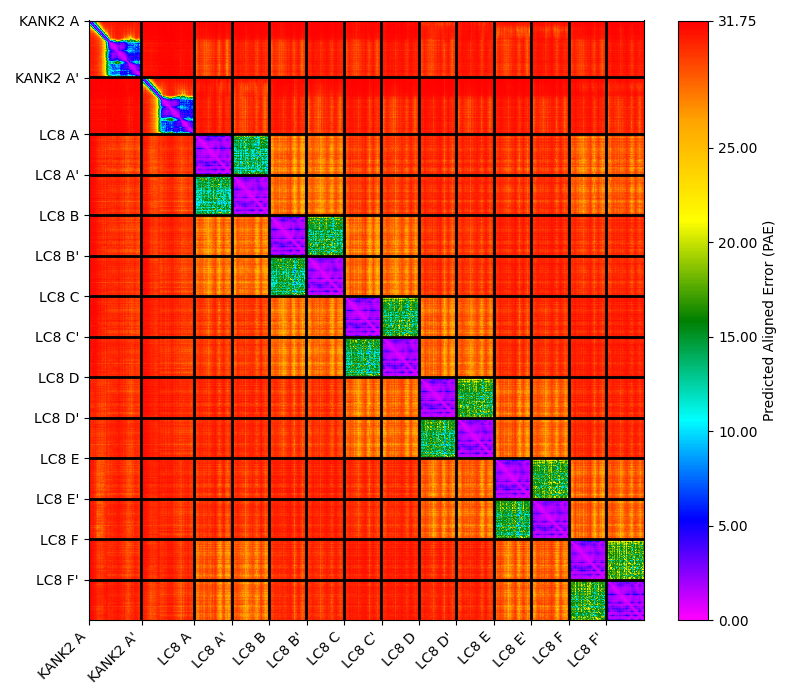

Fig. S7. PAE heatmaps of titration points in the AlphaFoldMultimer titration of LC8 with KANK2_542-666_. The C-terminus of the KANK2 construct suggests a bundle of intramolecular alpha helices. As consistent with experiments which show no binding, no titration point is confident in KANK2/LC8 interactions.

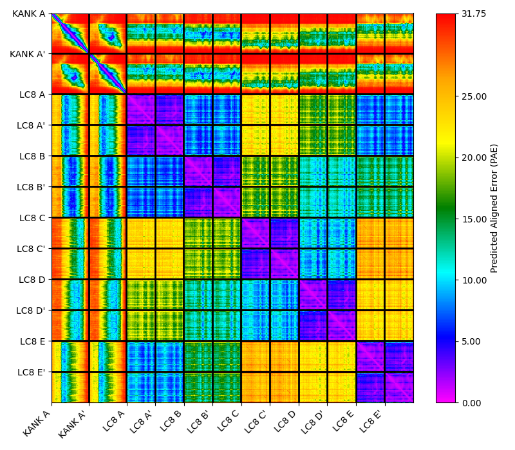

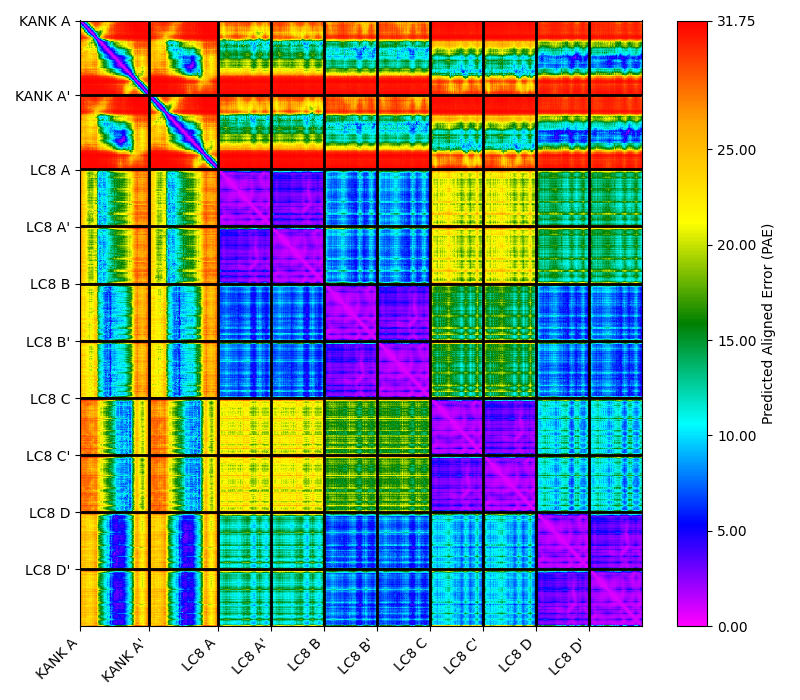

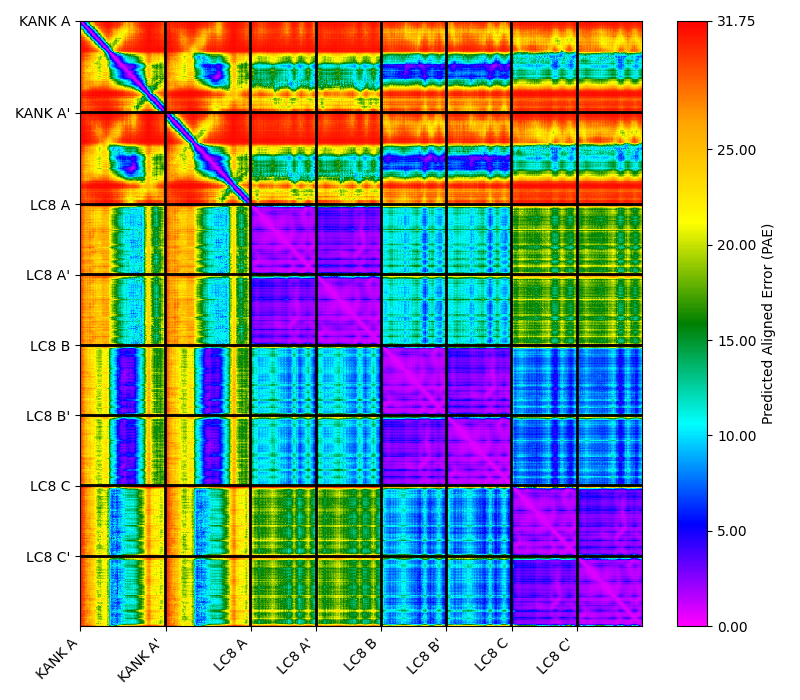

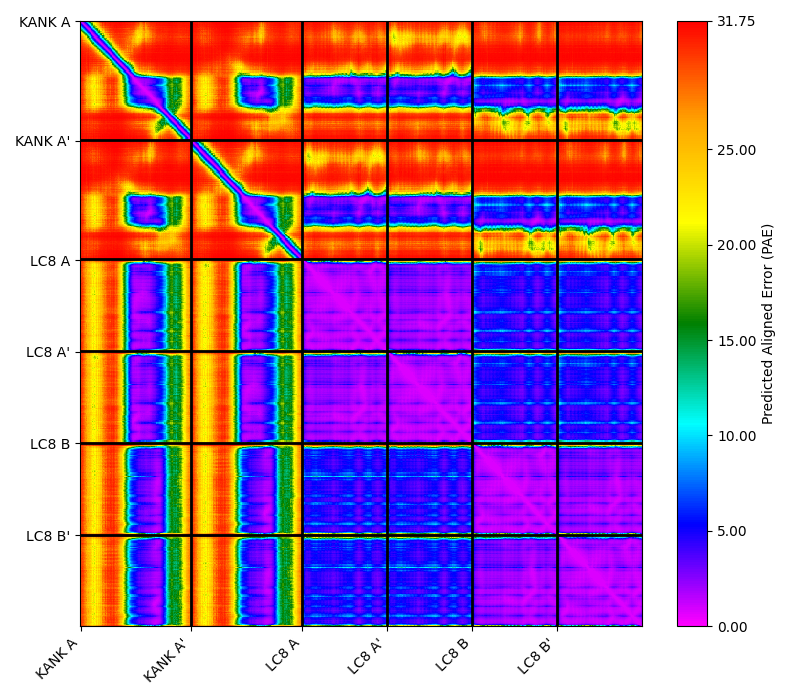

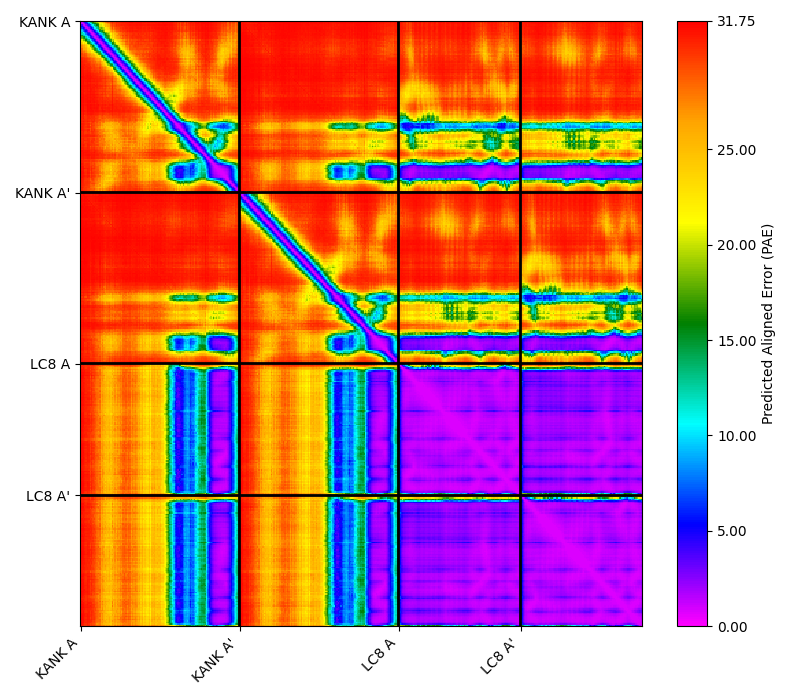

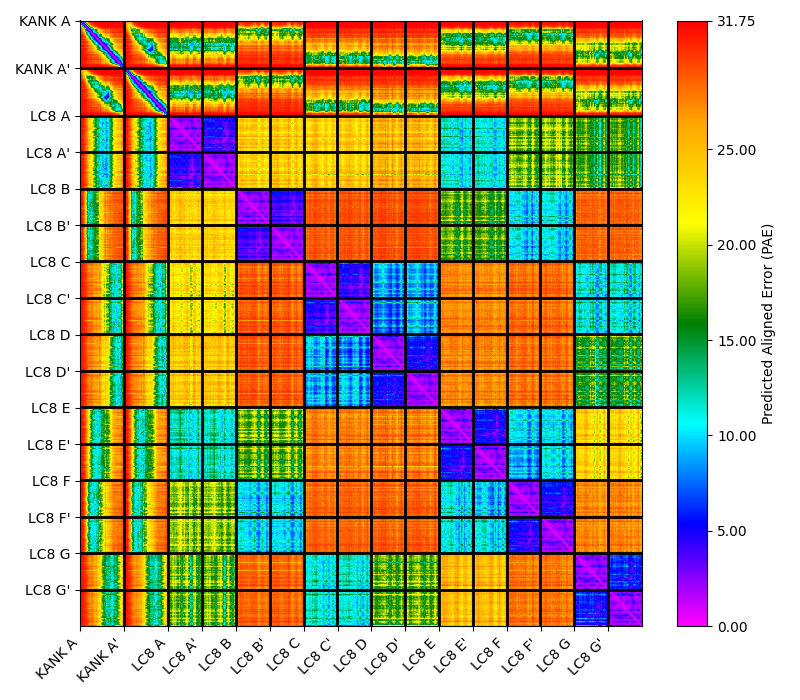

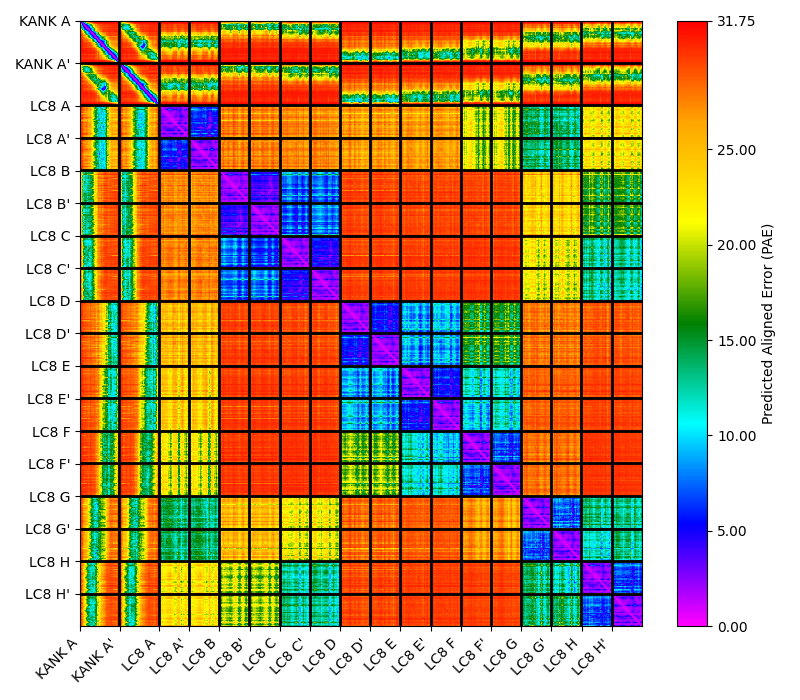

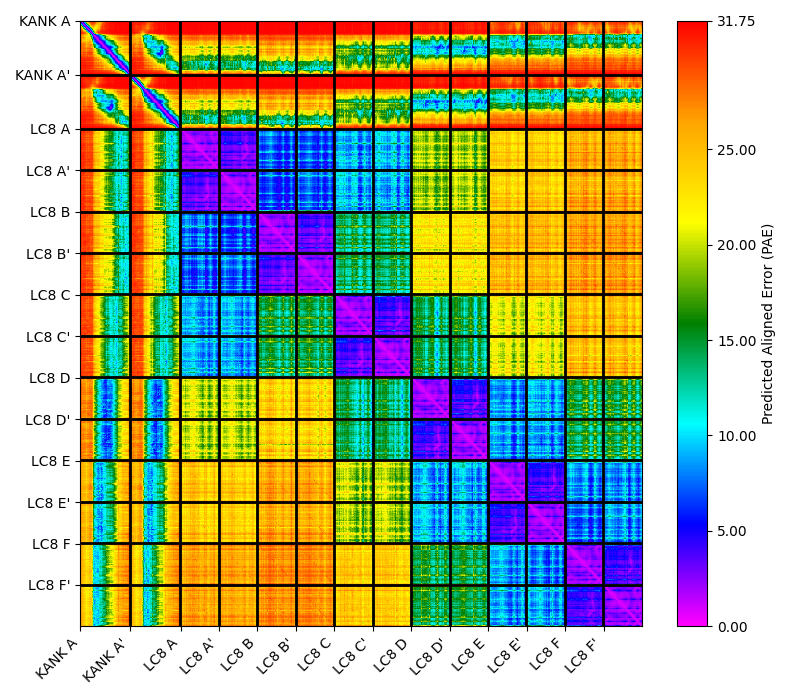

Fig. S8. PAE heatmaps of titration points in the AlphaFoldMultimer titration of LC8 with KANK1_500-615_. Consistent with experiments, AlphaFold is confident in KANK1/LC8 interaction at all 8 titration points, and LC8 dimers bind to the entire length of the KANK1 construct.

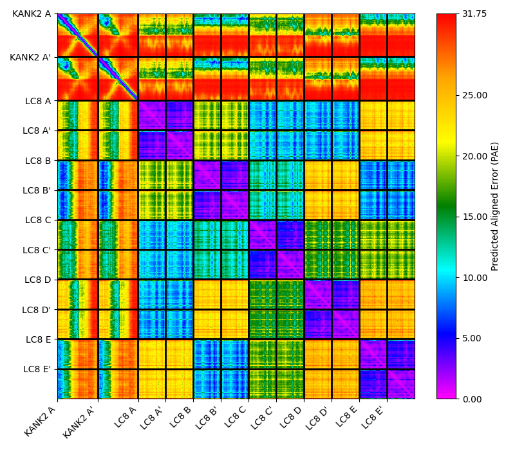

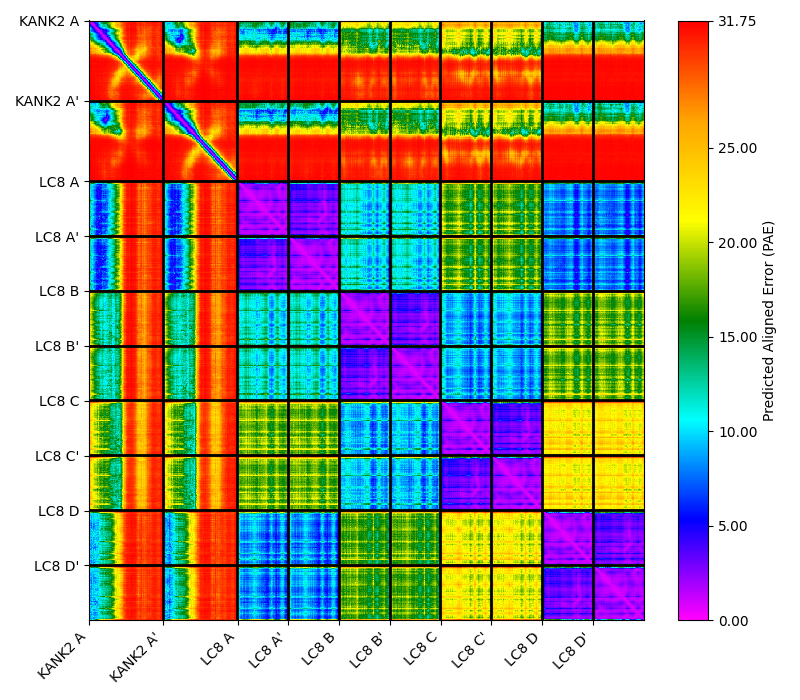

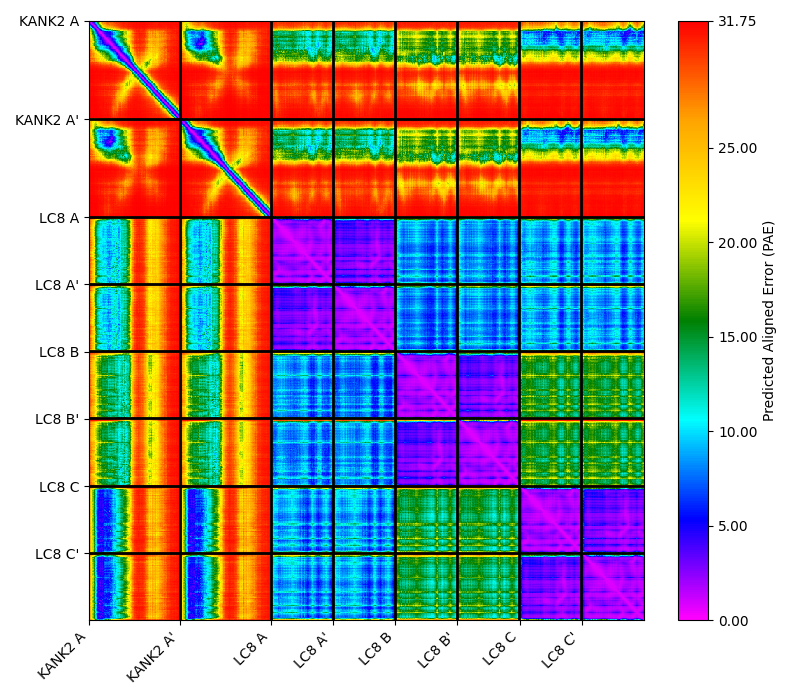

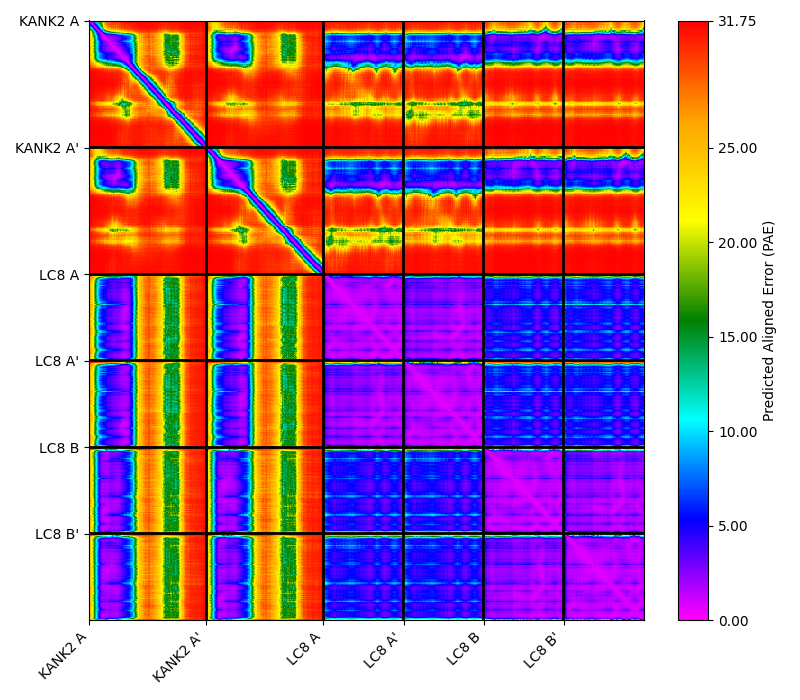

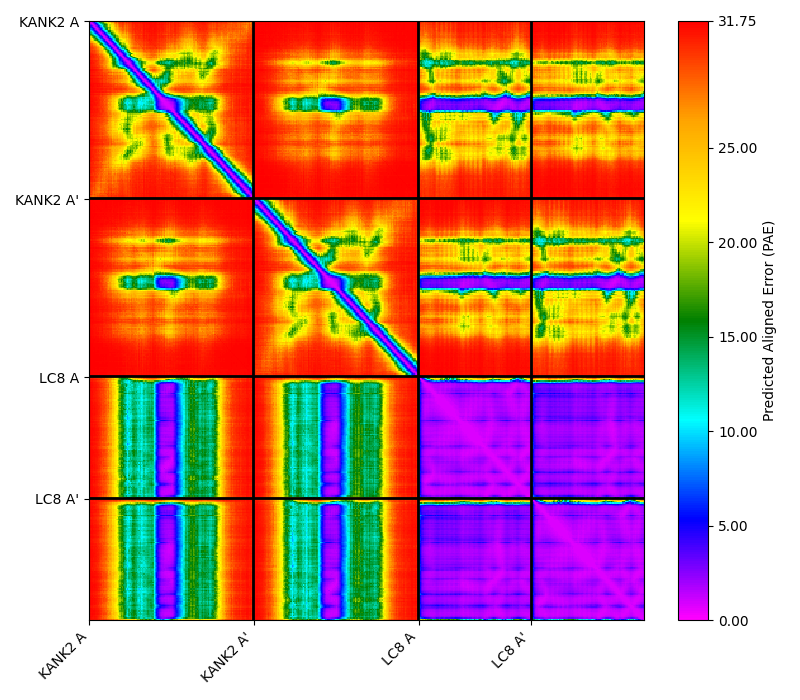

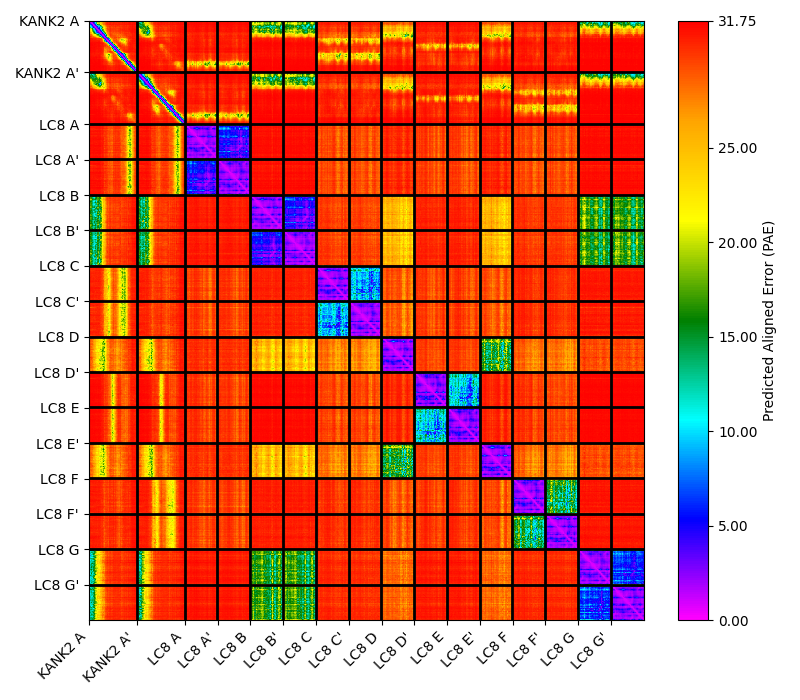

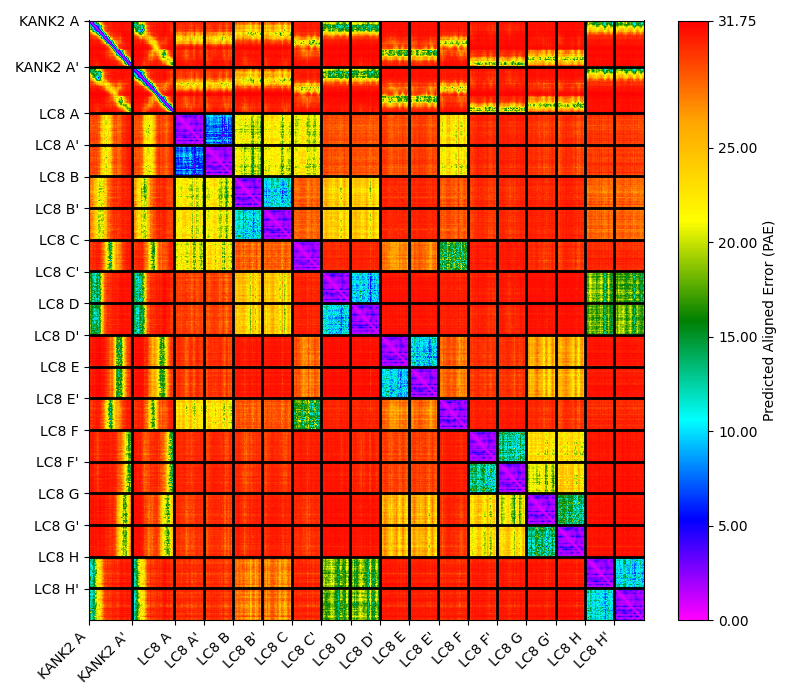

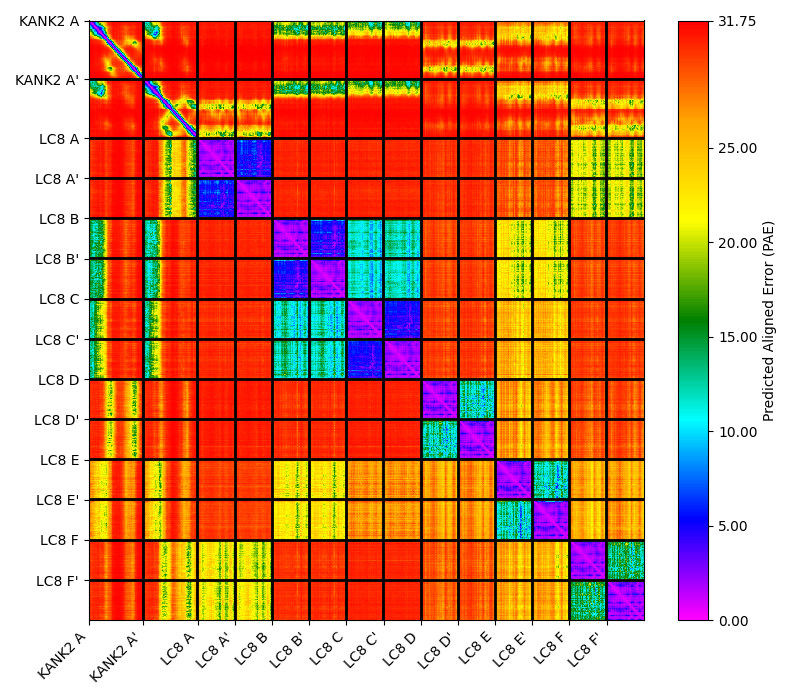

Fig. S9. PAE heatmaps of titration points in the AlphaFoldMultimer titration of LC8 with KANK1_541-670_. The reduced confidence in KANK1/LC8 interactions in this titration is consistent with the presence of the hinge in the rod which separates the two sections.

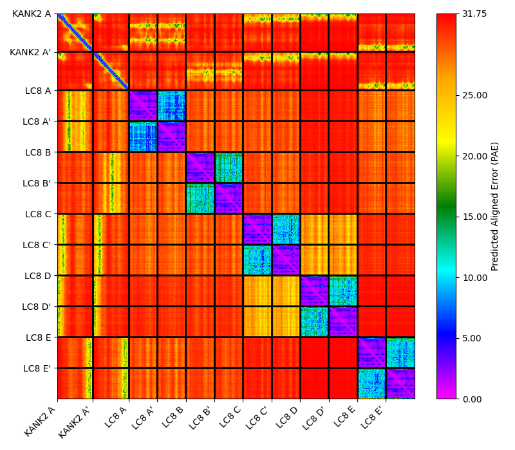

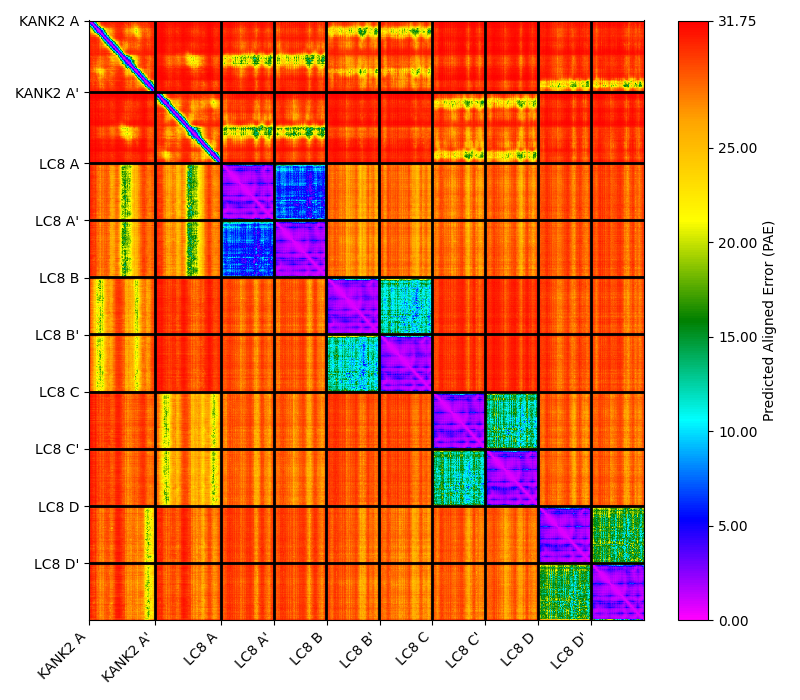

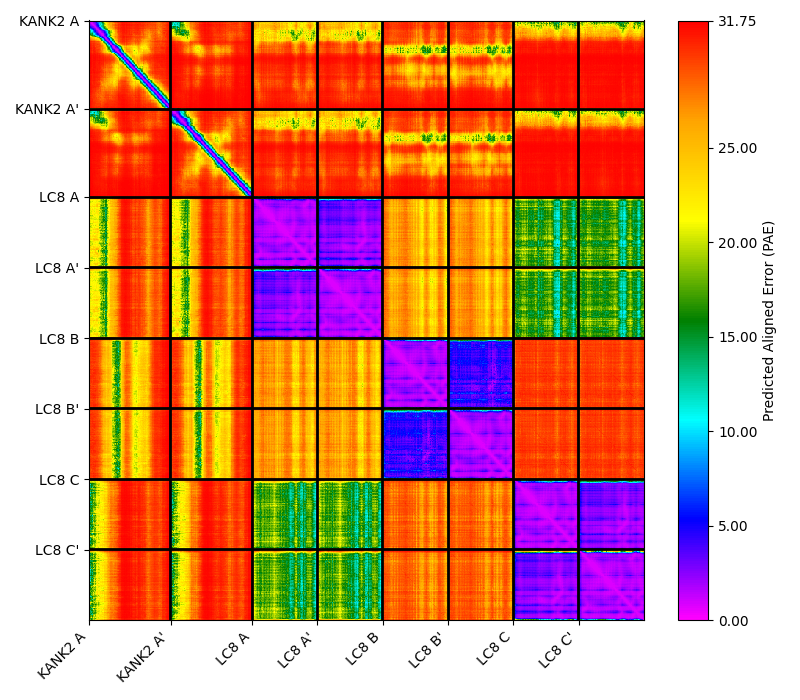

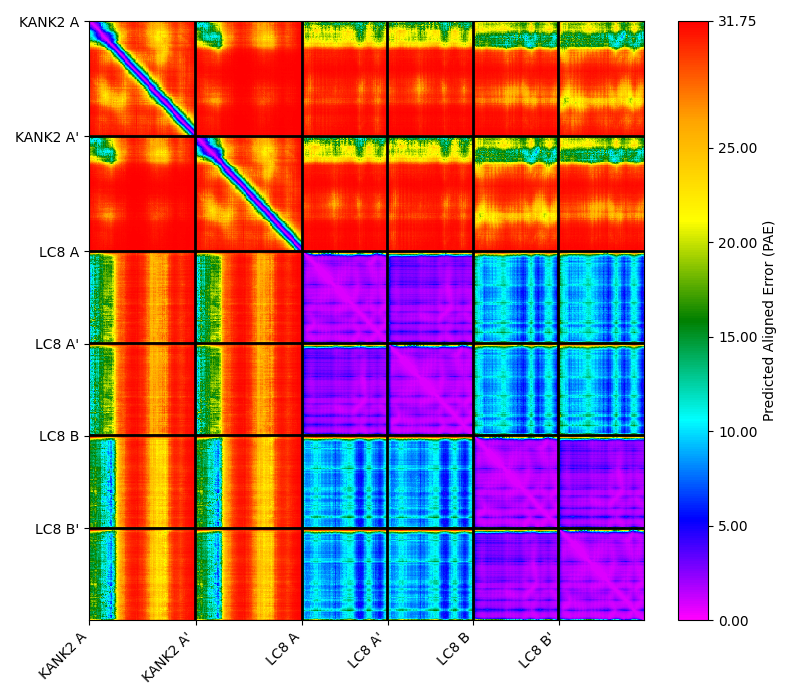

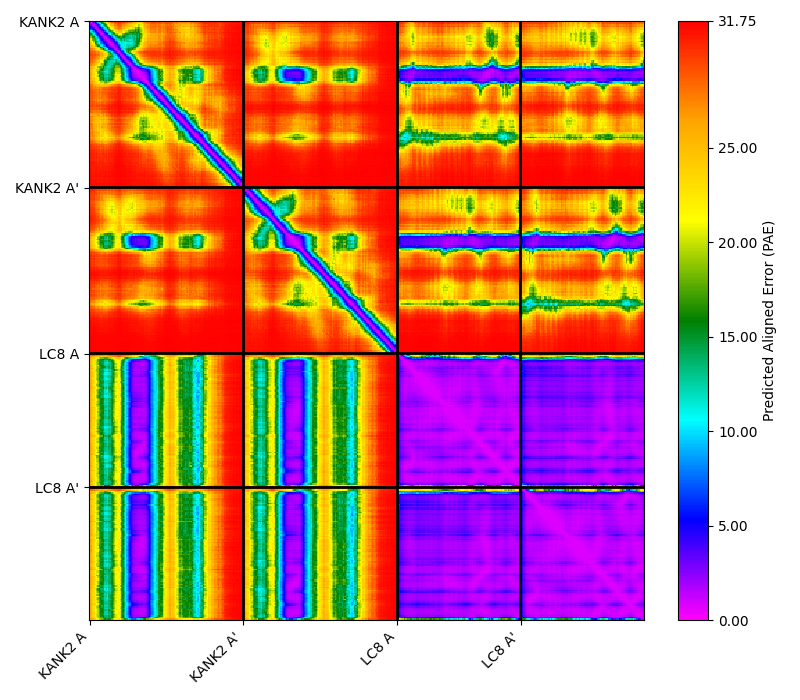

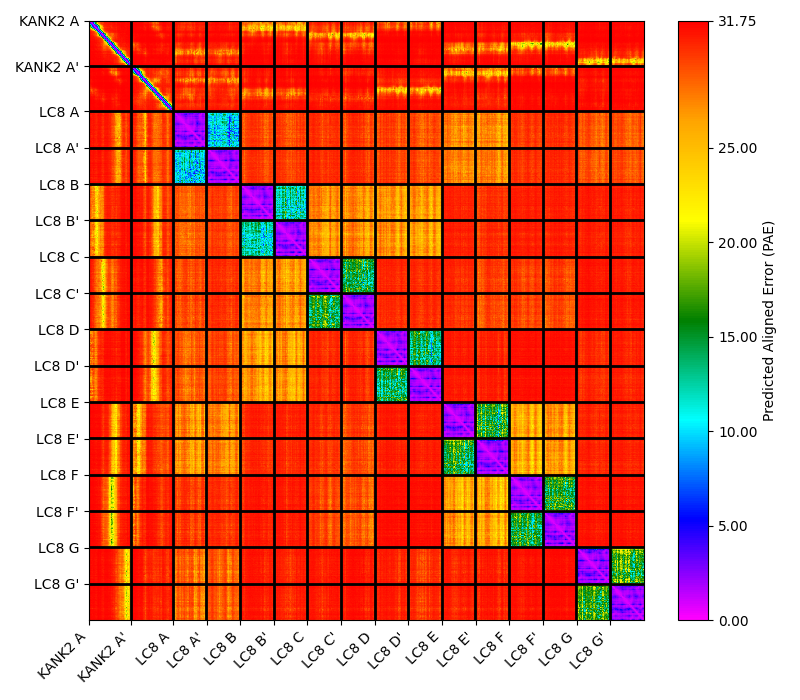

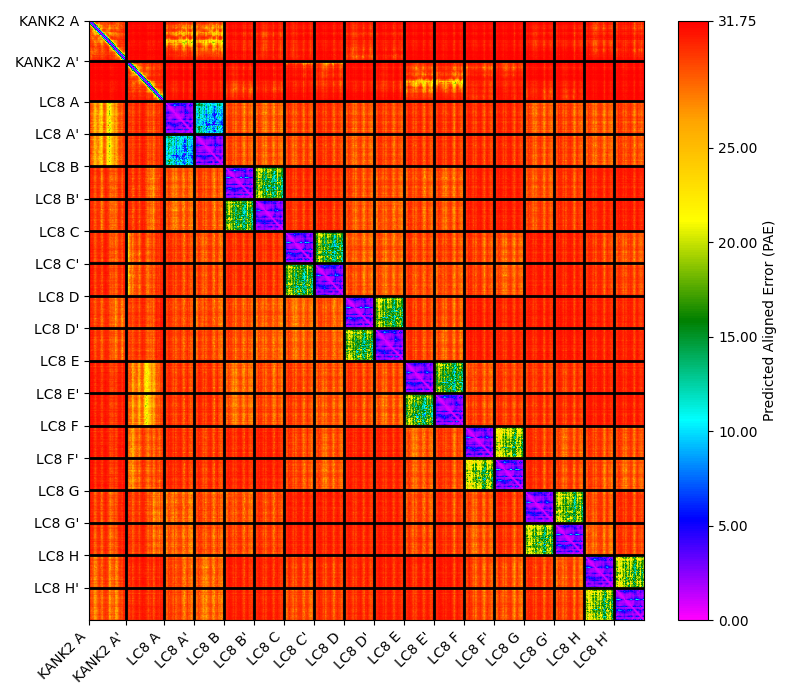

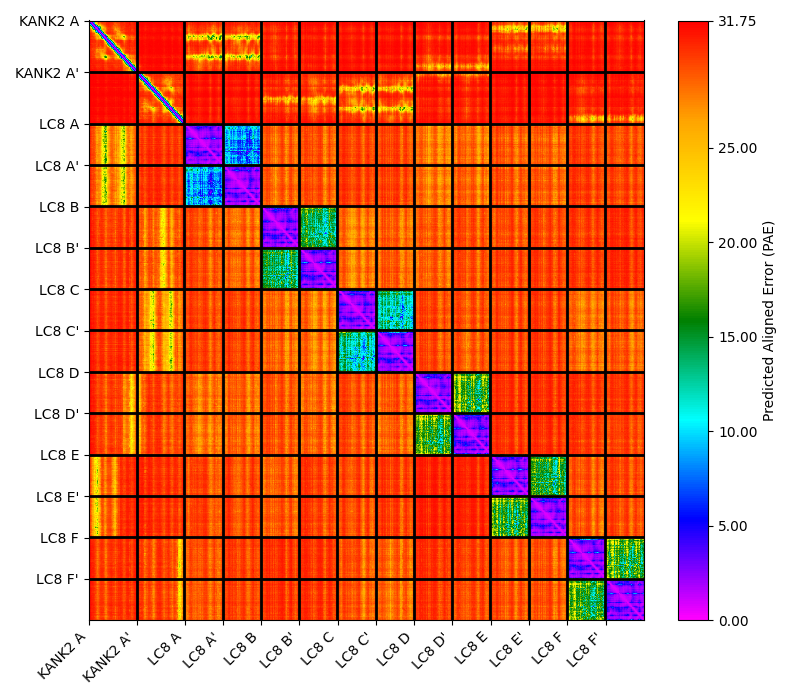

Fig. S10. PAE heatmaps of titration points in the AlphaFoldMultimer titration of LC8 with KANK1_700-810_. AlphaFold confidence in the KANK1/LC8 interactions indicates that only two LC8 dimers bind cooperatively in this construct, consistent with experiments at low concentrations which implicate 15 binding sites, corresponding to those in 500-720. However, at higher titration points (3-5), LC8 dimers still associate at positions predicted by AlphaFold-Pred, indicated by narrow regions of increased confidence. This behavior is consistent with weak binding sites that are not stabilized by cooperativity and rod formation.

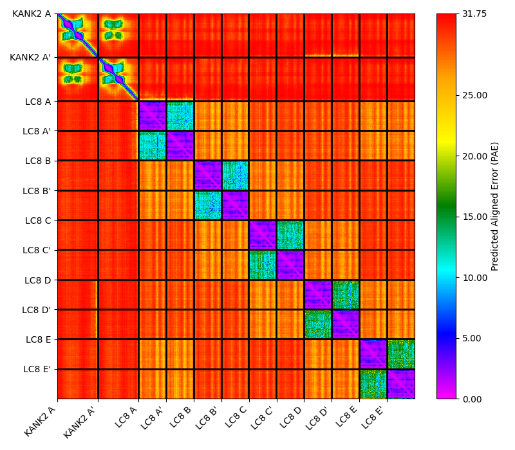

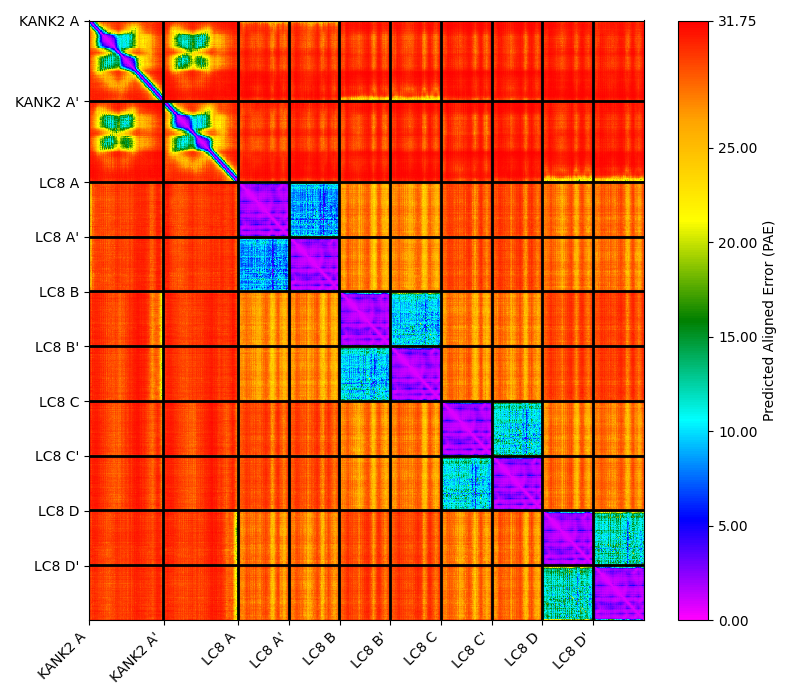

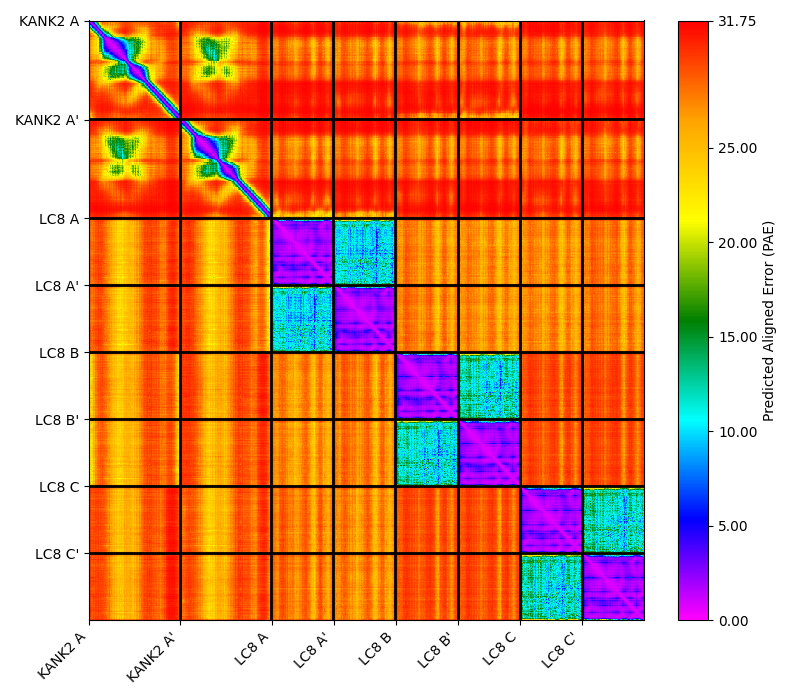

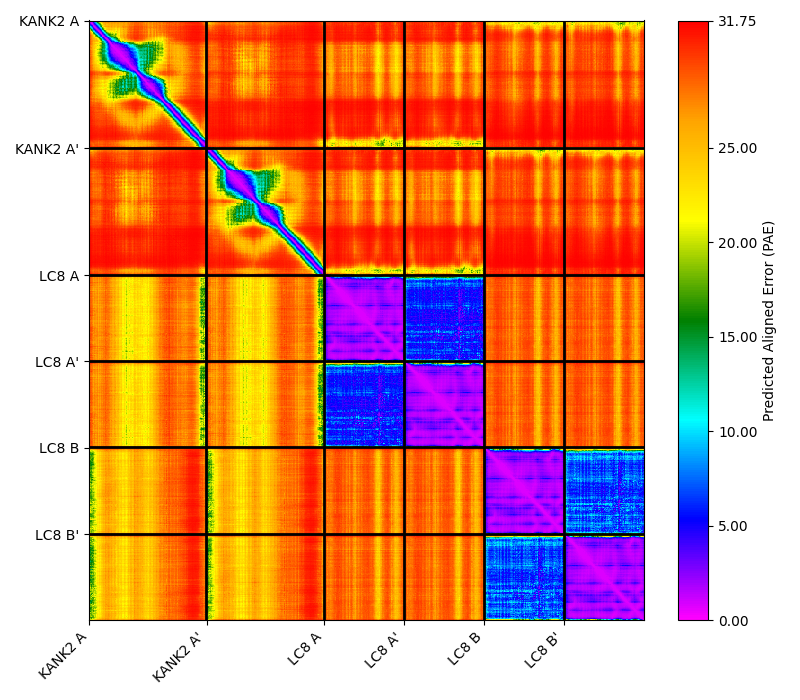

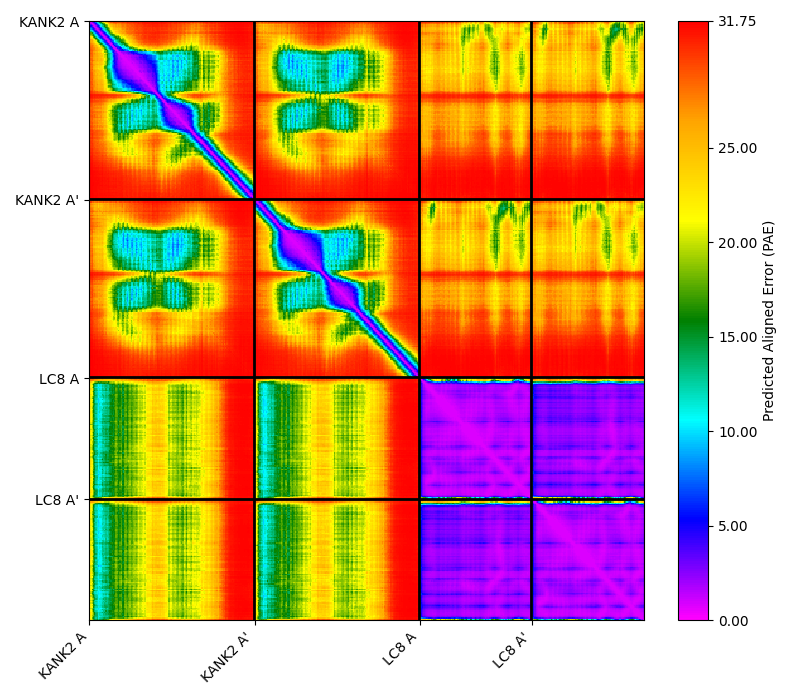

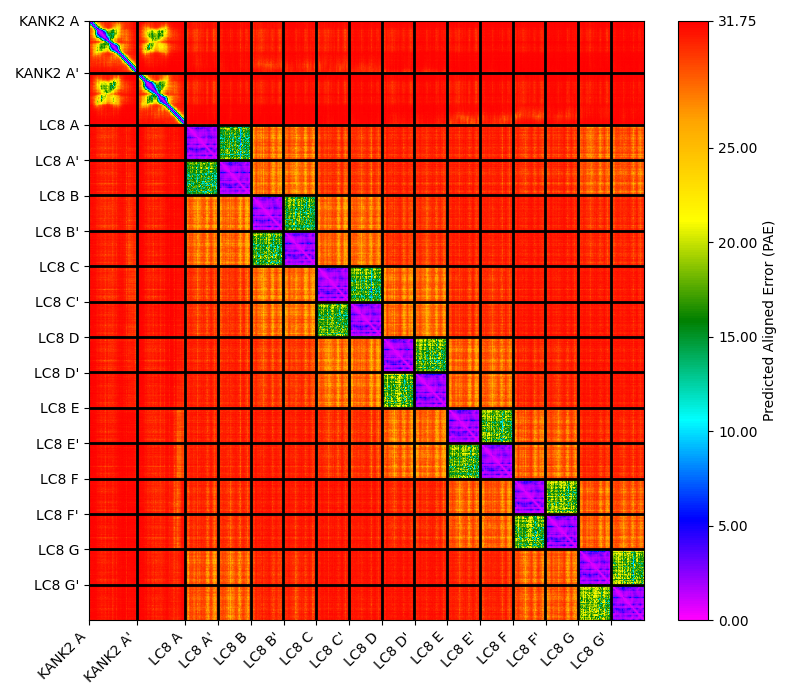

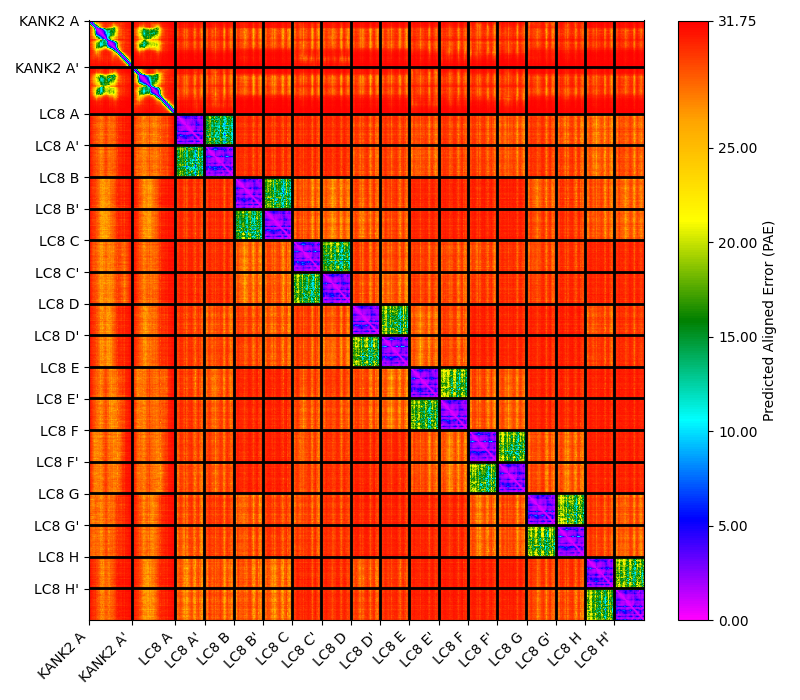

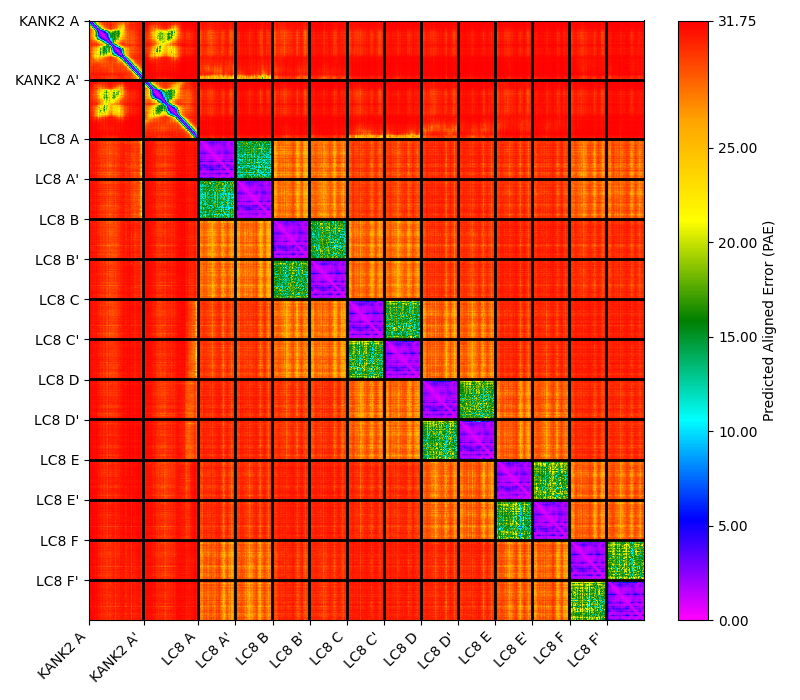

Fig. S11. PAE heatmaps of titration points in the AlphaFoldMultimer titration of LC8 with KANK1_800-930_. The KANK1/KANK1 interaction window suggests a mildly confident intermolecular 4-helix bundle. The lack of confidence in KANK1/LC8 interactions is consistent with pulldowns which show no interaction for this section of KANK1.

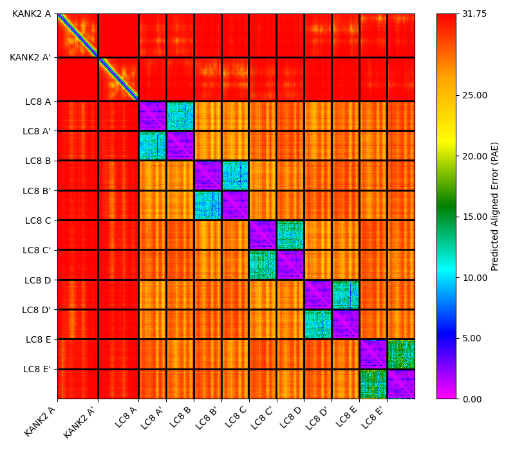

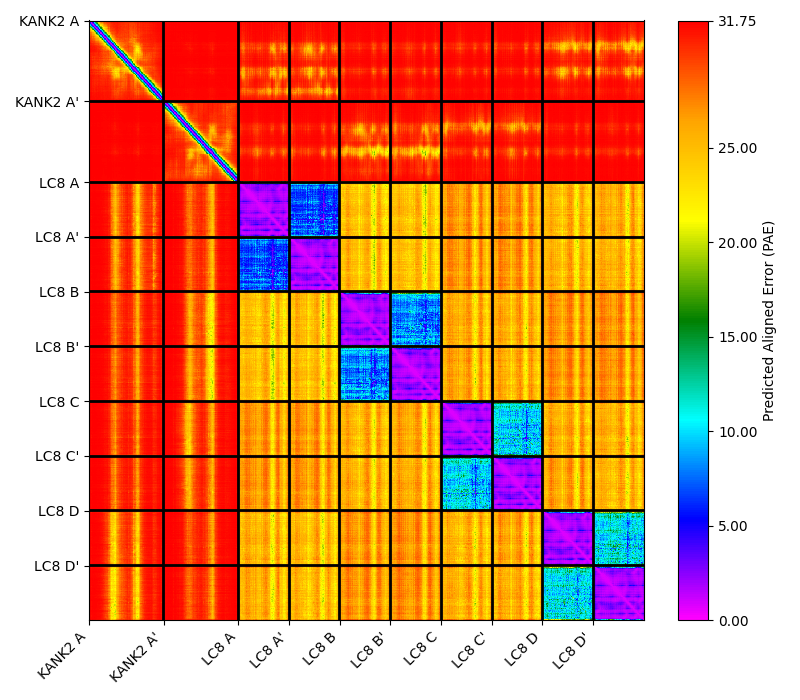

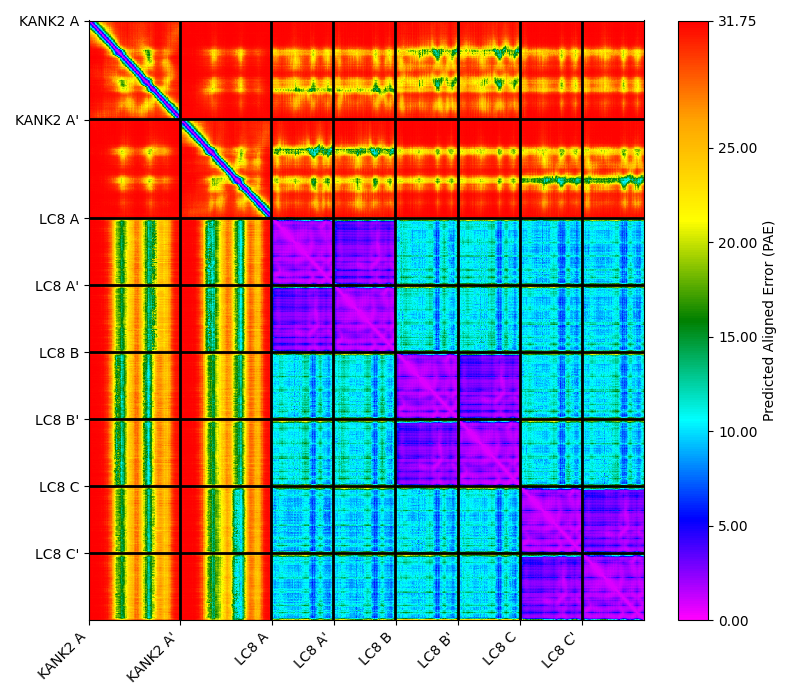

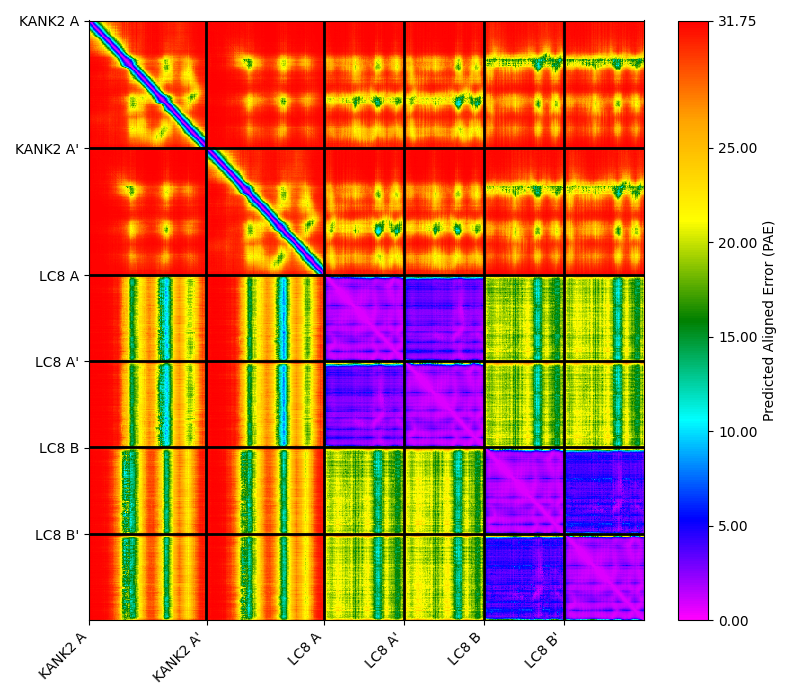

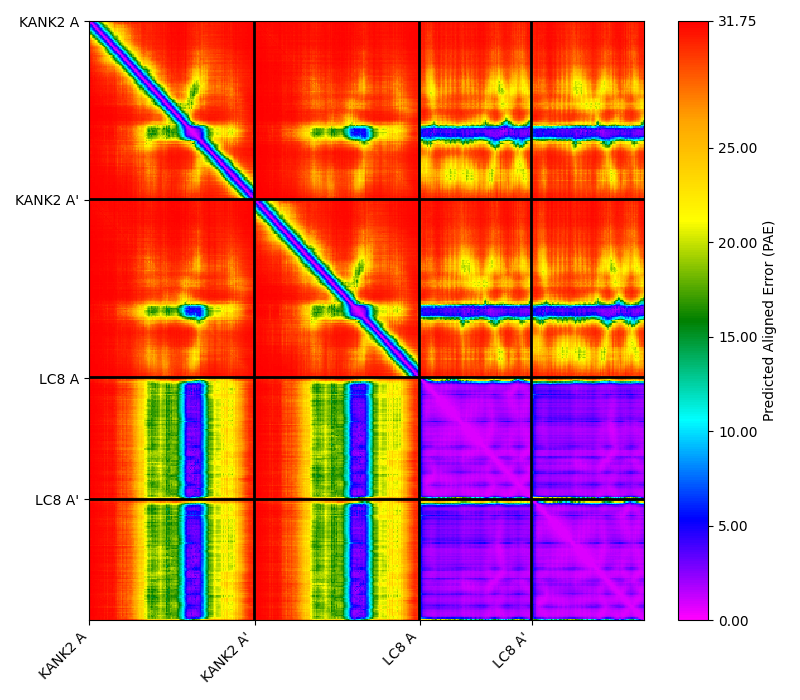

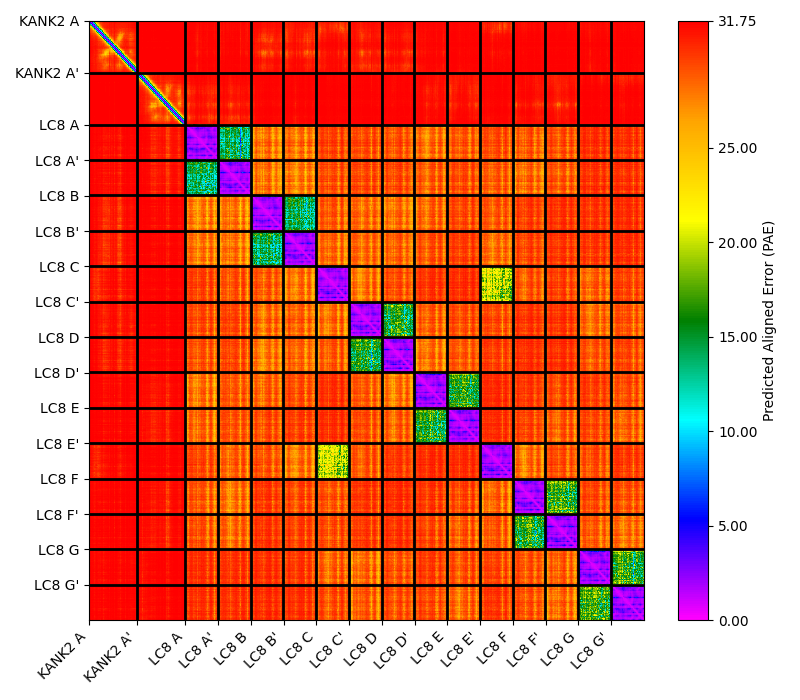

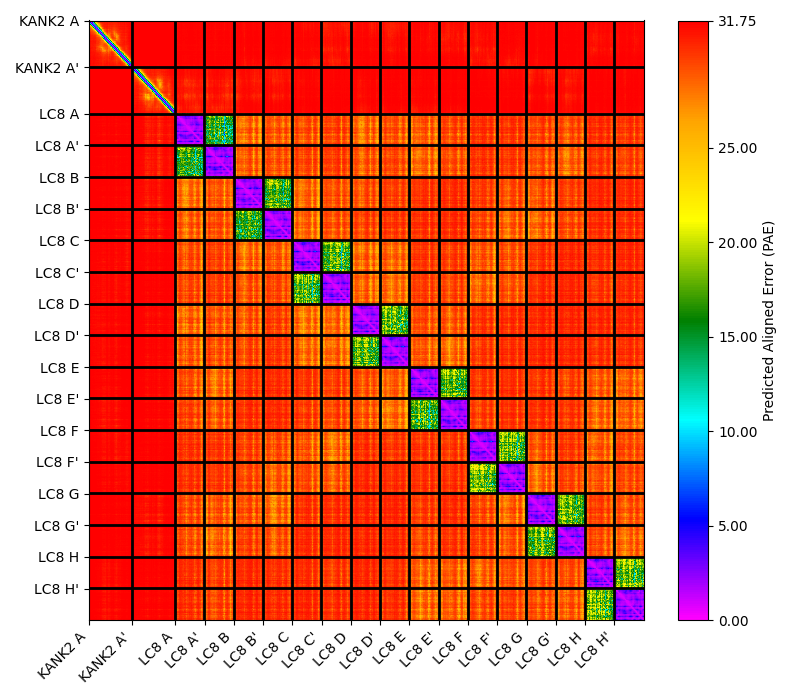

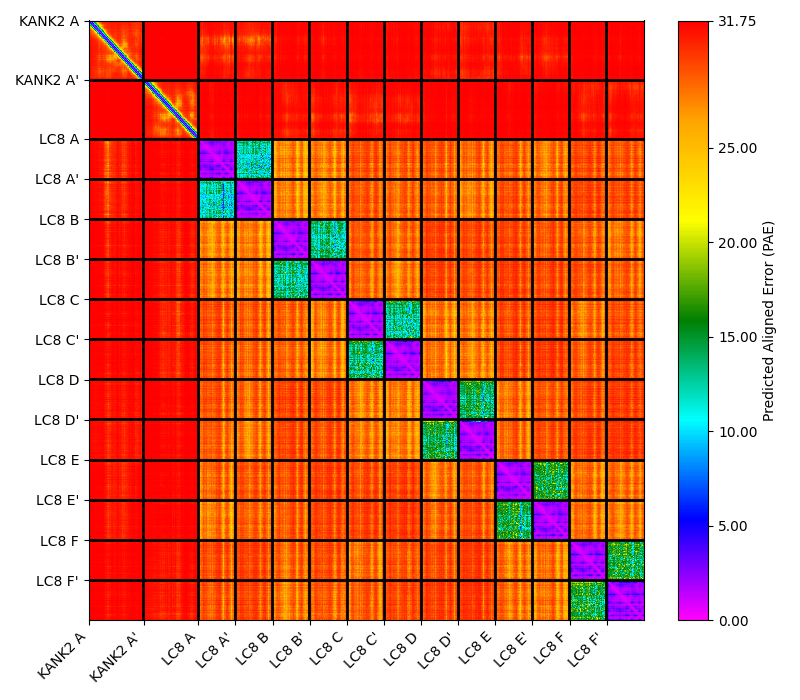

Fig. S12. PAE heatmaps of titration points in the AlphaFoldMultimer titration of LC8 with KANK1_915-1045_. The 1:1 ratio titration point is consistent with AlphaFold-Pred results that indicate one potential binding site in this region. However, according to pulldowns, this is a false positive prediction. The KANK1/LC8 interaction confidence of the rest of the titration is consistent with experiments which indicate no binding in this region.

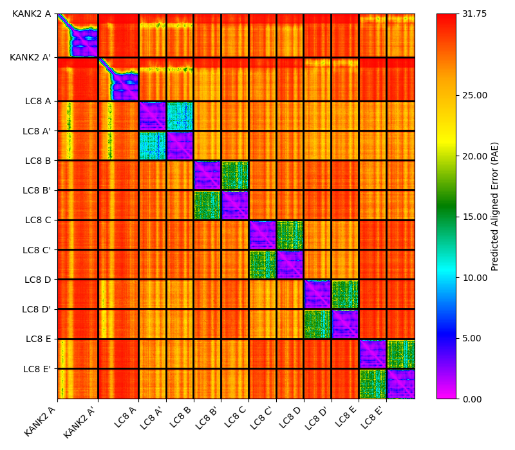

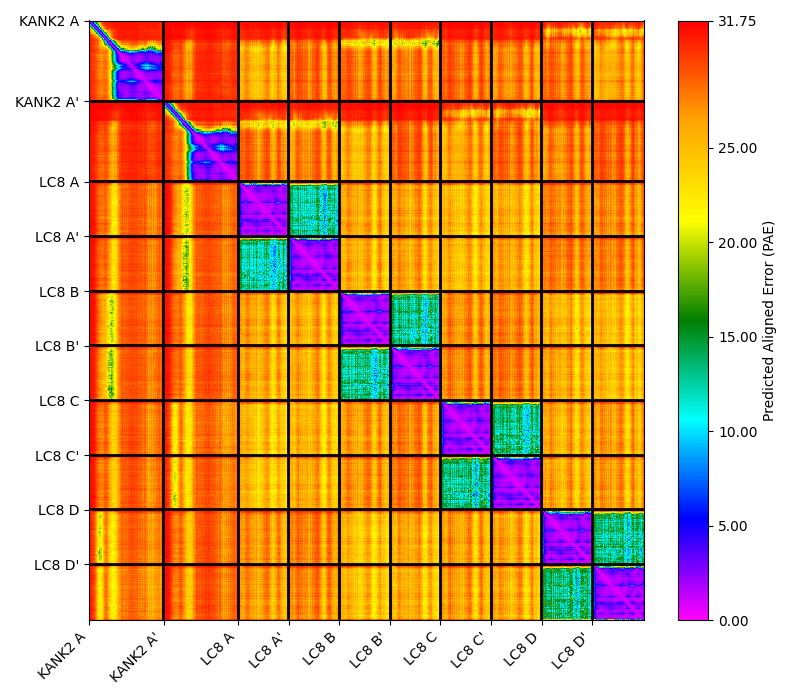

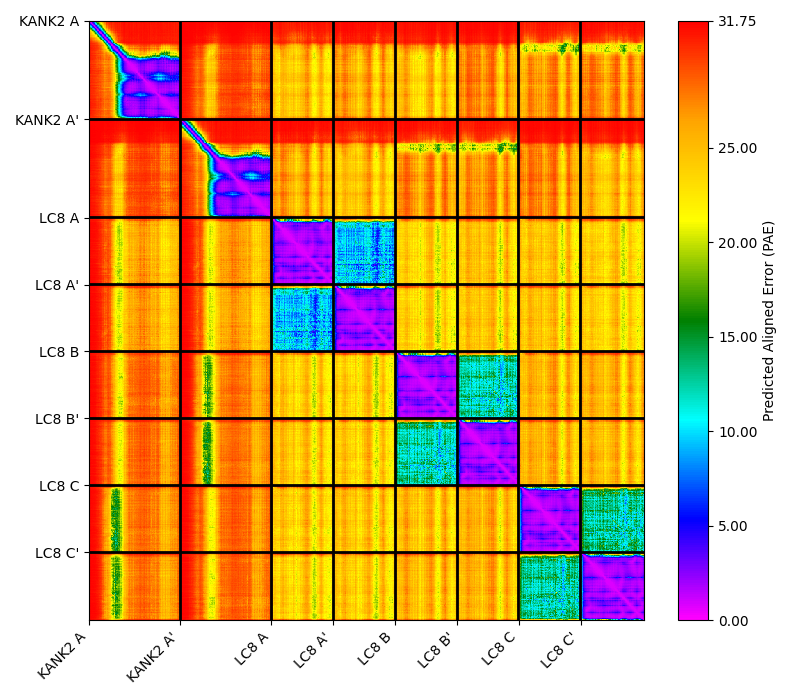

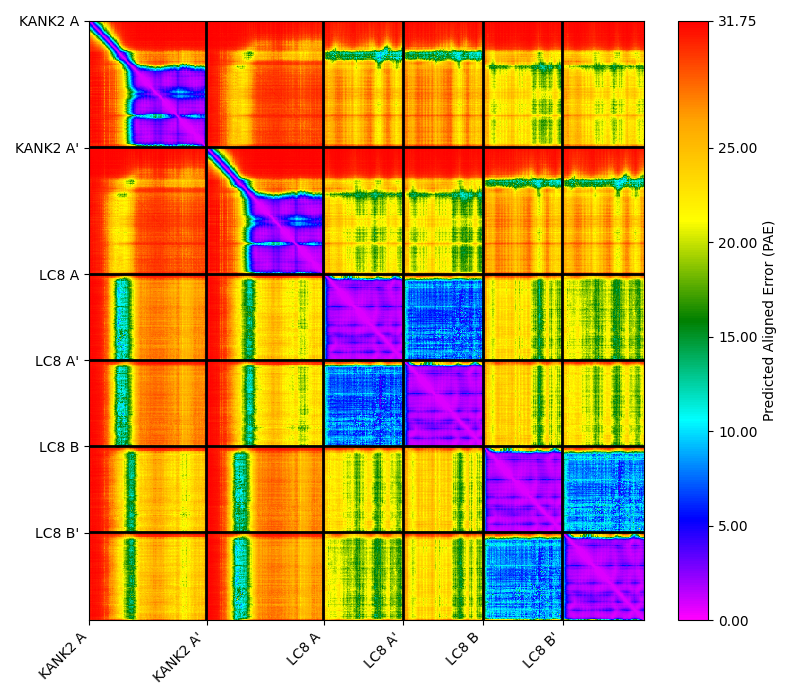

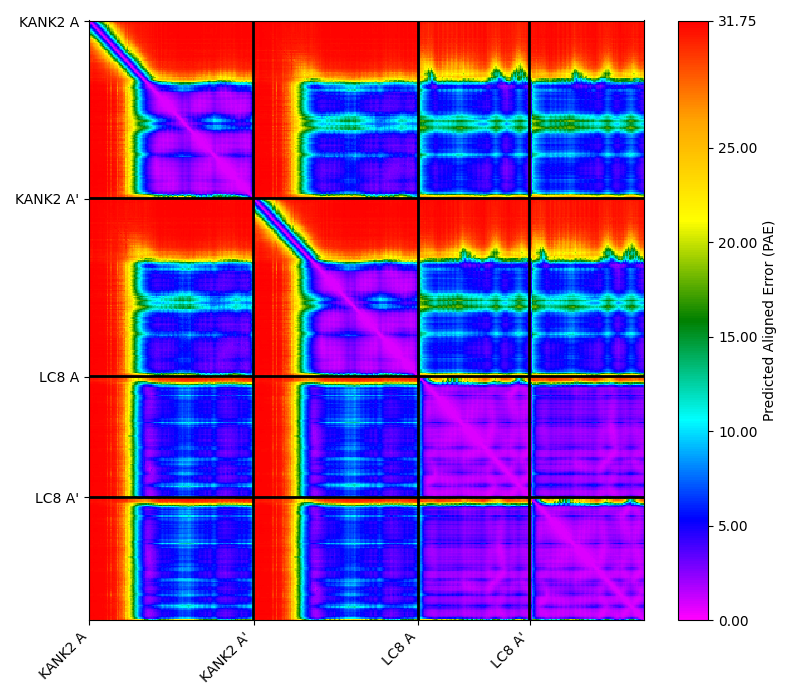

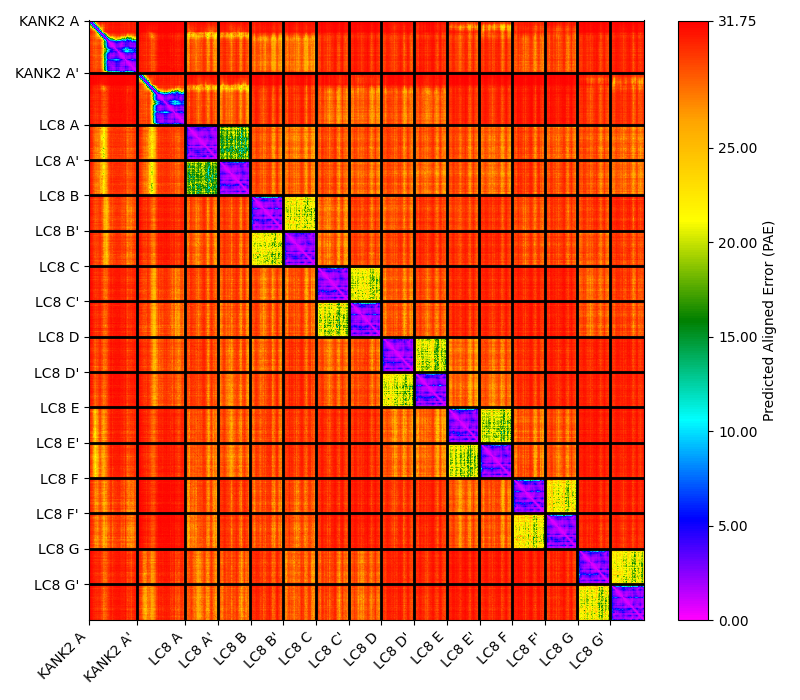

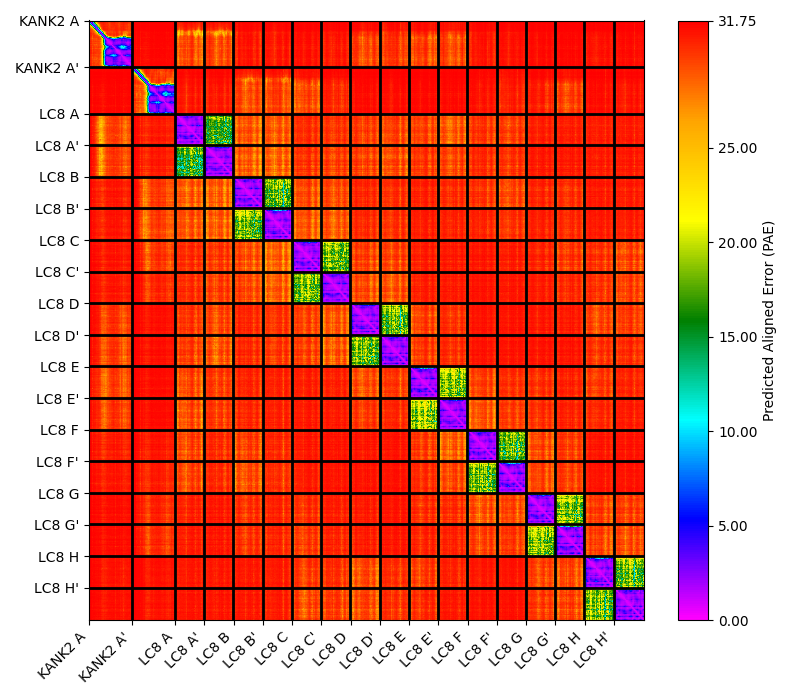

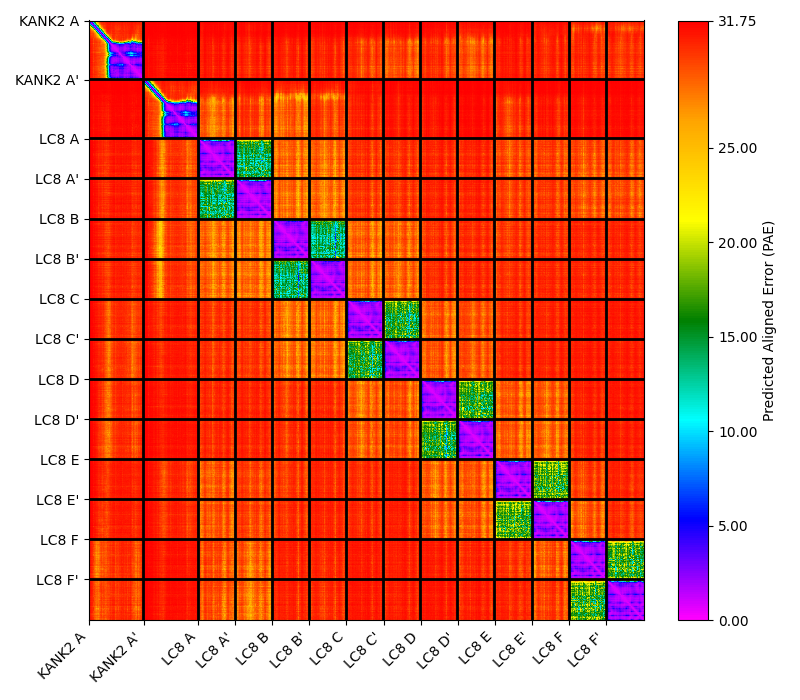

Fig. S13. PAE heatmaps of titration points in the AlphaFoldMultimer titration of LC8 with KANK1_1030-1161_. The C-terminal intramolecular region of confidence for this KANK1 construct is consistent with the presence of the Ankr0 domain. Lack of confidence in KANK1/LC8 interactions is consistent with experimental results.
